# Embryo-scale Visual Cell Sorting reveals a conserved transcriptomic signature of nucleolar size linked to proteostasis

**DOI:** 10.64898/2026.05.25.727721

**Authors:** Hyeon-Jin Kim, Sriram Pendyala, Josephine Lin, Tony Cooke, Gang Li, William Stafford Noble, Xinxian Deng, Christine M. Disteche, Cole Trapnell, Douglas M. Fowler

## Abstract

Nucleoli, nuclear speckles and other compartments regulate transcription, RNA processing, and chromatin organization within the nucleus, yet the relationship of their morphology to developmental gene expression programs *in vivo* is poorly understood. Here, we develop a high-throughput Visual Cell Sorting (VCS) workflow for fixed cells and nuclei that combines antibody-based photoconversion; GPU-accelerated, real-time image analysis; and three-level single-cell combinatorial indexing RNA-seq (sci-RNA-seq3) to link nuclear compartment morphology to single-nucleus transcriptomes at embryo scale. We use VCS to analyze and sort over 1 million mouse embryo-derived nuclei by nucleolar, nuclear speckle, or nuclear size and construct a transcriptional atlas annotated with nuclear compartment phenotypes. Nuclear compartment size varies both between and within lineages and is shaped by proliferation and differentiation. In extracellular matrix protein-producing cell types such as fibroblasts, chondrocytes, and osteoblasts, nucleolar enlargement is uncoupled from cell cycle, and in erythroid cells exhibit a sharp nucleolar contraction preceding cell-cycle exit. We identify a 41-gene transcriptional signature whose expression tracks nucleolar size, enriched for ribosome biogenesis, mitochondrial metabolism, unfolded protein response, stress granule, and ubiquitin–proteasome pathway components. We used this nucleolar transcriptional signature to annotate mouse, zebrafish and human developmental atlases with nucleolar size information, revealing a conserved coupling between nucleolar activity and proteostasis programs. Our work establishes Visual Cell Sorting as a scalable platform for mapping image-based phenotypes to molecular programs; details the relationship between nuclear compartment phenotypes and development; and provides a transcriptional signature to estimate nucleolar size from existing single-cell datasets.

## INTRODUCTION

Embryonic development involves coordinated changes in nuclear organization to execute gene expression programs that specify diverse cell fates. The nucleus is organized into functionally distinct compartments such as nucleoli and nuclear speckles that comprise membraneless, phase-separated condensates of proteins and RNAs^1–3^. These compartments are thought to occupy a large portion of the nuclear space, influencing cellular states by forming chromatin domains^4–8^, regulating transcription^9–11^, and processing mRNAs^12,13^.

Nucleoli and nuclear speckles are among the most well-characterized nuclear compartments. The nucleolus regulates diverse fundamental processes, including ribosome biogenesis^9,14^, the cell cycle^15,16^, stress responses^17,18^, and formation of chromatin domains^4,19–21^. Nuclear speckles, in contrast, serve as a reservoir for splicing factors and transcriptional regulators^12,22,23^, with the spatial proximity of genes to speckles dictating splicing efficiency^13^. While these compartments have well-defined molecular functions, nucleolar and nuclear speckle morphology varies across cell types and differentiation in *in vitro* systems^22,24,25^, suggesting that the morphology of these compartments may reflect changes in their function and in cell state. Moreover, their dynamics in the context of embryonic development remain poorly understood. Thus, investigating how the morphology of these compartments change during development could provide insight into how they contribute to the functional diversity of cell types during embryogenesis.

Recent advances in imaging-based and spatial transcriptomic methods can link morphology with transcriptional states. However, these approaches have several limitations. Probe-based imaging methods^26–31^ enable high-resolution visual phenotyping but are constrained by their reliance on pre-defined probe panels. Spatial transcriptomic approaches^32–42^, on the other hand, provide unbiased, genome-wide transcription profiles within native spatial contexts, yet they often lack the resolution to probe subcellular structures and often do not offer true single-cell resolution. Additionally, many of these methods are not readily scalable for high-throughput, quantitative studies of nuclear organization across large cell populations or whole-organism development, which can span tens to hundreds of millions of cells.

To enable subcellular visual phenotyping and single-nucleus transcriptional profiling at the scale of organismal development, we improved Visual Cell Sorting (VCS), an automated imaging workflow we developed that enables binning and physical separation of live cells based on pre-defined image-based phenotypes^43^. By adapting the workflow for cell fixation and single-nucleus combinatorial indexing-based RNA-seq3^44,45^ (sci-RNA-seq3), we vastly increased throughput and enabled the analysis of diverse biological samples. We used improved VCS to sort over 1 million E15 B6xCAST F1 mouse embryo-derived nuclei based on nucleolar or nuclear speckle size and then quantified the transcriptomes of the sorted nuclei. Our transcriptome-nuclear morphology data reveal that nuclear compartment size varies both between and within cell types. While nuclear compartment size generally scales with cell proliferation and inversely with differentiation, nucleolar size in hepatocytes, fibroblasts, chondrocytes, and osteoblasts instead appears to be driven primarily by the high translational requirements of these cells. Additionally, we identify a set of genes, including genes involved in ribosomal biogenesis and proteostasis, that are tightly associated with the size of the nucleolus relative to the nucleus. We show that these genes can be used to annotate mouse, human and zebrafish single-cell RNA-seq developmental atlases with nucleolar size. These transcriptome-morphology annotations revealed that nucleolar size is highly correlated with proteostasis-related gene expression programs during embryogenesis in mouse, zebrafish and humans.

## RESULTS

### Improved Visual Cell Sorting scales to millions of fixed nuclei for morphologically-defined transcriptomics

We sought to develop a method capable of image-based phenotyping and transcriptome profiling at a scale that can capture the complexity of developing mouse embryos. To achieve this goal, we built upon our previously established Visual Cell Sorting (VCS) workflow, leveraging its high-throughput capabilities^43^. In the original workflow, live cells are imaged, and cells exhibiting the desired image-based phenotype are identified and selectively targeted for recovery through photoconversion of a transgenically expressed photoconvertible protein, Dendra2^46^. Selective targeting is accomplished using a digital micromirror device, which can illuminate a subset of cells based on their phenotype, photoconverting Dendra2 and marking the cells. Following imaging and photoconversion of the entire population of cells, photoconverted populations are then lifted off of the slide or plate and sorted by the degree of Dendra2 photoconversion using fluorescence-activated cell sorting (FACS).

Applying Visual Cell Sorting to embryo-derived nuclei required a series of technical innovations. First, nuclei from embryos must be fixed to the surface of a well for imaging and photoconversion, and then must be dissociated to form a single cell or nucleus suspension for sorting. We developed a method to dissociate cells or nuclei fixed to wells for imaging by gently scraping the surface with a micropipette tip (**Figure 1A**). We tested the ability of our method to yield single nuclei suspensions by co-culturing U2OS cells expressing GFP- or miRFP-tagged histone 3, fixing them to the well with 4% paraformaldehyde (PFA), and then scraping the well to suspend them in sucrose nuclei buffer. Flow cytometry of the suspension showed a 1% GFP+/miRFP+ doublet rate, with the remaining nuclei having either GFP or miRFP but not both, indicating effective dissociation to single nuclei (**Figure 1B**). We confirmed the purity of these sorted nuclei by imaging (**Supplementary Figure 1A,B**). We further validated our dissociation strategy by applying VCS to separate a population of fixed U2OS cells each of which expressed either wild type lamin A, which forms a diffuse pattern in the nucleus, or N195K, a lamin A variant known to aggregate. Imaging of the sorted nuclei following photoconversion revealed high phenotypic purity, with 88.5% and 83% of WT and punctate nuclei correctly assigned, respectively (**Supplementary Figure 1C,D**). Thus, PFA fixation does not alter the ability to isolate visually distinct populations with VCS.

**Figure 1.**
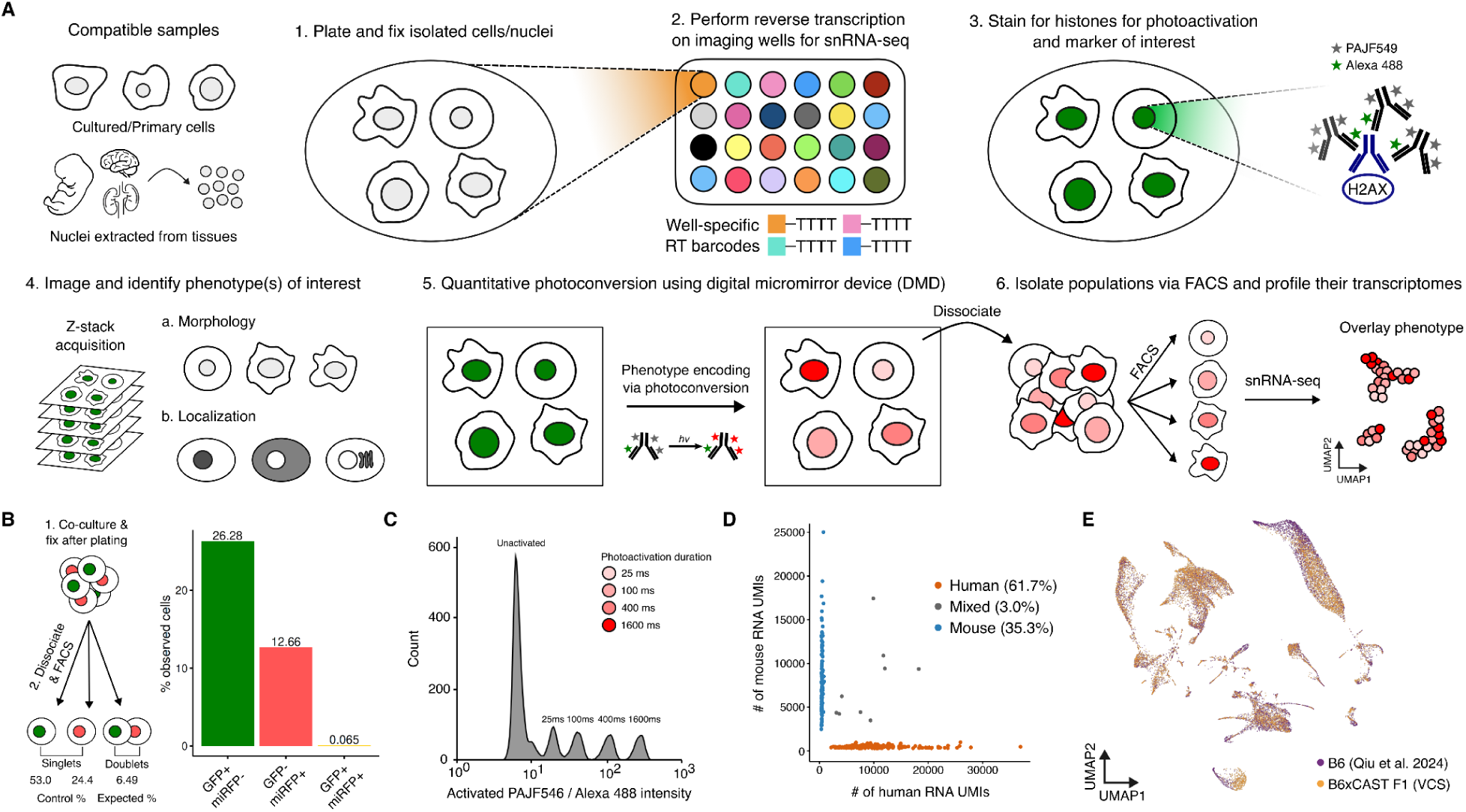
Overview of the improved Visual Cell Sorting (VCS) workflow. (A) Workflow of the VCS method, which is compatible with cultured cell lines and nuclei extracted from frozen or fresh tissues. Following sample plating and fixation with 4% PFA, *in situ* reverse transcription is performed directly within the imaging wells using well-specific barcoded RNA primers. Samples are then immunostained for target proteins and histones using photoconvertible antibodies, which serve to physically encode the visual phenotypes. After imaging and identifying visual phenotypes of interest, a digital micromirror device is used to direct the microscope to selectively illuminate cells/nuclei with the desired visual phenotypes. Labeled populations are then physically isolated via FACS for downstream sci-RNA-seq3 profiling. (B) Schematic and corresponding barplot demonstrating the integrity of the dissociation method by performing a GFP+/miRFP+ co-culture experiment, with an observed doublet rate of 1%. (C) Flow cytometry plot demonstrating that photoconversion can be tuned to label four visually distinct populations. (D) Human/mouse species mixture experiment demonstrating low collision rates (3.0%) for the optimized VCS-sci-RNA-seq3 protocol. (E) UMAP embedding of E15 mouse embryo nuclei, showing high concordance between the conventional sci-RNA-seq3 (purple) and VCS-sci-RNA-seq3 pipeline (gold).

Previously, VCS relied on the expression of a photoconvertible fluorescent protein, Dendra2, in live cells. However, many samples of interest, including mouse embryos, cannot easily be engineered to express Dendra2. Leveraging our success with imaging and resuspending fixed nuclei, we developed a method for using cell-staining dyes and fluorescent antibodies with VCS. For phenotype encoding, we replaced Dendra2 with an anti-histone H2AX antibody co-labeled with a photoconvertible dye, PAJF549^47^, and labeling control dye, Alexa 488 (**Figure 1A third panel**). This combination of dyes mimics the optical properties of Dendra2, enabling us to encode up to four visually distinct populations using different degrees of PAJF549 photoconversion (**Figure 1C** and **Supplementary Figure 1E**).

In the original version of Visual Cell Sorting, live cells were sorted and then processed using a commercial 10X single cell RNA sequencing platform. However, RNA molecules are prone to rapid degradation upon cell lysis and even after cell fixation. Therefore, to enable single nucleus transcriptome profiling of visually sorted nuclei, we performed reverse transcription on fixed cells or nuclei prior to antibody staining, imaging and sorting. A species mixture experiment with our optimized protocol revealed minimal doublet populations (**Figure 1D**; 3.0% observed), with RNA yield comparable to that of the standard sci-RNA-seq3 protocol. We also conducted pilot experiments using mouse neural progenitor cells and nuclei derived from frozen mouse embryos, and detected an average of 2,831 and 1,054 UMIs per nucleus, respectively (**Supplementary Figure 2A**).

Finally, the previous implementation of VCS was limited to 250,000 cells per day, partially because real-time image processing and cell photoconversion was restricted to the microscope software. To overcome this bottleneck, we wrote custom scripts to bridge the microscope to an external GPU for accelerated image analysis, thereby boosting throughput to ∼1 million cells per day (**Supplementary Figure 1F**). Concurrently, we fitted the microscope with a Piezo Z-stage for multi-plane z-stack acquisition, enabling visual phenotyping and single nucleus transcriptome profiling at a scale that captures the complexity of developing mouse embryos. To evaluate our optimized workflow, we performed a pilot experiment where nuclei derived from E15 B6xCAST F1 embryos were randomly photoconverted, sorted and subject to sci-RNA-seq3. Co-embedding of the VCS-derived transcriptome data with the published mouse embryogenesis single-cell atlas^48^ showed minimal batch effects (**Figure 1E**). Importantly, when compared to the reference atlas, the VCS data recovered expected cell type proportions, showed high concordance at the pseudobulk level, and preserved lineage-specific marker expression (**Supplementary Figure 2B-G**). However, we noted minor exceptions within the erythroid and hepatocyte populations. In the erythroid lineage, late-stage cells were underrepresented in our data because the minimum nuclear size filter likely excluded cells undergoing enucleation (**Supplementary Figure 3A,B**). In contrast, differences in hepatocyte profiles likely arise from allele-specific expression between the B6 and CAST genomes^49,50^ and enhanced growth due to hybrid vigor^51^ in B6xCAST hybrids, characterized by increased expression of metabolic (*Sik3, Mcu*) and growth regulators (*Yap1, Trim71*) (**Supplementary Figure 3C-G**).

### Visual Cell Sorting of embryo-derived nuclei by nucleolar and nuclear speckle size

Given the critical roles of nucleoli and nuclear speckles in ribosomal biogenesis and mRNA splicing, we applied Visual Cell Sorting to sort nuclei derived from the E15 B6xCAST F1 mouse embryos using stains that reveal nucleoli (nucleolin, Ncl) or speckles (Srrm2, SC35) (**Supplementary Figure 4** and **Supplementary Tables 1,2**). In order to identify regions that correspond to nuclear compartments, we trained a ResNet-34-based U-Net segmentation model (**Figure 2A** and Methods). Compared to manual annotations, our model achieved high performance for nucleoli (median Jaccard index: 0.80, median Dice coefficient: 0.89) and moderate performance for nuclear speckles (median Jaccard index: 0.49, median Dice coefficient: 0.66; **Supplementary Figure 5**). Following imaging, the mouse embryo nuclei were then photolabeled according to their nuclear phenotype using a quartile binning strategy (**Figure 2B,C**). Here, we refer to these bins as Q1, Q2, Q3, and Q4, where Q1 and Q4 indicate the nuclei from the bottom and top 25% of the distribution of the measured nuclear phenotype, respectively.

**Figure 2.**
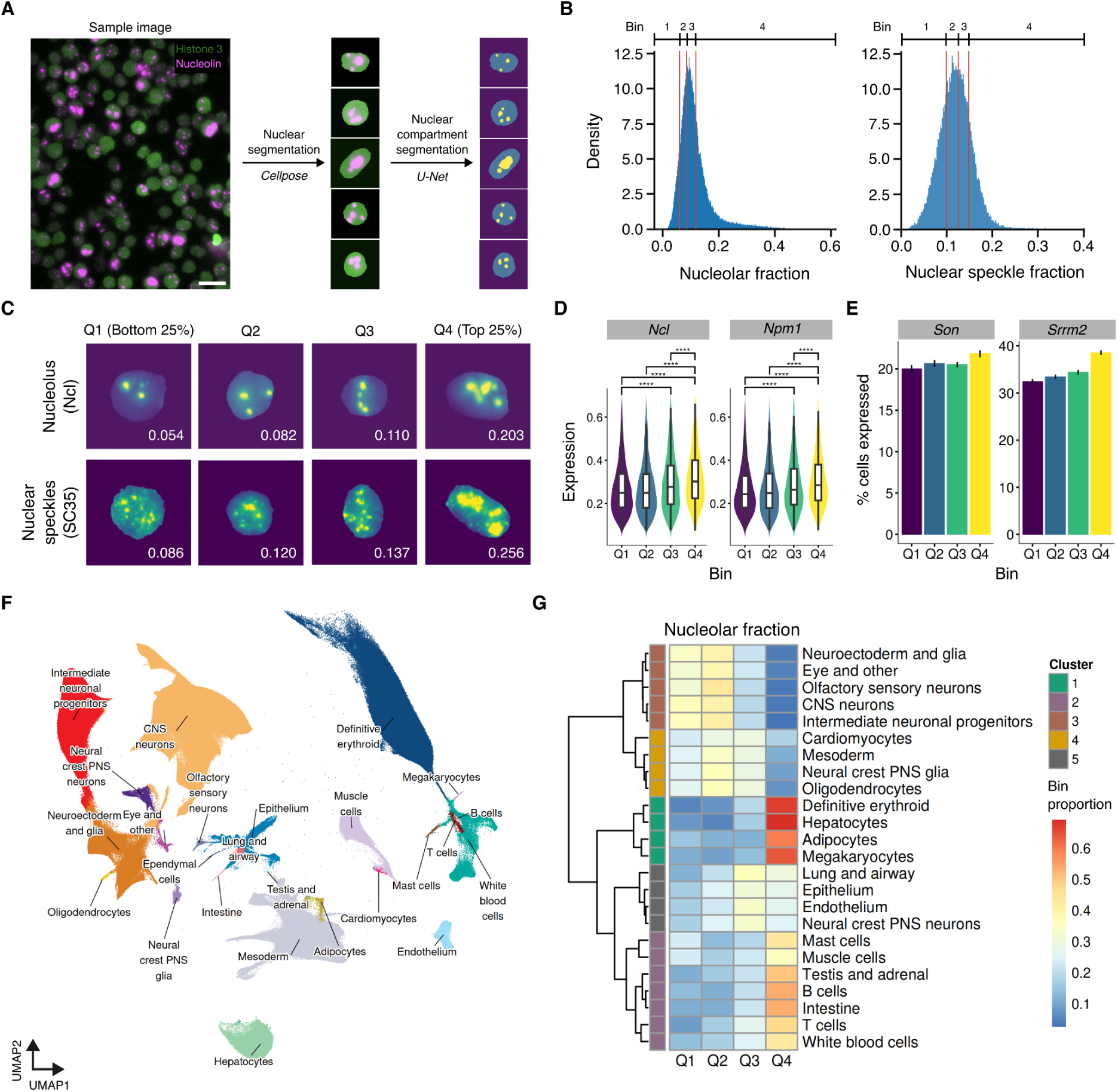
Visual Cell Sorting and snRNA-seq profiling of E15 mouse embryo derived nuclei. (A) Overview of image analysis workflow. Nuclei were stained for histone H2AX for photoconversion and either nucleolin or SC35. Following nuclear segmentation via Cellpose^106^, nuclear compartment areas were computed via segmentation with an in-house trained U-Net. Scale bar = 5 µm. (B) Density plots illustrating the nucleolar (left) and nuclear speckle (right) fractions of pre-imaged E15 mouse embryo nuclei, binned into equal quartiles (red lines). (C) Example crops for nucleoli (top) and speckles (bottom) across the four quartiles, with the relative fraction values displayed in each crop. (D) Distribution of nucleolar marker gene expression (*Ncl, Npm1*) across the four sorted quartiles. Significance levels: **** *p* < 0.00001. (E) Barplots indicating the percentage of cells within each speckle size bin expressing at least one UMI for *Son* and *Srrm2*. (F) UMAP embedding of visually sorted nuclei profiled in this experiment, colored by the main trajectory group. (G) Heatmap showing the distribution of nucleolar fraction bin labels across the major E15 trajectory groups.

We profiled the visually sorted nuclei with sci-RNA-seq3 to build a comprehensive mouse organogenesis atlas annotated with nuclear and sub-nuclear size phenotypes. For each marker, nuclei were sorted based on either absolute size (total segmented pixels) or fraction (proportion of nuclear area occupied by the compartment). In total, we profiled 1,156,362 nuclei (median UMI count of 1,049), with 547,533 nuclei (n = 13 embryos) phenotyped for nucleolar size, 310,355 nuclei (n = 4 embryos) phenotyped for nuclear speckle size, and 298,474 nuclei (n = 5 embryos) phenotyped for nuclear size as a control (**Supplementary Figure 6A-C**). Our data exhibited no apparent batch effects and recovered uniform lineage proportions with high correlation across embryo replicates (**Supplementary Figure 6D,E**). To annotate cell types, we co-embedded our data with the reference mouse embryogenesis atlas^48^ using a nearest-neighbor algorithm^35^ followed by manual refinement (**Figure 2F** and Methods). Reassuringly, our data showed high-concordance with atlas-defined lineage frequencies and marker gene expression patterns (**Supplementary Figure 6F,G**).

We next assessed whether the VCS bins and expression of markers used for visual phenotyping were correlated. As expected, nucleolar size quartile bin labels were positively correlated with the expression of main nucleolar components, such as *Ncl* and *Npm1* (**Figure 2D** and **Supplementary Figure 7A**). The SC35 antibody interacts with several splicing factors that localize in nuclear speckles, such as Srrm2, Srsf2, and Son^52^. Surprisingly, SC35 size bins did not significantly correlate with total expression of these marker proteins, but rather with the percentage of nuclei showing detectable expression of the markers (at least one UMI; **Figure 2E** and **Supplementary Figure 7B-D**). This observation suggests that nuclear speckle size reflects a general, population-wide activation of these genes, as opposed to upregulation within individual cells.

Next, we explored how the nuclear compartment size was distributed in the E15 B6xCAST F1 mouse embryos by grouping the main cell lineages based on the distribution of nuclei across the bins (**Supplementary Figure 7E-H**). The 24 annotated main cell lineages with at least 100 nuclei in our atlas were unevenly distributed with respect to nucleolar fraction, forming five distinct clusters (**Figure 2G**). For example, over 60% of nuclei from the erythroid, hepatocyte, adipocyte, and megakaryocyte lineages as well as more than 50% of nuclei from adrenocortical, pancreatic acinar, and gut epithelial cells were found in the bin corresponding to the highest nucleolar fraction (Q4; **Supplementary Figure 7H**). Conversely, postmitotic neuronal and glial lineages were highly concentrated within the smallest nucleolar areas, with 70% of these nuclei found in the bin containing the smallest nucleoli (Q1). The distribution of cells with respect to nuclear speckle fraction was similar but less pronounced, with immune cell lineages showing less enrichment in the higher size bins (Q3 and Q4) relative to their nucleolar fraction (**Supplementary Figure 7G**).

### Cell proliferation and differentiation partially account for variability in nucleolar and nuclear speckle size within lineages

Nuclear compartment size depends on the type, state and function of the cell^1,53,54^. To assess the differences in nuclear compartment size, we clustered nuclei based on their transcriptomes and then transformed the categorical VCS bin labels for all cells in a cluster into a weighted average phenotype score (**Figure 3A, Supplementary Figure 8A** and Methods). Our approach assigned a quantitative phenotype score to every cluster reflecting the measured nuclear compartment size, operating under the assumption that transcriptionally similar cells are also morphologically similar. By converting the bin labels to scores, we were able to quantitatively evaluate the differences in nucleolar and nuclear speckle size between and within cell lineages (**Figure 3B** and **Supplementary Figure 8B-D**). Although nucleolar and nuclear speckle size are correlated, our approach revealed clear differences in their distribution across certain lineages, reflecting their distinct functional roles within the nucleus (**Figure 3C** and **Supplementary Figure 8E**).

**Figure 3.**
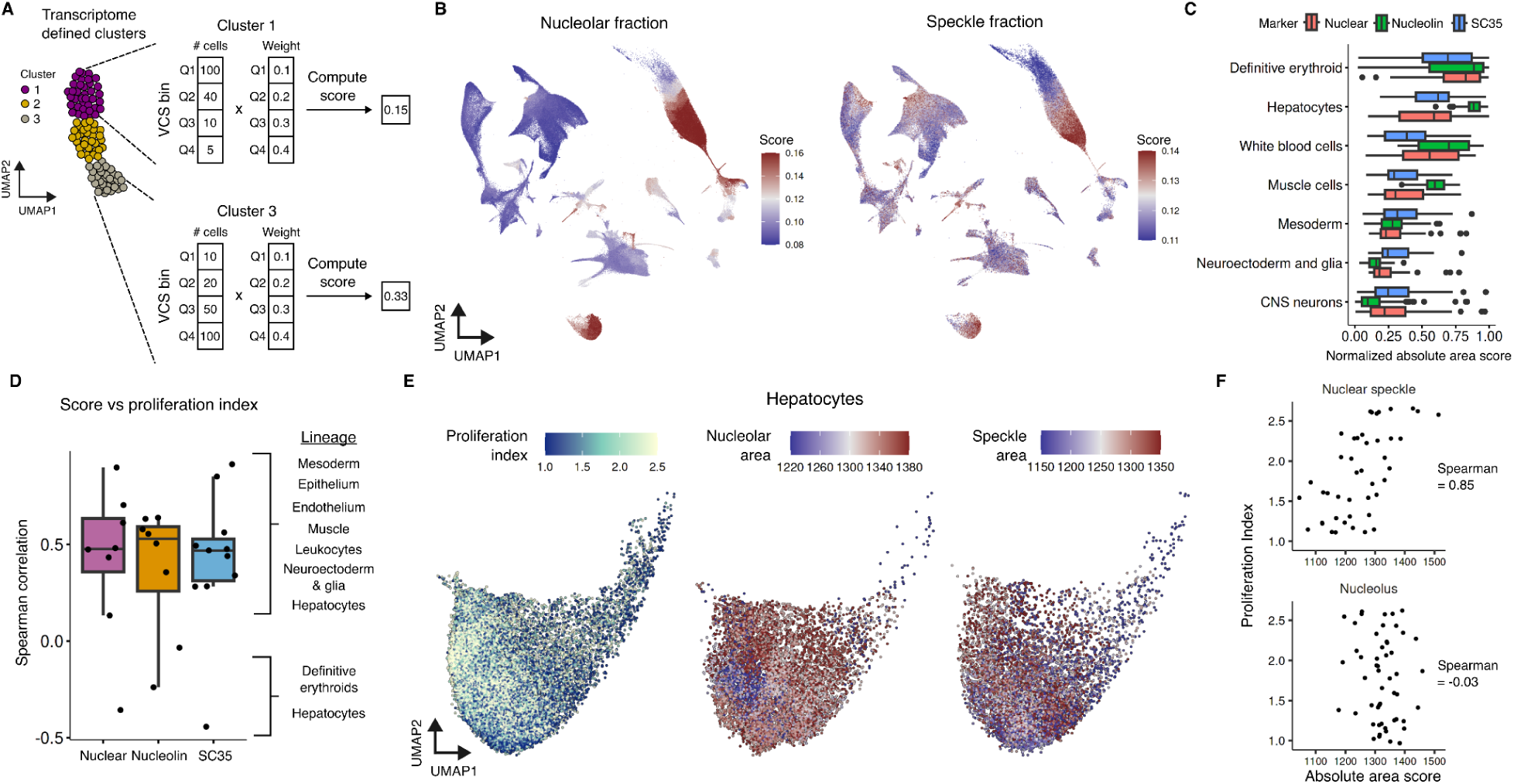
Correlation between cluster-level nuclear compartment scores and proliferation. (A) Overview of the cluster-level weighted average nuclear compartment score calculation. Clusters are assigned transcriptionally and the scores are computed using the bin counts and weights obtained from the image distributions. (B) UMAP embeddings colored by the nucleolar fraction (left) and nuclear speckle fraction scores (right). (C) Boxplot comparing the absolute size distributions of nuclei, nucleoli, and nuclear speckles across selected lineages. (D) Boxplots summarizing the repeated measures rank correlation between nuclear compartment size scores and the proliferation index across cell lineages. (E) UMAP embeddings of the hepatocyte lineage colored by the proliferation index (left), absolute nucleolar area (middle), and absolute nuclear speckle area (right). (F) Scatterplots showing the relationship between the proliferation index and the absolute compartment areas in hepatocytes.

Using these quantitative scores, we sought to identify the sources of variability in nucleolar and nuclear speckle size within the embryo. We hypothesized that, because cells must double their total genetic and cellular content to divide^55,56^, the cell cycle would be a major driver of nuclear compartment size. To assess the relationship between the cell cycle and absolute nuclear compartment sizes measured by VCS, we compared the size scores with a proliferation index^57^ computed from the transcriptome of each cell. Across well-represented lineages (≥ 10 clusters), nuclear, nucleolar and nuclear speckle size correlated with proliferation index, with 7 of 9 lineages following the expected positive relationship (**Figure 3D** and **Supplementary Figure 9**). However, two lineages deviated from this trend. Definitive erythroid cells exhibited a general negative correlation across all three size markers. In hepatocytes, this trend was isolated to the nuclear and speckle size, which exhibited positive correlation with proliferation, whereas the nucleolar area remained elevated at a high baseline regardless of proliferative state (**Figure 3E,F**). This suggests that the hepatocyte nucleolus size primarily reflects metabolic and secretory demands, supporting the continuous production of bile, albumin, and other secretory proteins^58^.

In addition to proliferation, differentiation drives changes in nuclear compartment size^59–61^. Trajectory analysis of ten well-resolved sub-lineages revealed that nuclear compartment size correlated negatively with pseudotime in six of these lineages (**Figure 4A** and **Supplementary Figures 10-14**), consistent with the reductions previously observed during *in vitro* differentiation of mouse embryonic stem cell into neural progenitor cells^24^. For example, as maturing erythroid cells approached cell cycle exit, both nucleolar and speckle fractions decreased by 66%, corresponding to a 52% and 44% drop in their absolute areas, respectively (**Figure 4B,C** and **Supplementary Figure 15A,B**). Uniquely, in erythroid cells, the effects of differentiation on nuclear compartment size dominated those of the cell cycle; even as the proliferation index continued to increase during differentiation, compartment sizes steadily decreased, preceding eventual cell cycle exit. Furthermore, the nucleolar fraction score tightly correlated with the expression of the key hemoglobin-regulating factors, *Bcl11a* and *Lin28b* (R = 0.998 and 0.994), reflecting Lin28b’s role as a ribosome-associated protein that suppresses *Bcl11a1* translation while both mRNAs are concurrently expressed during early maturation^62^ (**Figure 4D** and **Supplementary Figure 15C,D**). These results may suggest that erythroid progenitors rapidly scale up ribosome biogenesis to fuel an early proliferation burst, tapering only after reaching a threshold sufficient to sustain continued expansion and eventual terminal maturation.

**Figure 4.**
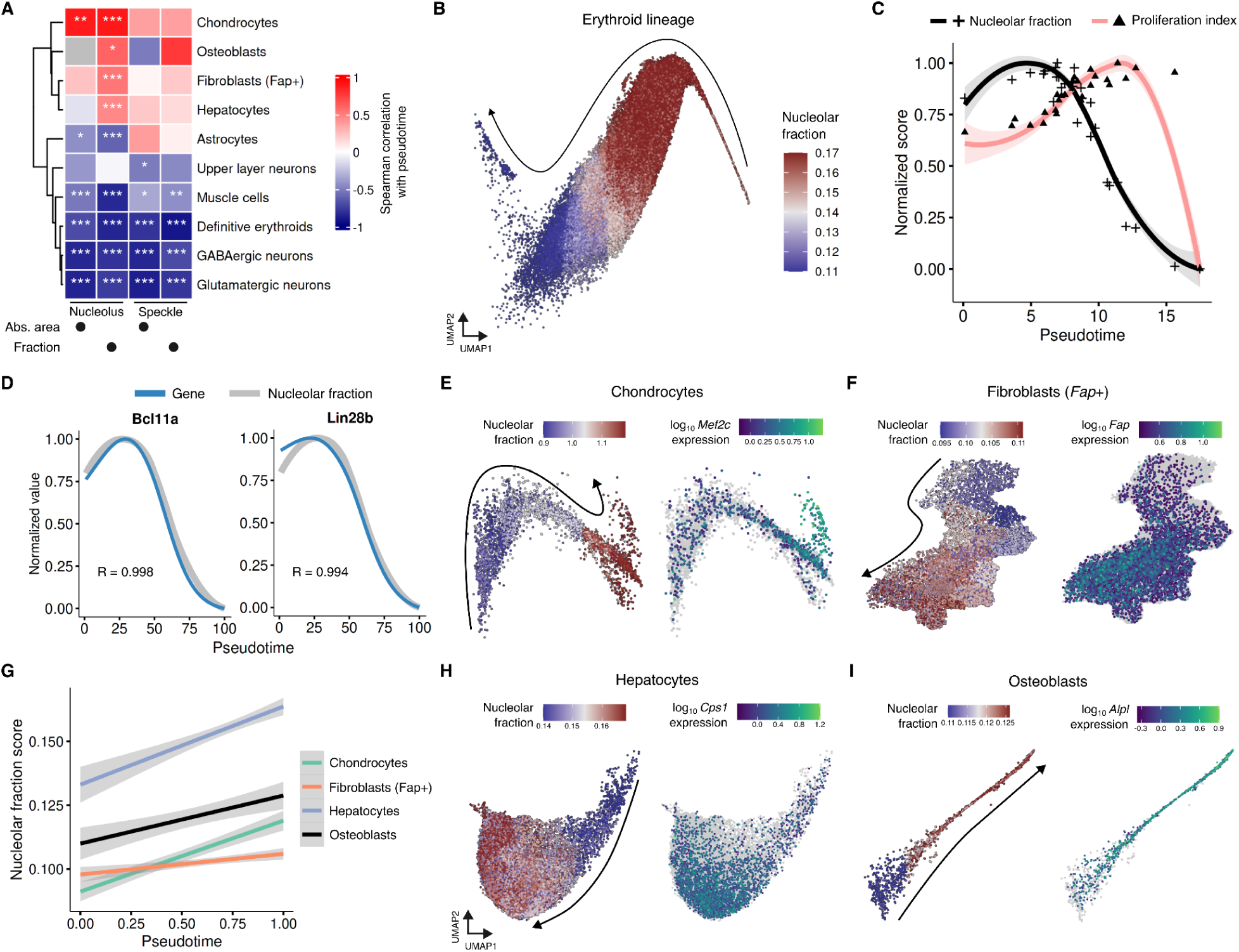
Dynamics of nuclear compartment scores during cellular differentiation. (A) Heatmap displaying repeated measures rank correlations between absolute and fraction nuclear compartment scores and pseudotime across E15 cell lineages. Significance levels: * *p* < 0.05, ** *p* < 0.01, *** *p* < 0.001. (B) UMAP embedding of the definitive erythroid lineage, colored by the nucleolar fraction score. (C) Pseudotemporal changes in the nucleolar fraction score (black crosses) and the proliferation index (pink triangles) during erythroid differentiation. Curves are fitted with a cubic spline function. (D) Expression profiles of *Bcl11a* (left) and *Lin28b* (right) overlaid against the nucleolar fraction score (grey). (E,F) UMAP embeddings of the chondrocyte (E) and fibroblast (F) lineages colored by the nucleolar fraction score (left) and the marker expression (*Mef2c* and *Fap*, respectively; right). (G) Lineplots demonstrating a positive correlation between pseudotime and the nucleolar fraction score for chondrocytes (green), fibroblasts (orange), hepatocytes (blue), and osteoblasts (black). Shaded regions indicate confidence intervals. (H,I) UMAP embeddings of the hepatocyte (H) and osteoblast (I) lineages colored by the nucleolar fraction score (left) and the marker expression (*Cps1* and *Alpl*, respectively; right). Arrows indicate the progression of differentiation.

In contrast, chondrocytes, osteoblasts, fibroblasts, and hepatocytes exhibited an increase in nucleolar size during differentiation (**Figure 4E-I** and **Supplementary Figure 14**). These cell types produce large amounts of proteins, including extracellular matrix proteins for endochondral ossification^63,64^ and wound healing^65,66^, suggesting enhanced nucleolar activity is necessary to support the increased demands for protein synthesis. Altogether, these results demonstrate that while proliferation and differentiation status are primary drivers of nuclear compartment size, functional demands play a key role in further determining the final size in some lineages.

### Shared transcriptional signatures associate with nucleolar fraction across lineages

Next, we sought to identify transcriptional signatures comprising genes whose expression levels were correlated with the size of nucleoli or speckles across cells. We focused on the nucleolar and speckle fraction phenotypes to account for differences in the absolute size of each compartment driven by variations in nuclear size. Using these fraction scores, we computed repeated measures rank correlations^67^ against each gene’s expression within individual lineages. We then combined these intra-lineage metrics and filtered for genes demonstrating strong correlations both within individual lineages (intra-lineage correlation ≥ 0.4 in at least six lineages) and globally across all clusters (global correlation ≥ 0.7; Methods). Surprisingly, no genes correlated with nuclear speckle fraction phenotype met these filtering criteria. This result is likely explained by the tight correlation between absolute nuclear speckle size and nuclear size; using the fraction score, which is normalized for nuclear area, removes the primary source of variation (**Supplementary Figure 16**).

However, we identified 41 upregulated genes whose expression was strongly associated with the nucleolar fraction score (**Figure 5A, Supplementary Figures 17-18** and **Supplementary Tables 3,4**). Overall, a higher nucleolar fraction score correlated with increased expression of genes driving rRNA transcription and ribosome biogenesis (e.g. *Nop58, Wdr43, Taf1d*), along with translational machinery (e.g. *Gspt1, Larp4, Gemin5*) and mitochondrial function (e.g.

**Figure 5.**
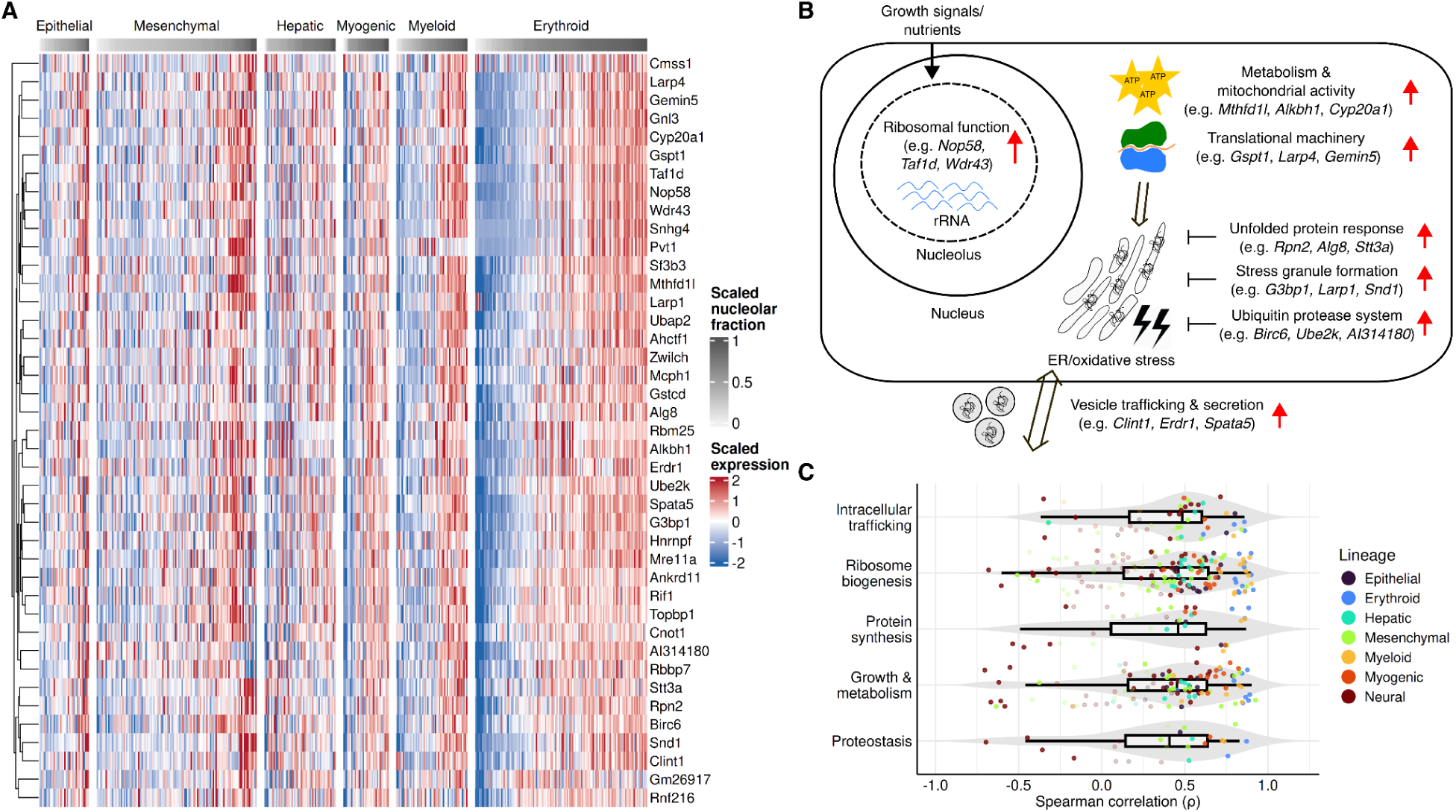
Transcriptional signatures associated with the nucleolar fraction score across and within lineages. (A) Heatmap displaying Z-score normalized expression values of the 41 genes most strongly correlated with the nucleolar fraction score (intra-lineage correlation ≥ 0.4 in at least six lineages and global correlation ≥ 0.7). Columns represent individual clusters, organized by major E15 cell lineages. (B) Graphical representation of biological processes upregulated in association with an increase in the nucleolar fraction score. (C) Distributions of Spearman correlation coefficients evaluating the relationship between the nucleolar fraction score and the aggregate expression of Hallmark^73^, GO:CC^74^ and KEGG^75^ gene sets. These gene sets were grouped by overarching biological function and filtered for significance using Stouffer’s method. Points represent individual cell lineages.

*Mthfd1l, Alkbh1, Cyp20a1*; **Figure 5B**). Genes involved in vesicle trafficking and secretion (e.g. *Clint1, Erdr1, Spata5*) were similarly upregulated. Elevated translational and secretory burden, however, is accompanied by cellular stress^68–70^, including oxidative and proteostatic stress. To mitigate this, protective pathways such as the unfolded protein response (e.g. *Stt3a, Rpn2, Alg8*), stress granule formation (e.g. *G3bp1, Larp1, Snd1*), and the ubiquitin-proteasome system (e.g. *Birc6, Ube2k, AI314180*) were also upregulated. Conversely, genes negatively correlated with the nucleolar fraction score primarily consisted of neuronal adhesion markers (**Supplementary Figure 20A,B** and **Supplementary Table 5**). An interesting exception was *Malat1*, which exhibited negative correlation across all embryonic lineages except in hepatocytes, where it is known to promote proliferation^71,72^ (**Supplementary Figure 20C**).

To further evaluate the association between nucleolar function and these processes, we compared the VCS-derived nucleolar size score against established biological gene sets (MSigDB Hallmark^73^, GO:CC^74^ and KEGG^75^). Employing a filtering strategy similar to our single gene analysis, we assessed gene set correlations across lineages (repeated measures rank correlation > 0.3 in at least seven lineages) and consolidated the top 30 into five functional categories. In agreement with the biological roles we identified for individual genes, these categories encompassed processes such as ribosome biogenesis, metabolism, and proteostasis (**Figure 5C** and **Supplementary Figure 19A**). Over-representation analysis of the 41 strongly correlated genes, complemented by gene set enrichment analysis of the total gene list ranked by their correlation, further supported these associations, with both methods converging on similar groups of genes (**Supplementary Figure 19B,C**). Altogether, these findings suggest that high translational demand is closely coupled with stress response activation, likely serving as a compensatory mechanism to preserve cellular proteostasis.

### A nucleolar size-associated set of genes can estimate nucleolar size across vertebrate development

We identified a nucleolar transcriptional signature composed of 41 upregulated genes that are associated with an increase in nucleolar fraction. We asked whether we could use these genes to estimate nucleolar fraction for a single-cell mouse embryogenesis atlas^48^ comprising ∼11 million nuclear transcriptomes spanning late gastrulation (E8.5) to birth (P0). To do so, we trained a generalized linear mixed model on the E15 VCS data to estimate nucleolar fraction from the aggregate expression of these signature genes. This approach assigned scores to individual cells, allowing comparisons across distinct cellular states (**Figure 6A,B, Supplementary Figure 21A** and Methods). The estimated score for the E15 cell types in the developmental atlas exhibited strong correlation with the score we computed from VCS-derived E15 embryos (R = 0.79; **Figure 6C**). Projection neurons, including cranial motor, dorsal root ganglion, parasympathetic, and retinal ganglion neurons, displayed a higher measured nucleolar fraction than predicted by the gene signature. Generally, our score estimates preserved broad cell-type differences and were highly correlated within most lineages (**Supplementary Figures 18B, 21B-E**).

**Figure 6.**
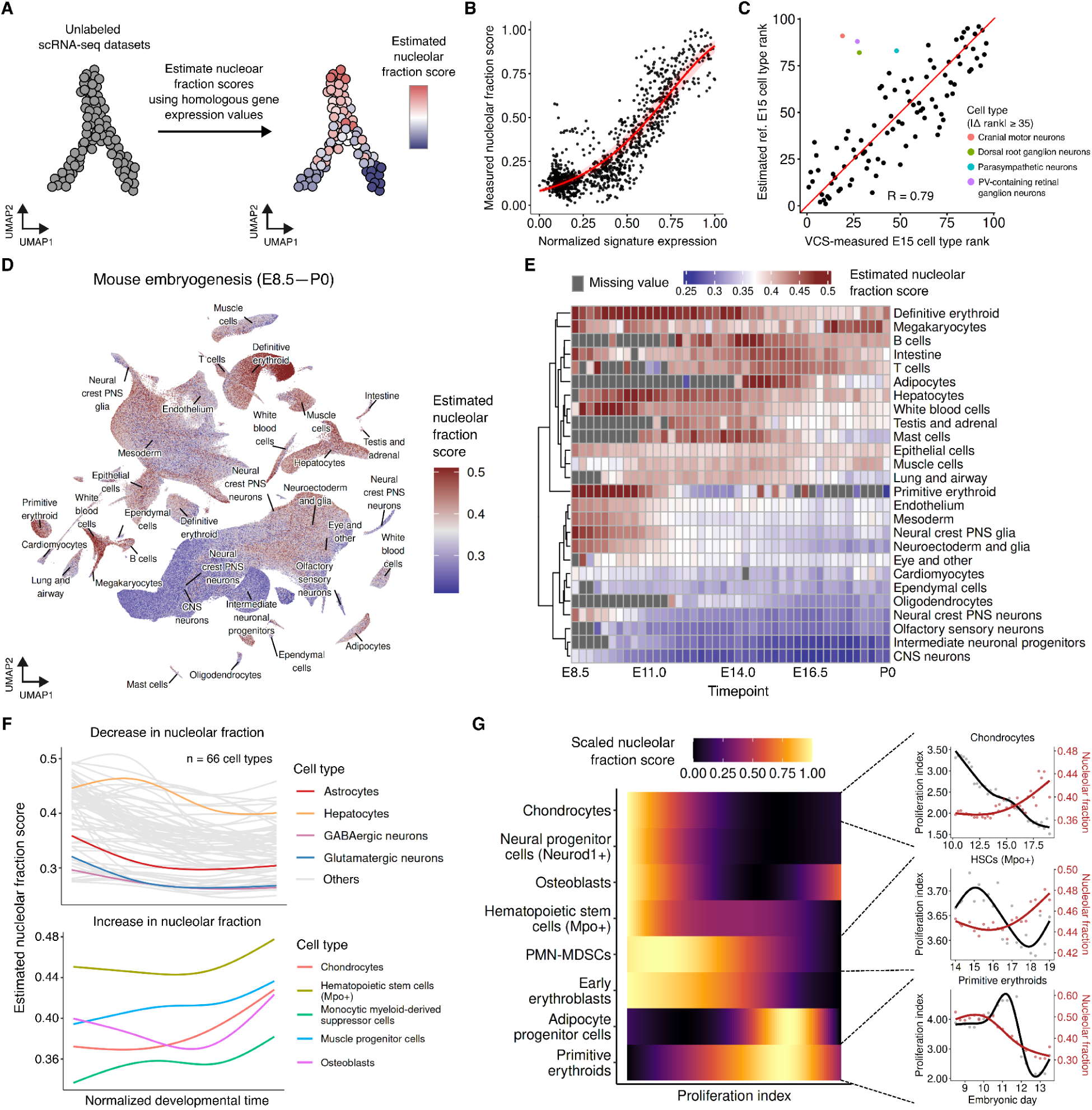
Estimating the nucleolar fraction score across a mouse embryogenesis atlas. (A) Schematic of the nucleolar fraction score estimation for unlabeled single-cell RNA-seq datasets. The min-max normalized aggregate expression of the 41-gene nucleolar fraction signature is converted to a score using the relationship derived from the E15 VCS data. (B) Scatterplot of the normalized signature expression vs the measured nucleolar fraction score across all clusters, overlaid with a generalized linear mixed model fit (red curve). (C) Scatterplot comparing the measured and estimated cell type nucleolar fraction ranks (red line represents y = x). Outliers with an absolute rank difference ≥ 35 are highlighted. (D) UMAP embedding of a mouse embryogenesis atlas^48^, colored by the estimated nucleolar fraction score. (E) Temporal distribution of the median estimated nucleolar fraction score across the main trajectory groups. (F) Lineplots illustrating temporal changes in the estimated nucleolar fraction score for each cell type during embryogenesis. To visualize the changes, the median score for each cell type across the embryonic stage was fitted using a cubic spline function. Among 93 well-represented cell types (defined as having at least 300 cells across 10+ time points), 66 showed a decreasing trend (top) and 5 showed an increasing trend (bottom) across embryonic stages. (G) Heatmap (left) highlighting cell types where the estimated nucleolar fraction score is negatively correlated with the proliferation index. Values were min-max scaled within each cell type. Dual-axis scatterplots (right) showing the changes in proliferation (black) and the estimated nucleolar fraction score (red) for chondrocytes, *Mpo*-positive hematopoietic stem cells, and primitive erythroids.

Next, we investigated how the estimated nucleolar size varies during mouse embryogenesis (**Figure 6D** and **Supplementary Figure 22**). On average, the estimated nucleolar fraction score gradually decreased over embryonic development, likely reflecting progression of differentiation. Cell lineages with similar metabolic demands across time points were grouped together according to the score (**Figure 6E**). For example, several neuronal cell types, which acquire their post-mitotic identity as early as E10.5^76^, exhibited the lowest estimated score, consistent with our observations in E15 embryos. In contrast, cell types with high regenerative potential or heightened protein synthesis demands, such as intestinal cells, hepatocytes, and megakaryocytes had the highest average estimated score. Of 93 well-represented cell types having at least 300 cells across more than 10 timepoints, we detected 66 cell types (71%) with reduced nucleolar fraction during differentiation (**Figure 6F top panel** and **Supplementary Figure 23**). Interestingly, the estimated nucleolar fraction score for hepatocytes in the reference atlas decreased both across embryogenesis and at the E15 timepoint, in contrast to the increase captured in our E15 VCS data. This discrepancy may be explained by the previously observed differences in early fetal and growth marker expression between the B6 and B6xCAST genetic backgrounds (**Supplementary Figure 3F,G**). Consistent with our E15 data, the estimated nucleolar fraction increased as chondrocytes and osteoblasts matured (**Figure 6F bottom panel, Supplementary Figure 24**). A similar increase was observed in progenitor cells, likely reflecting the high translational demands of their rapid expansion. As in our VCS data, estimated nucleolar fraction was positively correlated with proliferation in most cell types, except in ECM-producing lineages, erythroid cells and several progenitor populations (**Figure 6G** and **Supplementary Figure 25**).

Because the nucleolus is highly conserved across vertebrates^17,77^, we reasoned that our approach could be applied to other species. We thus estimated the nucleolar fraction score for a human fetal atlas^78,79^ collected by fetal dissection yielding organs labeled based on physical isolation rather than inference from single-cell transcriptomes (**Supplementary Figure 26**). The estimated nucleolar fraction score varied both within organs and among the same cell types across different organs (**Figure 7A** and **Supplementary Figure 27**). For example, the distribution of the estimated nucleolar fraction score reflected organ-specific variation in the proportions of erythroblast progenitors and mature erythrocytes (**Figure 7B**). We also noticed a discrepancy in the estimated nucleolar fraction score of pancreatic acinar cells between mouse (E16.5-18.5) and human (days 110, 117; **Figure 7C**) that is suggestive of the postnatal growth of acinar cells during murine but not human development^80^.

**Figure 7.**
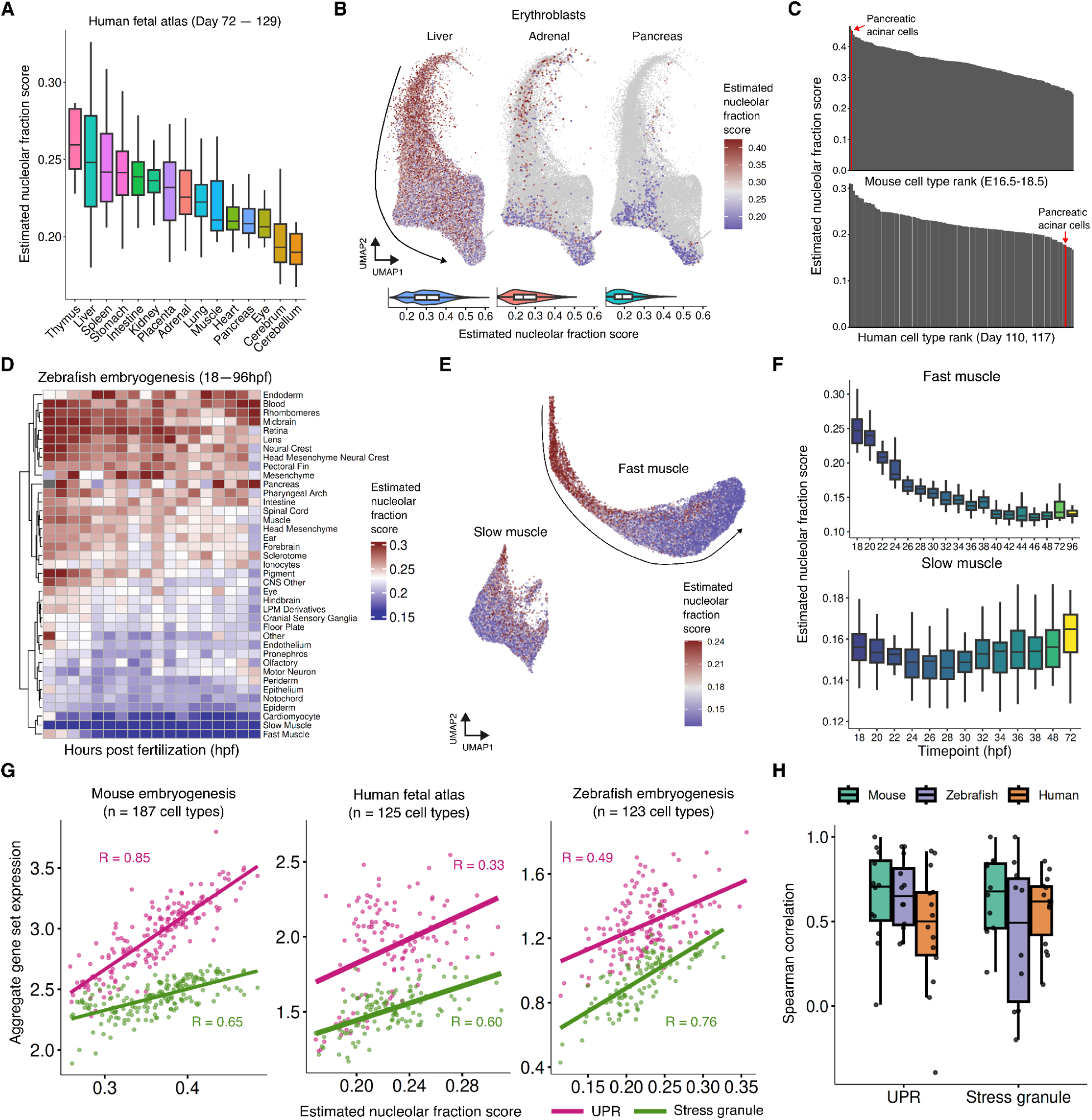
Conserved links between nucleolar fraction and cellular stress programs across vertebrate development. (A) Tissue-specific distribution of the estimated nucleolar fraction score for each physically isolated tissue within a human fetal atlas^78^. (B) UMAP embeddings (top) and violin plots (bottom) displaying the estimated nucleolar fraction score distributions of erythroblasts derived from liver, adrenal gland, and pancreas. The arrow indicates the progression of differentiation. (C) Rank plots of the estimated nucleolar fraction score for E16.5-18.5 mouse (top) and day 110, 117 human (bottom) cell types. Red bars highlight the rank differences for pancreatic acinar cells (mouse: 2nd rank of 182, human: 121st rank of 125). (D) Heatmap showing changes in the median estimated nucleolar fraction score for major tissue types across zebrafish embryogenesis^79^. (E) UMAP embeddings of zebrafish fast and slow muscle lineages colored by the estimated nucleolar fraction score, with the arrow indicating the progression of differentiation. (F) Boxplot showing the gradual decrease in the estimated nucleolar fraction score in developing fast muscle vs a uniformly low score in slow muscle. (G) Scatterplots with linear fits illustrating a positive correlation between the estimated nucleolar fraction score and the aggregate expression of UPR (pink) and stress granule (green) gene sets across mouse (left), human (middle), and zebrafish (right). Pearson correlation coefficients are indicated for each fit. (H) Distribution of Spearman correlation coefficients measuring the relationship between the estimated nucleolar fraction score and the expression of UPR and stress granule gene sets, calculated across developmental trajectories in mouse and distinct tissues in human and zebrafish.

Expanding beyond mammalian development, we estimated the nucleolar fraction score across a zebrafish embryogenesis atlas^79^ spanning 18 to 96 hours post-fertilization (**Supplementary Figure 28A,B**). Similar to the mouse embryogenesis estimates, we observed a gradual reduction in the estimated nucleolar fraction score over developmental time, with 67% of tissue types exhibiting a negative correlation (Spearman correlation < -0.3; **Figure 7D** and **Supplementary Figure 28C**). The fast muscle lineage was the most pronounced example of this trend, whereas the slow muscle lineage showed no such decline (**Figure 7E,F**). Globally, the cell types with the highest estimated nucleolar fraction score comprised a mix of highly proliferative progenitors (e.g. hepatoblasts and myeloid progenitors) and metabolically active cells (e.g. hepatocytes, erythrocytes, and pancreatic cells; **Supplementary Figure 28D**). Included among these was the hatching gland, a transient secretory organ that displayed the highest estimated nucleolar fraction score during the early 18-26 hpf stages, reflecting the massive production of proteolytic enzymes and the corresponding upregulation of unfolded protein response (UPR) machinery^81,82^.

We previously observed that an elevated nucleolar fraction score is strongly associated with stress response signatures, including the UPR and stress granules (**Figure 5B,C**). We therefore investigated whether this association is conserved across mouse, human, and zebrafish. Indeed, our estimated nucleolar fraction score positively correlated with both UPR and stress granule gene set scores across the identified cell types in all three species (**Figure 7G**). Furthermore, we found modest-to-strong positive correlations within major lineages, organs, and tissues (**Figure 7H**). Thus, activation of stress responses to maintain proteostasis in contexts requiring enhanced nucleolar function is conserved across humans, mice and zebrafish. Taken together, these analyses demonstrate that the VCS-derived nucleolar gene set serves as a robust tool for quantitatively assessing nucleolar activity in scRNA-seq datasets.

## DISCUSSION

We introduce an improved Visual Cell Sorting workflow that enables automated visual phenotyping of fixed cells or nuclei at the scale of developing mouse embryos. Because fixation allows the use of fluorescent antibodies, the method is broadly compatible with diverse cell lines and tissue-derived nuclei exhibiting a wide range of cellular and nuclear phenotypes. We applied this VCS workflow to nuclei isolated from E15 B6xCAST F1 mouse embryos to generate over 1 million single-nucleus transcriptomes from cells sorted based on nuclear, nucleolar and nuclear speckle size. To our knowledge, our dataset is the first single-cell developmental atlas annotated with subnuclear information, enabling us to explore the relationship between nuclear phenotypes and transcriptional programs across diverse cell types.

By converting VCS bin labels to cluster-level weighted average scores, we identified factors contributing to variability in nucleolar and nuclear speckle size during mouse organogenesis. Consistent with previous observations in various cell types^59,61,83^, nuclear compartment size is strongly influenced by proliferative activity and differentiation status. Our results suggest that most cell types undergo similar nuclear structural changes during differentiation, characterized by the rapid proliferation of physically larger progenitor cells followed by their maturation into smaller, more heterochromatic terminal states. These dynamics likely underlie the observed differences in nuclear compartment size across cell types, ranging from post-mitotic neurons to highly metabolically active erythroid cells.

Surprisingly, we identified cell types that deviated from these trends. For example, in differentiated fibroblasts, chondrocytes, and osteoblasts nucleoli occupied a larger fraction of the nucleus than in their less differentiated progenitors. These cell types are characterized by high levels of protein synthesis and secretion, particularly of ECM components such as collagens, fibronectin, and proteoglycans^65,84,85^. In these specialized secretory lineages, nucleolar enlargement may reflect enhanced ribosomal biogenesis and translational capacity to meet ECM production demands. During erythroid differentiation, we observed a transient rise followed by a gradual reduction in nucleolar and nuclear speckle size, preceding a sudden, asynchronous drop in proliferation. This pattern suggests a “front-loading” of ribosomal biogenesis to support the massive translational demands of early, rapid division. Intriguingly, our nucleolar fraction score correlated almost perfectly with *Bcl11a* and *Lin28b* expression. Given that Lin28b is a direct ribosome-associated protein that regulates *Bcl11a* translation^62^, we speculate that the tight correlation with the nucleolar fraction score reflects the physical association of these regulators with the increased ribosome pool to control erythroid maturation timing prior to nuclear condensation^86^.

We identified 41 genes whose expression is positively correlated with the nucleolar fraction score, indicative of upregulated transcriptional programs associated with enlarged nucleoli and increased ribosomal biogenesis. Based on this correlation, we used these genes as a signature for estimating nucleolar fraction from single-cell RNA-seq data. We used this gene set to estimate the nucleolar fraction score for mouse, human and zebrafish developmental atlases, capturing a conserved relationship between nucleolar size and stress response gene expression. These results suggest that the balance between protein synthesis, reflected by nucleolar fraction, and the cellular stress response is tightly regulated to ensure proper embryonic development. Moreover, our results highlight the potential role of stress granules in mitigating proteostatic stress during development, consistent with previous observations of stress granule formation during the highly proliferative stages of erythroid differentiation^87^. Beyond developmental atlases, our nucleolar gene signature could prove valuable in disease-relevant contexts, such as cancer^88–90^ and aging^91,92^, where nucleolar structure and function are frequently altered. However, application of the gene set to neuronal lineages should be approached with caution, as the correlation between gene set expression and the nucleolar fraction score was lower. The poor correlation was most pronounced in projection neurons, where physically larger nucleoli than estimated by our gene signature likely reflect the massive ribosomal supply required to support their extreme axon-to-soma volume ratios^93,94^.

Despite its broad utility, our improved VCS workflow has several limitations. Although VCS enables high-throughput visual phenotyping of million cells per day, VCS generates categorical bin labels for single phenotypes rather than continuous, quantitative measurements. This limitation becomes a challenge when a large fraction of cells fall in the largest or smallest bin (e.g. neuronal cells in our mouse embryo dataset), In such cases, additional VCS experiments focusing on these specific populations would be required to enhance resolution and better capture phenotypic differences. Moreover, the phenotype of interest must be pre-defined and carefully selected prior to experiments, which may constrain the discovery of unexpected cellular features. Finally, unlike *in situ* sequencing or spatial transcriptomics approaches, VCS requires dissociation of cells or nuclei from their native tissue context, resulting in the loss of spatial information that can be critical for interpreting spatially regulated gene expression patterns.

In summary, improved Visual Cell Sorting provides a versatile platform for linking image-based phenotypes to rich molecular readouts at single-cell resolution. While we illustrate its utility in the context of developmental biology, the approach should be broadly applicable to other systems where visual phenotypes are informative, including disease models and cancer biology. Importantly, we demonstrate that VCS can be integrated with sci-RNA-seq3 to achieve comprehensive transcriptional profiling of sorted populations, capturing the molecular programs underlying distinct visual states. Beyond transcriptomics, our preliminary experiments suggest that VCS is compatible with additional genomic and epigenomic assays, including DNA sequencing and ATAC-seq, enabling multimodal characterization of cell states defined by morphology or subcellular structure. Taken together, VCS offers a flexible and scalable framework for dissecting the molecular basis of complex phenotypes, bridging imaging and genomics in diverse biological contexts.

## DATA AVAILABILITY

Acquired raw images and manually annotated nuclear crops from the Visual Cell Sorting experiments will be uploaded to BioImage Archive (www.ebi.ac.uk/bioimage-archive/). Raw and processed sci-RNA-seq3 data generated from this study are in the process of being submitted to the NCBI Gene Expression Omnibus (GEO) and will be made publicly available upon publication. The published mouse, human, and zebrafish embryogenesis atlases were retrieved from https://omg.gs.washington.edu/jax/public/download.html, https://descartes.brotmanbaty.org/bbi/human-gene-expression-during-development/, and https://cole-trapnell-lab.github.io/zscape/, respectively.

## CODE AVAILABILITY

Metamorph, Python, and R scripts used for Visual Cell Sorting experiments and sci-RNA-seq3 analysis are available at https://github.com/khj3017/vcs-embryo.

## ACKNOWLEDGMENTS

We thank the past and present members of the Trapnell and Fowler labs and the UW 4D GENOME Center for helpful discussion. We also thank C. Lee and UW Pathology Flow Cytometry Core for their assistance in nuclei sorting. This work was supported by the National Institutes of Health (R35GM152106 to DMF, RM1HG010461 to DMF and CT, R01HG012761 to CT, UM1HG011586 to WSN, JS, and CMD from the National Institutes of Health Common Fund, and R35GM131745 to CMD).

## COMPETING INTERESTS

The authors compare the following competing interests: DMF is a member of the scientific advisory board of Alloz Bio and Acrobat Genomics. CT is a scientific advisory board member or consultant for Algen Biotechnologies, Altius Therapeutics and 10X Genomics. The Visual Cell Sorting method is patented (Number 12,298,297 issued May 13, 2025, “Visual Cell Sorting” Douglas Fowler, Nicholas Hasle, Sriram Pendyala, Hyeon-Jin Kim).

## METHODS

### Cell culture

U2-OS (ATCC HTB-96), A549 (ATCC CCL-185), and NIH3T3 (ATCC CRL-1658) cells were cultured in Dulbecco’s Modified Eagle Medium (DMEM) media containing 10% FBS (Hyclone, SH30071.03) and 1% Pen/Strep (Gibco, 15140-122) at 37°C with 5% CO_2_. Cells were dissociated with TrypLE Express (Thermo Fisher Scientific, 12605010) for passaging.

### F1 hybrid mouse embryo collection and staging

C57BL6/J (B6) and CAST/EIJ (CAST) mice were obtained from The Jackson Laboratory and maintained at the University of Washington (UW) Animal Facility via standard husbandry procedures under UW Animal Care and Use Committee approved protocol 201600084. As described previously^48^, timed matings between female B6 and male CAST mice were set up in the afternoon and checked the following morning. Noon of the day a plug was found was defined as E0.5 gestationally. Individual embryos were dissected from extraembryonic membranes, imaged, and snapped frozen in liquid nitrogen. A portion of each yolk sac was collected for sex-based genotyping using PCR with genomic DNA and primers specific for a Y-linked multicopy gene Ssty (Ssty-F: CTGGAGCTCTACAGTGATGA and Ssty-R: CAGTTACCAATCAACACATCAC). Samples were stored at -80 °C until further processing. To ensure reproducible and accurate comparisons across developmental stages between embryos within our sample set, we differentiated between gestational age and developmental stage of each individual embryo. Gestational age, based upon the observation of a vaginal plug, provides an estimation of the time from conception to the time of harvest for the whole litter, assuming the mice mate around midnight. On the contrary, the development of embryonic shape or morphogenesis is structured, reproducible, and distinguishable along a morphogenic trajectory for each individual embryo. A modified staging tool that exhibited better performance on E14.75–E15.0 samples was used to confirm staging of samples within this window (documentation and Python scripts available at https://github.com/marcomusy/welsh_embryo_stager). Embryos beyond E15.0 could not be automatedly staged due to limb morphology complexity toward the end of embryo development. Samples were manually binned into E0.25 increments by close inspection of limb morphogenesis.

### Antibody conjugation

To prepare photoconvertible-dye-labeled antibodies, 58 μL of Donkey Anti-Goat IgG H&L (1.25 mg/mL; Abcam, ab7120) was incubated with 9 μg of PAJF549 (Tocris, 6149) and 1 μg of Alexa 488 (Invitrogen, A20000) in 2x Borate Buffer (Thermo Scientific, 28341) for 40 mins at room temperature. The resulting conjugates were then purified using a 7kDa Zeba Spin Desalting Column (Thermo Scientific, 89882) and eluted in 85 μL PBS (70 μL of antibodies and 15 μL PBS stacker).

### Co-culture mixture experiment

For the fixed-cell mixture experiment, U2-OS cells stably expressing either H3-GFP or H3-miRFP were co-cultured at a density of 100k per well in a glass-bottom 24 well plate (Cellvis, P24-1.5H-N). Following 24 hr incubation, cells were fixed with 4% PFA (Electron Microscopy Sciences, 15710) for 10 mins and permeabilized with 0.1% Triton X-100 (Thermo Scientific, A16046-AE) for 20 mins at room temperature. To mimic VCS imaging conditions, fixed cells were maintained in 1x PBS for three days prior to dissociation. Nuclei were then dissociated from the plate via gentle scraping with a micropipette tip in sucrose nuclei buffer (0.3M sucrose (VWR, 97061-428); 3 mmol MgCl_2_ (Millipore Sigma, 68475-100ML-F); 0.5% (w/v) BSA (Bioworld, 22070038-1); 1x PBS (Thermo Scientific, 14200075)). Collected nuclei were pooled, spun down at 500g for 5 mins, and resuspended in 1 mL of hypotonic nuclei buffer for flow cytometry.

### Visual Cell Sorting equipment and settings

Imaging was performed on a Leica DMi8 inverted microscope equipped with a HC PL APO 40×1.10 W CORR CS2 objective and Adaptive Focus Control. Fluorescence excitation was provided by a Lumencor Spectra X LED Light Engine, and emissions were filtered using Semrock multi-band dichroic filters and bright-line band-pass emissions filters for DAPI (433/24 nm), GFP (520/35 nm), RFP (600/37 nm), and NIR (680/22 nm). An ASI Piezo-Z top plate (150 um range), ASI XY automated stage, and Three-Axis Stage Controller were installed to enable high-speed, automated z-stack acquisition. Patterned illumination was achieved via an Andor Mosaic3 Digital Micromirror Device, coupled to a Mosaic SS 405 nm/1.1 W laser and mapped to an iXon 888 Ultra EMCCD monochrome camera. All hardware components were integrated and controlled using the Metamorph software suite (Molecular Devices).

### Nuclei isolation, plating, and in situ reverse transcription

Flash frozen E15 mouse embryos were placed into an aluminum foil packet and powdered on a slab of dry ice with a hammer^45^. The powdered tissue was then lysed in 20 mL of the lysis buffer (3 mmol MgCl_2_, 0.5% (w/v) BSA, 0.02% IGEPAL (Millipore Sigma, I8896), 1% DEPC (Millipore Sigma, D5758), 1x PBS-hypotonic) for 10 mins on ice. The resulting homogenate was filtered with a 40 µm cell strainer, spun down at 300g for 5 min at 4°C, and resuspended in 20 mL of sucrose nuclei buffer with 1% DEPC. Nuclei were then counted and ∼400,000 nuclei were immediately plated on 24 well poly-lysine-coated glass-bottom plates via centrifugation (200g for 5 mins at 4°C). After plating, the supernatant was carefully removed and the nuclei were fixed in 4% PFA in 1x PBS for 30 mins on ice. Each well was then washed twice with 1X PBS supplemented with 0.1% SuperaseIn (Invitrogen, AM2696). Following the washes, 200 µL of reverse transcription mix (40 µL 5X RT buffer, 8 uL 10mM dNTPs (New England Biolabs, N0447L), 20 µL 40 µM well-specific RT primers, 10 µL RNAseOUT (Thermo Scientific, 10777019), 8 µL Maxima H Minus Reverse Transcriptase (200U/µL; Thermo Scientific, EP0751), 114 µL Nuclease-free water (Invitrogen, AM9932)) was added to each well and incubated overnight at 37°C. Nuclease-free water was added in between the wells to minimize evaporation.

### Immunostaining

After reverse transcription, nuclei were incubated with blocking buffer (1% w/v BSA + 0.125% Triton X-100 in 1x PBS) for 40 min at room temperature. They were then incubated for one hour with primary antibodies (all 1:300) in blocking buffer: Goat anti-H2AX (Bethyl, A303-837A), Rabbit anti-Nucleolin (Invitrogen, PA5-85972), and Mouse anti-SC35 (Abcam, ab11826). The samples were washed with PBS twice and incubated for one hour with secondary antibodies (all in 1:250) in blocking buffer: Donkey anti-mouse Alexa 647 (Abcam, ab150111) and Donkey anti-rabbit Alexa 647 (Abcam, ab150063). Finally, the samples were washed twice with PBS and stored in 1x PBS.

### Segmentation and nucleolar signal classifier training

All nuclear compartment crops used for model training were standardized to a resolution of 224×224 pixels. To generate the training datasets required for segmentation, we manually annotated approximately 1000 nucleolar and speckle crops. These were used to fine-tune a pre-trained U-Net (ResNet-34 backbone) using the *fastai*^95^ library. The model was trained for a total of 60 epochs using a 1cycle learning rate policy^96^ (model.fit_one_cycle): 15 epochs with a frozen backbone (max learning rate of 1e-4) followed by 45 unfrozen epochs (1e-6 to 5e-5). This approach yielded final validation Jaccard indices of 0.80 for nucleoli and 0.49 for nuclear speckles.

For the nucleolar signal classifier, we manually annotated nuclear crops into three categories: (1) clear nucleolar signal, (2) weak or no signal, and (3) boundary-localized signal, representing potential segmentation artifacts or fluorescence bleed-through from adjacent nuclei. A CNN-based classifier utilizing a ResNet-34 backbone was trained for a total of 40 epochs using the *fastai* package. Specifically, the classifier was trained for 10 epochs with a frozen backbone (1e-3 learning rate) followed by 30 fine-tuning epochs (1e-5 learning rate) using a 1cycle policy, achieving a final validation accuracy of 92%.

### Automated image acquisition and real-time analysis

For each field of view, a single z-plane image for H2AX and a 7-plane z-stacks (3 µm range) for the nucleolus or nuclear speckles were acquired. Real-time communication between the microscope and a dedicated GPU was established via a client-server architecture using the Remote Python Call (RPyC) library. This framework allowed the microscope to remotely execute the analysis pipeline and retrieve segmentation masks. Image analysis was performed in Python on an NVIDIA RTX 4090 GPU with 32GB DDR5 5600 MT/s. Cellpose (v2.1.0) was used for nuclear segmentation (diameter = 50, cellprop_threshold = 0.5, flow_threshold = 0.2), while the in-house trained U-Net was used for segmenting nucleolar and nuclear speckle signals on max-z projected images. For nucleolar phenotypes, nuclei lacking detectable nucleolar signals were removed using a CNN-based classifier. Finally, nuclei smaller than 1,200 pixels were excluded to remove debris, and nuclear compartments smaller than 40 pixels were discarded.

### Visual Cell Sorting

Prior to each VCS experiment, a subset of wells was imaged to establish size-based quartile cutoffs. During the experiment, images captured at each site were passed to the GPU for analysis, where nuclei were segmented and assigned to the pre-defined bins. A mask containing the binning information was generated and sent back to the microscope to guide photoconversion. Photoconversion was performed using a digital micromirror device (DMD) as described in our previous VCS approach, including DMD realignment and execution of a MetaMorph journal for quantitative photoconversion^43^. For our experiments, nuclei were photoconverted using a 405 nm laser at 100% power for 50, 175, 610, 2100 ms, corresponding to the Q1, Q2, Q3, and Q4 bins, respectively. These steps were automated and repeated across 18 wells of a 24 well plate, covering 95% of the well surface area via a 29×29 grid. Following VCS, plates were stored at 4°C. We proceeded to the sorting step once a batch of at least 72 wells (18 wells × 4 plates) were profiled, which typically took 10-12 days.

### Fluorescence-activated cell sorting

To prepare nuclei for fluorescence-activated cell sorting (FACS), each well was gently scraped with a micropipette tip in 1x PBS containing 0.33% BSA. Wells from the same imaging plate were pooled into a 15 mL conical tube and centrifuged at 600g for 5 mins. The supernatant was carefully aspirated, and the nuclei were resuspended in 1 mL of sucrose nuclei buffer containing 0.1µg/mL DAPI. Nuclei were sorted on a FACS Aria II and III (BD Biosciences). Initial gating was performed on a DAPI-A vs DAPI-H plot to isolate intact nuclei. The photoconverted populations were identified based on Alexa 488 (FITC-A) vs converted PAJF549 (PE-YG-A) signals and four-way sorted into microcentrifuge tubes (pre-coated with 5% BSA) containing 200 µL of sucrose nuclei buffer.

### sci-RNA-seq3

sci-RNA-seq3 was performed following the published protocol^45^ starting from the ligation step with few modifications.

Following sorting, an equal number of fixed NIH3T3 nuclei were spiked into the samples to help with pelleting. The nuclei were then spun down at 500g for 5 mins at 4°C and resuspended in 1200 µL sucrose nuclei buffer. Next, 11 µL of nuclei were added to each well of a 96 well plate, followed by addition of 2 µL of ligation primers and 2 µL of ligase mix (0.5 µL T4 DNA ligase (New England Biolabs, M0202M), 1.5 µL 10x T4 ligation buffer). The plate was incubated for 30 mins at room temperature, and the reaction was stopped by placing the plate on ice and adding 10 µL of cold sucrose nuclei buffer to each well. The samples were pooled into two cold 2 mL microcentrifuge tubes and centrifuged. The nuclei were then combined into a single tube and washed once with sucrose nuclei buffer supplemented with 0.5% BSA. After the wash step, the nuclei were resuspended in 600 µL of the same buffer and counted.

The nuclei were then distributed into new 96 well plates for second strand synthesis. To minimize barcode collisions, the number of nuclei that went into each well was scaled according to the number of wells imaged (RT barcodes). For 72 RT x 96 ligation barcode combinations, 1600-1700 nuclei were distributed into each well, which accounts for the spiked-in NIH3T3 cells. For second strand synthesis, 1 µL of second strand mix (0.25 µL of second strand enzyme (New England Biolabs, E6111L), 0.075 µL of buffer, 0.675 µL water) was added to each well and incubated for 3 hours at 16°C.

After second strand synthesis, 5 µL of tagmentation mix (5 µL of TD buffer, 0.025 µL of N7-loaded Tn5 (Diagenode)) was added to each well and incubated at 55°C for 5 mins. Reaction was stopped by adding 10 μL DNA binding buffer per well and incubating at room temperature for 5 min. Each well was purified using a 1.2X SPRI bead cleanup (Beckman Coulter Life Sciences, A63882) and eluting in 16 µL EB buffer (Qiagen, 19086). Indexed PCR amplification was performed by adding 24 µL of PCR mix (20 µL of 2x NEBNext PCR Master Mix (New England Biolabs, M0541L), 2 µL of indexed i5 primer, 2 µL of indexed i7 primer) to each well. Amplification was carried out using the following program: 70°C for 3 min, 98°C for 30 sec, 17 cycles of (98°C for 10 sec, 63°C for 30 sec, 72°C for 1 min) and a final 72°C for 5 min.

After the indexed PCR, products from all wells were pooled and purified using a 0.8X SPRI bead cleanup. The resulting library was quantified by Qubit and DNA ScreenTape and sequenced on the Illumina NextSeq 2000 platform using a V2 100 cycle kit (Read 1: 34 cycles, Read 2: 66 cycles, Index 1: 10 cycles, Index 2: 10 cycles).

### Processing of sci-RNA-seq3 data

Sequencing data was processed as described previously^44,97^. Briefly, reads were demultiplexed using bcl2fastq, followed by reverse transcription (RT) barcodes identification and poly(A) sequences trimming using TrimGalore (https://github.com/FelixKrueger/TrimGalore). Trimmed reads were aligned to the *mm10* genome using STAR^98^ with default settings, filtered for MAPQ ≥ 30, deduplicated, collapsed by unique molecular identifiers, and annotated. True barcodes were distinguished from debris using knee plot thresholding. The resulting reads were aggregated into a cell by gene count matrix and used to generate a Monocle3^44^ CDS object. Doublet scores were calculated for each transcriptome using Scrublet^99^, with an expected doublet rate of 0.06. Transcriptomes were removed based on the following criteria: doublet score > 0.3, number of genes expressed < 100, percentage of mitochondrial reads > 0.1, and total RNA UMIs < 200. Finally, gene expression profiles from all experiments were merged to a single Monocle3 object.

### Dimensionality reduction and cell type label transfer

For dimensionality reduction, gene expression profiles were size-factor normalized and subjected to principal component analysis (PCA) based on the top 2500 highly dispersed genes. The first 30 principal components were then used to embed the data into a three-dimensional Uniform Manifold Approximation and Projection^100^ (UMAP) space using Monocle3 (v1.4.23).

Cell type label transfer was performed as previously described^35^. Briefly, the VCS gene expression profiles were co-embedded with the E15-15.5 timepoints from the mouse embryogenesis atlas^48^. For each cell in the VCS data, 20 nearest reference neighbors were identified in the PCA space, and major trajectory and cell type labels were assigned via a majority vote. Finally, each major lineage was evaluated individually and re-annotated based on lineage-specific UMAP clustering.

### Weighted average nuclear compartment scoring

To convert categorical bin labels into quantitative scores, Leiden clustering^101^ was first performed on each major trajectory group (resolutions ranging from 5e-5 to 3e-4). To ensure equal representation of the bin labels at the embryo level, transcriptomes were then downsampled to match the size of the least abundant bin. For each cluster-embryo combination, a weighted average score was computed using the following formula:

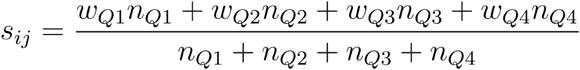

where *s_ij_* represents the weighted average score for trajectory *i* and cluster *j ^n_Q1_^* represents the number of Q1 cells in the cluster, and *w_Q1_* represents the weight associated with the Q1 bin. Weights were assigned based on the median bin-specific area or fraction values derived from the image-based distributions, ensuring that the resulting scores reflected the original size measurements. To account for well-to-well variability, weights were calculated separately for each imaging (RT) well, and the median weight was used for each cluster-embryo combination. Finally, scores were averaged across 30 downsampling iterations for downstream analyses.

### Correlation analysis of nuclear compartment size, proliferation and differentiation

Proliferation index was derived by aggregating the expression of previously reported cell cycle genes^57^. To evaluate the relationship between proliferation and nuclear compartment size, repeated measures rank correlations^67^ between the proliferation index and absolute size scores were calculated for each major trajectory group. Central nervous system (CNS) neurons were excluded from this analysis as they are post-mitotic and lacked sufficient dynamic range in the scores.

The repeated measures rank correlation (*r _rm_*) between the compartment score (*X*) and proliferation index (*Y*) for cell *j* in embryo *i* was computed as:

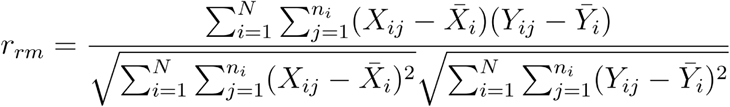

where *X _i j_* and *Y _i j_* are rank-transformed values, *n_i_* is the number of cells in embryo *i*, *N* is the total number of embryos, and *X̄_i_* and *Ȳ_i_* are the global mean ranks.

For trajectory analysis, ten distinct sub-lineages within the E15 embryo were identified based on the expression profiles of canonical marker genes from Qiu et al. 2024. Cells from these lineages were ordered along pseudotime using the “learn_graph” and “order_cells” functions in Monocle3. Similar to the proliferation analysis, repeated measures rank correlations were computed between pseudotime and the compartment scores for each identified lineage.

For the erythroid lineage, the nuclear compartment scores and proliferation index were modeled along pseudotime using cubic spline functions to pinpoint the peak values prior to the drop. To identify genes that match the dynamics of nuclear compartment size, differential expression analysis was performed using the “graph_test” function in Monocle3 to extract genes that vary significantly along pseudotime. Pearson correlations were subsequently computed between the smoothed compartment scores and gene expression profiles, filtering for genes with a correlation coefficient > 0.98.

### Identification of the nucleolar fraction transcriptomic signature

To identify a transcriptomic signature correlated with nucleolar fraction, repeated measures rank correlation coefficients were computed between individual gene expression and the nucleolar fraction score for each embryo replicate across ten distinct lineage groups. Specifically, eight sub-lineages identified for trajectory analysis and two major lineages (myeloid and epithelial) were included to capture broader cellular heterogeneity of the embryo. The GABAergic and glutamatergic neuronal sub-lineages were excluded due to limited dynamic range. We additionally computed a global repeated measures rank correlation across all defined clusters. The resulting p-values were then aggregated using Stouffer’s method^102^. Genes were retained if they met these criteria: Stouffer’s Z score > 5, rank correlation ≥ 0.3 in at least six lineages, median intra-lineage correlation ≥ 0.4, and global rank correlation ≥ 0.7. Applying these filters yielded a final set of 41 genes associated with the nucleolar fraction score, whereas no genes met these criteria for speckle fraction.

### Functional enrichment analysis

We used Hallmark^73^, GO:CC^74^, and KEGG^75^ gene sets from MSigDB for enrichment analysis. To identify gene sets associated with the nucleolar fraction score, we followed an approach analogous to the individual gene selection. Specifically, we calculated repeated measures rank correlations between the score and the aggregate expression of each gene set across lineages, and the results were combined using Stouffer’s method. Gene sets were retained if they met these following criteria: Stouffer’s Z score > 8, rank correlation > 0.3 in at least 7 lineages, median rank correlation > 0.4, and no more than two lineages with a rank correlation below -0.3. The top 30 filtered gene sets were used for Figure 5.

To functionally characterize the identified candidate genes, we performed two complementary analyses. First, gene set enrichment analysis was performed on the full correlation-ranked gene list using the R *fgsea*^103^ package. Next, over-representation analysis of the final candidate gene sets were performed using the R *clusterProfiler*^104^ package.

### Calculation of the estimated nucleolar fraction score in published atlases

To estimate nucleolar fraction in published scRNA-seq datasets, a generalized linear mixed model was fitted to assess the relationship between min-max normalized aggregate expression of nucleolar signature genes and the cluster-level VCS-derived score. The expected nucleolar fraction (*μ_ij_*) for cluster *i* within embryo *j* was modeled via a beta distribution using a complementary log-log (cloglog) link function:

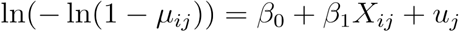

where *X_i j_* represents the normalized aggregate gene expression, β _0_is the global fixed intercept, β _1_ is the fixed effect size of the gene signature, and *u _j_*is the random intercept for embryo *j*.

The fixed effects of this model were then used to estimate the nucleolar fraction score in individual cells of non-VCS datasets. For human and zebrafish atlases, we converted the mouse nucleolar fraction genes to their respective orthologs using the R orthogene^105^ package and manual review, yielding 40 and 34 orthologous genes, respectively (**Supplementary Table 6**).

### Temporal and correlation analysis of embryogenesis atlases

For the analysis of embryogenesis atlases, the estimated nucleolar fraction score was first aggregated by calculating the median value at the embryo replicate level, partitioned by cell type and species-specific groupings (mouse: major trajectory group, human: organ, zebrafish: tissue). These replicate-level values were subsequently collapsed across embryos to derive a final consensus median value for each cell type within its respective trajectory, organ, or tissue.

To evaluate the temporal dynamics of the estimated nucleolar fraction score during mouse embryogenesis, the score was modeled as a function of time using cubic spline regression. The significance of each fit was assessed by comparing the spline models against null models using ANOVA F-tests. Smoothed temporal profiles generated from these fits were then hierarchically clustered to identify cell types that share similar dynamics.

To identify cell types where the estimated nucleolar fraction score was uncoupled from proliferation within the mouse embryogenesis atlas, natural cubic splines were fitted to both the nucleolar fraction and proliferation index scores for cell types comprising over 300 cells across at least 10 time points. The fitted curves were evaluated to locate descending segments, defined by peak-to-trough boundaries. Within these specific windows, localized Spearman rank correlations were calculated. Cell types were classified as having an inverse relationship if they met these three criteria: (1) a minimum of 5% decrease in the fitted curve, (2) negative slope spanning over 30% of the evaluated proliferation range, and (3) segment rank correlation ≤ -0.3.

## REFERENCES

1. Belmont, A. S. Nuclear compartments: An incomplete primer to nuclear compartments, bodies, and genome organization relative to nuclear architecture. Cold Spring Harb. Perspect. Biol. 14, a041268 (2022).

2. Staněk, D. & Fox, A. H. Nuclear bodies: news insights into structure and function. Curr. Opin. Cell Biol. 46, 94–101 (2017).

3. Shan, L., Li, P., Yu, H. & Chen, L.-L. Emerging roles of nuclear bodies in genome spatial organization. Trends Cell Biol. 34, 595–605 (2024).

4. Németh, A. et al. Initial genomics of the human nucleolus. PLoS Genet. 6, e1000889 (2010).

5. van Steensel, B. & Belmont, A. S. Lamina-associated domains: Links with chromosome architecture, heterochromatin, and gene repression. Cell 169, 780–791 (2017).

6. Ahanger, S. H. et al. Distinct nuclear compartment-associated genome architecture in the developing mammalian brain. Nat. Neurosci. 24, 1235–1242 (2021).

7. Chen, Y. et al. Mapping 3D genome organization relative to nuclear compartments using TSA-Seq as a cytological ruler. J. Cell Biol. 217, 4025–4048 (2018).

8. Quinodoz, S. A. et al. Higher-order inter-chromosomal hubs shape 3D genome organization in the nucleus. Cell 174, 744–757.e24 (2018).

9. Abraham, K. J. et al. Nucleolar RNA polymerase II drives ribosome biogenesis. Nature 585, 298–302 (2020).

10. Kurihara, M. et al. Genomic profiling by ALaP-seq reveals transcriptional regulation by PML bodies through DNMT3A exclusion. Mol. Cell 78, 493–505.e8 (2020).

11. Chiodi, I. et al. RNA recognition motif 2 directs the recruitment of SF2/ASF to nuclear stress bodies. Nucleic Acids Res. 32, 4127–4136 (2004).

12. Lamond, A. I. & Spector, D. L. Nuclear speckles: a model for nuclear organelles. Nat. Rev. Mol. Cell Biol. 4, 605–612 (2003).

13. Bhat, P. et al. Genome organization around nuclear speckles drives mRNA splicing efficiency. Nature 629, 1165–1173 (2024).

14. Reeder, R. H. rRNA synthesis in the nucleolus. Trends Genet. 6, 390–395 (1990).

15. Hayashi, Y. et al. Nucleolar integrity during interphase supports faithful Cdk1 activation and mitotic entry. Sci. Adv. 4, eaap7777 (2018).

16. Lindström, M. S. et al. Nucleolus as an emerging hub in maintenance of genome stability and cancer pathogenesis. Oncogene 37, 2351–2366 (2018).

17. Boulon, S., Westman, B. J., Hutten, S., Boisvert, F.-M. & Lamond, A. I. The nucleolus under stress. Mol. Cell 40, 216–227 (2010).

18. Yang, K. et al. A redox mechanism underlying nucleolar stress sensing by nucleophosmin. Nat. Commun. 7, 13599 (2016).

19. van Koningsbruggen, S. et al. High-resolution whole-genome sequencing reveals that specific chromatin domains from most human chromosomes associate with nucleoli. Mol. Biol. Cell 21, 3735–3748 (2010).

20. Pontvianne, F. et al. Identification of nucleolus-associated chromatin domains reveals a role for the nucleolus in 3D organization of the A. thaliana genome. Cell Rep. 16, 1574–1587 (2016).

21. Peng, T. et al. Mapping nucleolus-associated chromatin interactions using nucleolus Hi-C reveals pattern of heterochromatin interactions. Nat. Commun. 14, 350 (2023).

22. Spector, D. L. & Lamond, A. I. Nuclear speckles. Cold Spring Harb. Perspect. Biol. 3, (2011).

23. Galganski, L., Urbanek, M. O. & Krzyzosiak, W. J. Nuclear speckles: molecular organization, biological function and role in disease. Nucleic Acids Res. 45, 10350–10368 (2017).

24. Meshorer, E. & Misteli, T. Chromatin in pluripotent embryonic stem cells and differentiation. Nat. Rev. Mol. Cell Biol. 7, 540–546 (2006).

25. Gupta, S. & Santoro, R. Regulation and Roles of the Nucleolus in Embryonic Stem Cells: From Ribosome Biogenesis to Genome Organization. Stem Cell Reports 15, 1206–1219 (2020).

26. Eng, C.-H. L. et al. Transcriptome-scale super-resolved imaging in tissues by RNA seqFISH. Nature 568, 235–239 (2019).

27. Chen, K. H., Boettiger, A. N., Moffitt, J. R., Wang, S. & Zhuang, X. RNA imaging. Spatially resolved, highly multiplexed RNA profiling in single cells. Science 348, aaa6090 (2015).

28. Lee, J. H. et al. Fluorescent in situ sequencing (FISSEQ) of RNA for gene expression profiling in intact cells and tissues. Nat. Protoc. 10, 442–458 (2015).

29. Codeluppi, S. et al. Spatial organization of the somatosensory cortex revealed by osmFISH. Nat. Methods 15, 932–935 (2018).

30. Marshall, J. L. et al. HyPR-seq: Single-cell quantification of chosen RNAs via hybridization and sequencing of DNA probes. Proc. Natl. Acad. Sci. U. S. A. 117, 33404–33413 (2020).

31. Wang, F. et al. RNAscope: a novel in situ RNA analysis platform for formalin-fixed, paraffin-embedded tissues. J. Mol. Diagn. 14, 22–29 (2012).

32. Ståhl, P. L. et al. Visualization and analysis of gene expression in tissue sections by spatial transcriptomics. Science 353, 78–82 (2016).

33. Rodriques, S. G. et al. Slide-seq: A scalable technology for measuring genome-wide expression at high spatial resolution. Science 363, 1463–1467 (2019).

34. Stickels, R. R. et al. Highly sensitive spatial transcriptomics at near-cellular resolution with Slide-seqV2. Nat. Biotechnol. 39, 313–319 (2021).

35. Srivatsan, S. R. et al. Embryo-scale, single-cell spatial transcriptomics. Science 373, 111–117 (2021).

36. Cho, C.-S. et al. Microscopic examination of spatial transcriptome using Seq-Scope. Cell 184, 3559–3572.e22 (2021).

37. Vickovic, S. et al. High-definition spatial transcriptomics for in situ tissue profiling. Nat. Methods 16, 987–990 (2019).

38. Fu, X. et al. Polony gels enable amplifiable DNA stamping and spatial transcriptomics of chronic pain. Cell 185, 4621–4633.e17 (2022).

39. Liu, Y. et al. High-spatial-resolution multi-omics sequencing via deterministic barcoding in tissue. Cell 183, 1665–1681.e18 (2020).

40. Lee, Y. et al. XYZeq: Spatially resolved single-cell RNA sequencing reveals expression heterogeneity in the tumor microenvironment. Sci. Adv. 7, (2021).

41. Poovathingal, S. et al. Nova-ST: Nano-patterned ultra-dense platform for spatial transcriptomics. *Cell Rep*. Methods 4, 100831 (2024).

42. Schott, M. et al. Open-ST: High-resolution spatial transcriptomics in 3D. Cell 187, 3953–3972.e26 (2024).

43. Hasle, N. et al. High-throughput, microscope-based sorting to dissect cellular heterogeneity. Mol. Syst. Biol. 16, e9442 (2020).

44. Cao, J. et al. The single-cell transcriptional landscape of mammalian organogenesis. Nature 566, 496–502 (2019).

45. Martin, B. K. et al. Optimized single-nucleus transcriptional profiling by combinatorial indexing. Nat. Protoc. 18, 188–207 (2023).

46. Gurskaya, N. G. et al. Engineering of a monomeric green-to-red photoactivatable fluorescent protein induced by blue light. Nat. Biotechnol. 24, 461–465 (2006).

47. Grimm, J. B. et al. A general method to fine-tune fluorophores for live-cell and in vivo imaging. Nat. Methods 14, 987–994 (2017).

48. Qiu, C. et al. A single-cell time-lapse of mouse prenatal development from gastrula to birth. Nature 626, 1084–1093 (2024).

49. Goncalves, A. et al. Extensive compensatory cis-trans regulation in the evolution of mouse gene expression. Genome Res. 22, 2376–2384 (2012).

50. Matthews, B. J., Melia, T. & Waxman, D. J. Harnessing natural variation to identify cis regulators of sex-biased gene expression in a multi-strain mouse liver model. PLoS Genet. 17, e1009588 (2021).

51. Naso, R. Variations in rat liver chromatin composition during growth and the heterotic response. Biochem. Genet. 14, 281–292 (1976).

52. Ilik, İ. A., et al. SON and SRRM2 are essential for nuclear speckle formation. Elife 9, (2020).

53. Dundr, M. & Misteli, T. Biogenesis of nuclear bodies. Cold Spring Harb. Perspect. Biol. 2, a000711 (2010).

54. Gholamalamdari, O. et al. Major nuclear locales define nuclear genome organization and function beyond A and B compartments. Elife 13, RP99116 (2025).

55. Jevtić, P., Edens, L. J., Vuković, L. D. & Levy, D. L. Sizing and shaping the nucleus: mechanisms and significance. Curr. Opin. Cell Biol. 28, 16–27 (2014).

56. Balachandra, S., Sarkar, S. & Amodeo, A. A. The nuclear-to-cytoplasmic ratio: Coupling DNA content to cell size, cell cycle, and biosynthetic capacity. Annu. Rev. Genet. 56, 165–185 (2022).

57. Tirosh, I. et al. Dissecting the multicellular ecosystem of metastatic melanoma by single-cell RNA-seq. Science 352, 189–196 (2016).

58. Schulze, R. J., Schott, M. B., Casey, C. A., Tuma, P. L. & McNiven, M. A. The cell biology of the hepatocyte: A membrane trafficking machine. J. Cell Biol. 218, 2096–2112 (2019).

59. Homma, S., Beermann, M. L., Yu, B., Boyce, F. M. & Miller, J. B. Nuclear bodies reorganize during myogenesis in vitro and are differentially disrupted by expression of FSHD-associated DUX4. Skelet. Muscle 6, 42 (2016).

60. Baker, N. E. Developmental regulation of nucleolus size during Drosophila eye differentiation. PLoS One 8, e58266 (2013).

61. Potolitsyna, E. et al. Cytoskeletal rearrangement precedes nucleolar remodeling during adipogenesis. *Commun*. Biol. 7, 458 (2024).

62. Basak, A. et al. Control of human hemoglobin switching by LIN28B-mediated regulation of BCL11A translation. Nat. Genet. 52, 138–145 (2020).

63. Mackie, E. J., Ahmed, Y. A., Tatarczuch, L., Chen, K.-S. & Mirams, M. Endochondral ossification: how cartilage is converted into bone in the developing skeleton. Int. J. Biochem. Cell Biol. 40, 46–62 (2008).

64. Aghajanian, P. & Mohan, S. The art of building bone: emerging role of chondrocyte-to-osteoblast transdifferentiation in endochondral ossification. Bone Res. 6, 19 (2018).

65. Tracy, L. E., Minasian, R. A. & Caterson, E. J. Extracellular matrix and dermal fibroblast function in the healing wound. Adv. Wound Care (New Rochelle) 5, 119–136 (2016).

66. Li, B. & Wang, J. H.-C. Fibroblasts and myofibroblasts in wound healing: force generation and measurement. J. Tissue Viability 20, 108–120 (2011).

67. Bakdash, J. Z. & Marusich, L. R. Repeated measures correlation. Front. Psychol. 8, 456 (2017).

68. Goldfinger, M., Shmuel, M., Benhamron, S. & Tirosh, B. Protein synthesis in plasma cells is regulated by crosstalk between endoplasmic reticulum stress and mTOR signaling. Eur. J. Immunol. 41, 491–502 (2011).

69. Malhotra, J. D. et al. Antioxidants reduce endoplasmic reticulum stress and improve protein secretion. Proc. Natl. Acad. Sci. U. S. A. 105, 18525–18530 (2008).

70. Torrence, M. E. et al. The mTORC1-mediated activation of ATF4 promotes protein and glutathione synthesis downstream of growth signals. Elife 10, e63326 (2021).

71. Li, C. et al. The role of lncRNA MALAT1 in the regulation of hepatocyte proliferation during liver regeneration. Int. J. Mol. Med. 39, 347–356 (2017).

72. Malakar, P. et al. Long noncoding RNA MALAT1 promotes hepatocellular carcinoma development by SRSF1 upregulation and mTOR activation. Cancer Res. 77, 1155–1167 (2017).

73. Liberzon, A. et al. The Molecular Signatures Database Hallmark Gene Set Collection. Cell Systems vol. 1 417–425 Preprint at 10.1016/j.cels.2015.12.004 (2015).

74. Gene Ontology Consortium. The Gene Ontology knowledgebase in 2026. Nucleic Acids Res. 54, D1779–D1792 (2026).

75. Kanehisa, M. & Goto, S. KEGG: kyoto encyclopedia of genes and genomes. Nucleic Acids Res. 28, 27–30 (2000).

76. Patel, T. et al. Transcriptional dynamics of murine motor neuron maturation in vivo and in vitro. Nat. Commun. 13, 5427 (2022).

77. Pederson, T. The nucleolus. Cold Spring Harb. Perspect. Biol. 3, a000638–a000638 (2011).

78. Cao, J. et al. A human cell atlas of fetal gene expression. Science 370, (2020).

79. Saunders, L. M. et al. Embryo-scale reverse genetics at single-cell resolution. Nature 623, 782–791 (2023).

80. Anzi, S. et al. Postnatal exocrine pancreas growth by cellular hypertrophy correlates with a shorter lifespan in mammals. Dev. Cell 45, 726–737.e3 (2018).

81. Wang, Y. et al. Gene module reconstruction identifies cellular differentiation processes and the regulatory logic of specialized secretion in zebrafish. Dev. Cell 60, 581–598.e9 (2025).

82. Dorrity, M. W. et al. Proteostasis governs differential temperature sensitivity across embryonic cell types. Cell 186, 5015–5027.e12 (2023).

83. Pham, A. T., Mani, M., Wang, X. & Vafabakhsh, R. Multiscale biophysical analysis of nucleolus disassembly during mitosis. Proc. Natl. Acad. Sci. U. S. A. 121, e2312250121 (2024).

84. Akkiraju, H. & Nohe, A. Role of chondrocytes in cartilage formation, progression of Osteoarthritis and cartilage regeneration. J. Dev. Biol. 3, 177–192 (2015).

85. El-Amin, S. F. et al. Extracellular matrix production by human osteoblasts cultured on biodegradable polymers applicable for tissue engineering. Biomaterials 24, 1213–1221 (2003).

86. Chen, K. et al. Resolving the distinct stages in erythroid differentiation based on dynamic changes in membrane protein expression during erythropoiesis. Proc. Natl. Acad. Sci. U. S. A. 106, 17413–17418 (2009).

87. Ghisolfi, L., Dutt, S., McConkey, M. E., Ebert, B. L. & Anderson, P. Stress granules contribute to α-globin homeostasis in differentiating erythroid cells. Biochem. Biophys. Res. Commun. 420, 768–774 (2012).

88. Zink, D., Fischer, A. H. & Nickerson, J. A. Nuclear structure in cancer cells. Nat. Rev. Cancer 4, 677–687 (2004).

89. Montanaro, L., Treré, D. & Derenzini, M. Nucleolus, ribosomes, and cancer. Am. J. Pathol. 173, 301–310 (2008).

90. Belin, S. et al. Antiproliferative effect of ascorbic acid is associated with the inhibition of genes necessary to cell cycle progression. PLoS One 4, e4409 (2009).

91. Buchwalter, A. & Hetzer, M. W. Nucleolar expansion and elevated protein translation in premature aging. Nat. Commun. 8, 328 (2017).

92. Gutierrez, J. I. & Tyler, J. K. A mortality timer based on nucleolar size triggers nucleolar integrity loss and catastrophic genomic instability. *Nat*. Aging 4, 1782–1793 (2024).

93. Hetman, M. & Pietrzak, M. Emerging roles of the neuronal nucleolus. Trends Neurosci. 35, 305–314 (2012).

94. Holt, C. E., Martin, K. C. & Schuman, E. M. Local translation in neurons: visualization and function. Nat. Struct. Mol. Biol. 26, 557–566 (2019).

95. Howard, J. & Gugger, S. Fastai: A Layered API for Deep Learning. Information 11, 108 (2020).

96. Smith, L. N. A disciplined approach to neural network hyper-parameters: Part 1 -- learning rate, batch size, momentum, and weight decay. arXiv [cs.LG] (2018) doi:10.48550/arXiv.1803.09820.

97. Kim, H.-J. et al. Nuclear oligo hashing improves differential analysis of single-cell RNA-seq. Nat. Commun. 13, 2666 (2022).

98. Dobin, A. et al. STAR: ultrafast universal RNA-seq aligner. Bioinformatics vol. 29 15–21 Preprint at 10.1093/bioinformatics/bts635 (2013).

99. Wolock, S. L., Lopez, R. & Klein, A. M. Scrublet: Computational identification of cell Doublets in Single-cell transcriptomic data. Cell Syst. 8, 281–291.e9 (2019).

100. McInnes, L., Healy, J. & Melville, J. UMAP: Uniform Manifold Approximation and Projection for Dimension Reduction. arXiv [stat.ML] (2018).

101. Traag, V. A., Waltman, L. & van Eck, N. J. From Louvain to Leiden: guaranteeing well-connected communities. Sci. Rep. 9, 5233 (2019).

102. Stouffer, S., Suchman, E., DeVinney, L. C., Star, S. & Williams, R. The American soldier: Adjustment during army life. (Studies in social psychology in World War II), Vol. 1. Oxford, England: Princeton Univ. Press The American soldier: Adjustment during army life. (Studies in social psychology in World War II) 1, 599 (1949).

103. Korotkevich, G. et al. Fast gene set enrichment analysis. bioRxiv (2016) doi:10.1101/060012.

104. Yu, G., Wang, L.-G., Han, Y. & He, Q.-Y. clusterProfiler: an R package for comparing biological themes among gene clusters. OMICS 16, 284–287 (2012).

105. Schilder, B. M., Murphy, A. E. & Skene, N. G. orthogene: a Bioconductor package to easily map genes within and across hundreds of species. bioRxiv (2026) doi:10.64898/2026.01.17.700094.

106. Stringer, C., Wang, T., Michaelos, M. & Pachitariu, M. Cellpose: a generalist algorithm for cellular segmentation. Nat. Methods 18, 100–106 (2021).

