## Supplemental Information for "Embryo-scale Visual Cell Sorting reveals a conserved transcriptomic signature of nucleolar size linked to proteostasis"

### TABLE OF CONTENTS

**Supplementary Figure 1.** Improved Visual Cell Sorting is compatible with cell fixation.

**Supplementary Figure 2.** Fidelity of sci-RNA-seq3 profiles in visually sorted nuclei compared to the reference data.

**Supplementary Figure 3.** Transcriptome comparison of erythroid and hepatocyte lineages between the reference (B6) and VCS (B6xCAST) data.

**Supplementary Figure 4.** Visual Cell Sorting of E15 mouse embryos.

**Supplementary Figure 5.** Performance of the nucleolar signal classifier and nuclear compartment segmentation models.

**Supplementary Figure 6.** Quality control and reference comparison of E15 VCS-sci-RNA-seq3 data.

**Supplementary Figure 7.** Comparison of VCS bins with nuclear compartment markers and lineage-specific distribution across bins.

**Supplementary Figure 8.** Distribution of the VCS-derived nuclear compartment scores.

**Supplementary Figure 9.** Lineage-specific comparison of the absolute size scores against the proliferation index.

**Supplementary Figure 10.** Marker gene expression and pseudotime trajectory of E15 embryonic lineages (from astrocytes to GABAergic neurons).

**Supplementary Figure 11.** Marker gene expression and pseudotime trajectory of E15 embryonic lineages (from glutamatergic to upper-layer neurons).

**Supplementary Figure 12.** Mapping of the nuclear compartment scores across E15 embryonic lineages (from astrocytes to GABAergic neurons).

**Supplementary Figure 13.** Mapping of the nuclear compartment scores across E15 embryonic lineages (from glutamatergic to upper-layer neurons).

**Supplementary Figure 14.** Lineage-specific comparison of the nuclear compartment size scores against differentiation.

**Supplementary Figure 15.** Dynamics of the nuclear compartment size scores during erythroid differentiation.

**Supplementary Figure 16.** Pairwise correlations of the nuclear compartment size scores.

**Supplementary Figure 17.** Relationship between gene expression and the nucleolar fraction score for the 41 candidate genes.

**Supplementary Figure 18.** Intra-lineage correlation between the nucleolar fraction score and the 41 candidate genes.

**Supplementary Figure 19.** Functional enrichment analysis of the 41-gene nucleolar signature.

**Supplementary Figure 20.** Genes negatively correlated with the nucleolar fraction score.

**Supplementary Figure 21.** Concordance between the estimated and measured nucleolar fraction score.

**Supplementary Figure 22.** Mapping of the nucleolar fraction score across the major lineages in the mouse embryogenesis atlas.

**Supplementary Figure 23.** Dynamics of the estimated nucleolar fraction score across mouse embryonic development.

**Supplementary Figure 24.** Cell types positively correlated between the estimated nucleolar fraction score and developmental time.

**Supplementary Figure 25.** Cell types negatively correlated with the estimated nucleolar fraction score and proliferation index.

**Supplementary Figure 26.** Mapping of the nucleolar fraction score across organs in the human fetal atlas.

**Supplementary Figure 27.** Comparison of the estimated nucleolar fraction score in cell types shared across organs.

**Supplementary Figure 28.** Estimation of the nucleolar fraction score on the zebrafish embryogenesis atlas.

### SUPPLEMENTARY TABLES (*Provided as a separate Excel file*)

**Supplementary Table 1.** Reagents and resources used for Visual Cell Sorting and sci-RNA-seq3.

**Supplementary Table 2.** Summary of embryos used for VCS experiments.

**Supplementary Table 3.** Transcriptomic signature of genes positively associated with the nucleolar fraction score.

**Supplementary Table 4.** Functional annotation of 41-gene nucleolar fraction signature.

**Supplementary Table 5.** Genes negatively correlated with the nucleolar fraction score.

**Supplementary Table 6.** Human and zebrafish orthologs of the nucleolar fraction signature.

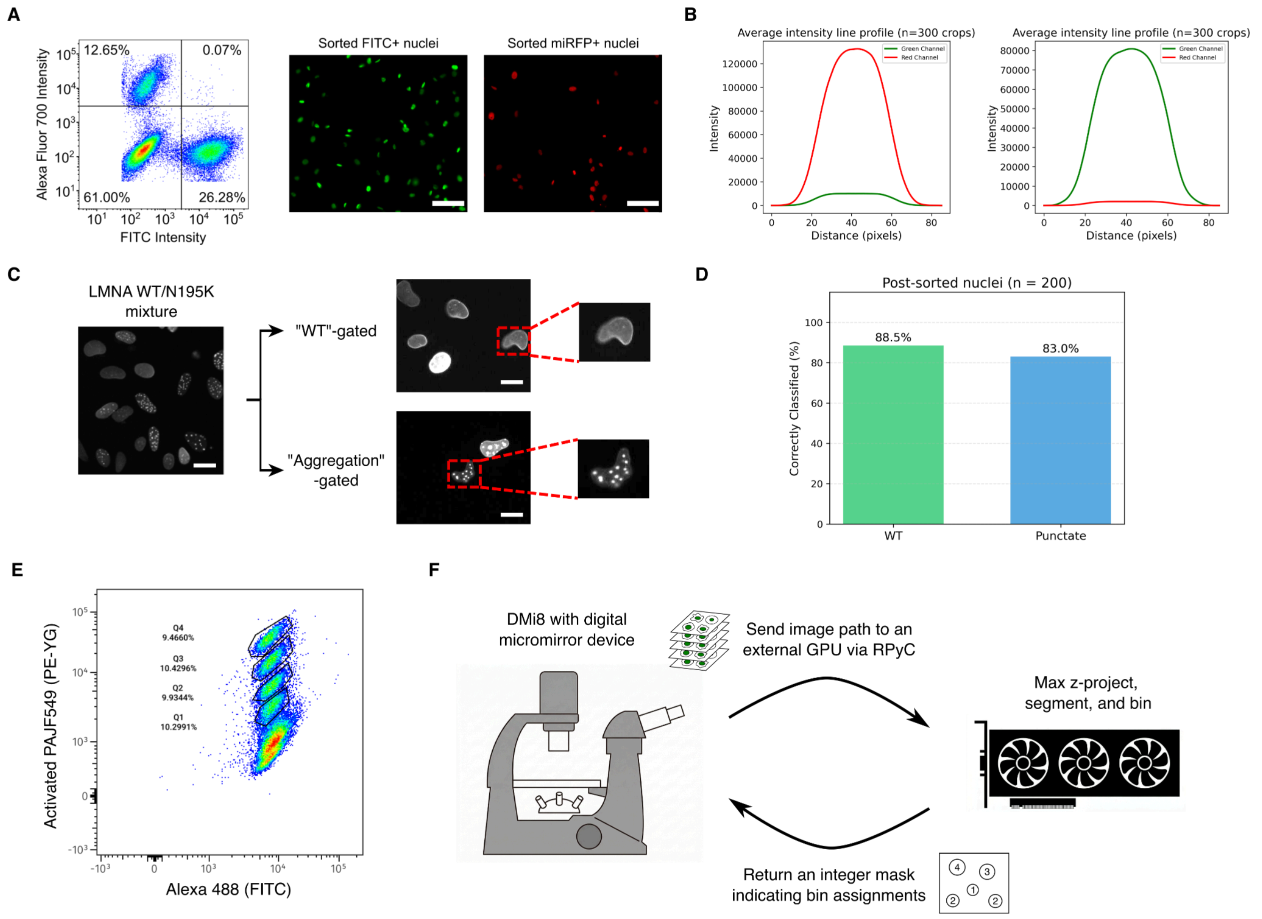


**Supplementary Figure 1. Improved Visual Cell Sorting is compatible with cell fixation.** (A) Flow cytometry of co-cultured, 4% PFA-fixed GFP- and RFP-expressing cells, yielding a 1% doublet rate (left). Fluorescence imaging of the post-sorted nuclei confirms purity (right). Scale bar = 5 µm. (B) Average line intensity profiles of sorted miRFP- and GFP-positive cells (n = 300 crops). (C) Pilot VCS experiment with a mixture of U2OS cells over-expressing WT (diffuse) or N195K (aggregate) *LMNA* variants. These cells were sorted based on the *LMNA* phenotype. (D) Quantification of post-sorted U2OS nuclei based on manual *LMNA* phenotype classification. (E) Flow cytometry plot of randomly photoactivated A549 cells stained with antibodies co-labeled with Alexa 488 and PAJF549. (F) Schematic of the improved VCS image analysis pipeline. We wrote custom Metamorph and Python scripts to enable communication between the microscope and an external computer.

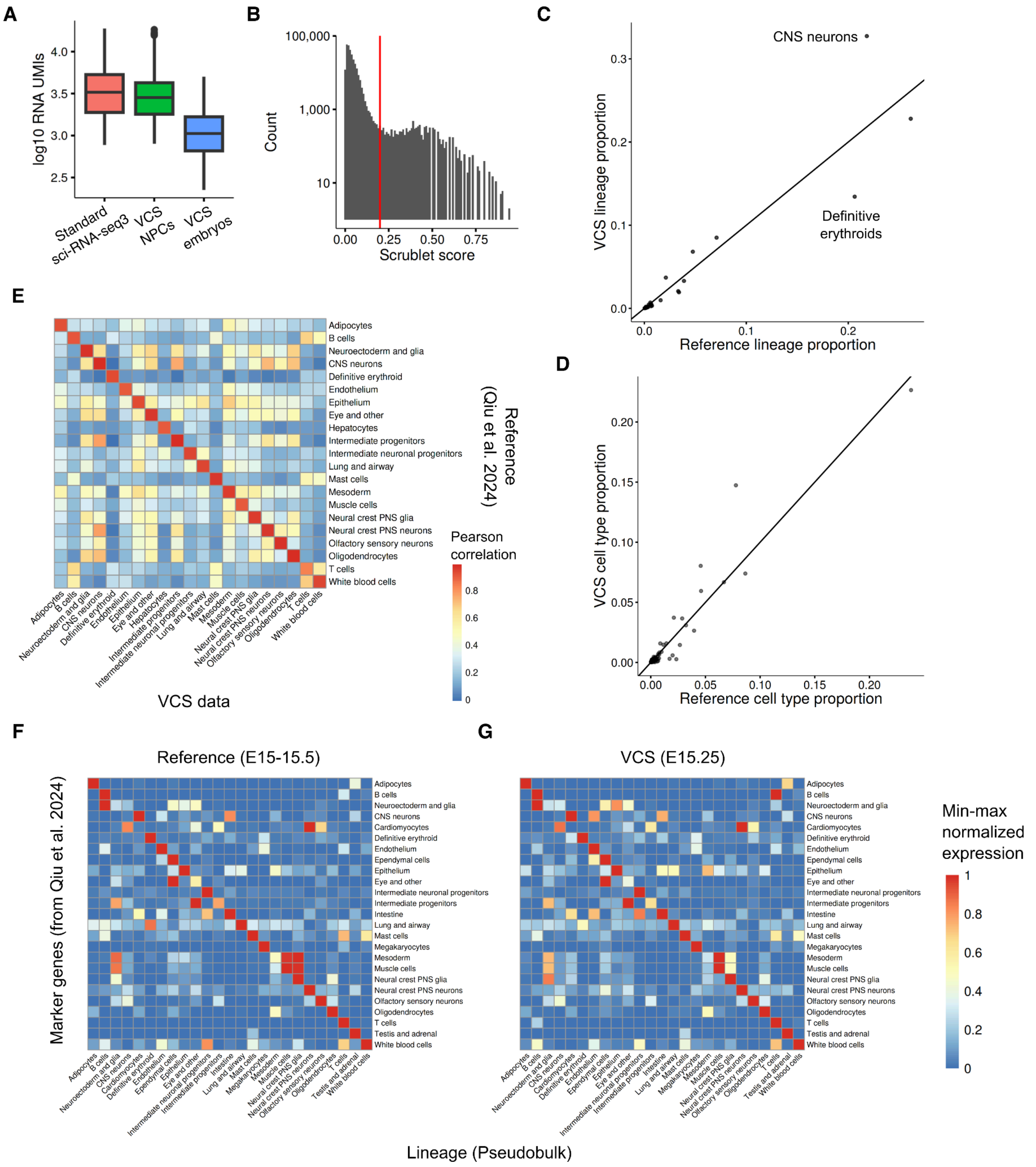


**Supplementary Figure 2. Fidelity of sci-RNA-seq3 profiles in visually sorted nuclei compared to the reference data.** (A) Boxplots of total RNA UMIs recovered comparing standard sci-RNA-seq3 vs VCS-sci-RNA-seq3. (B) Distribution of Scrublet doublet scores from a mock embryo VCS experiment. (C,D) Concordance of lineage (major trajectory group; C) and cell type (D) proportions between the reference and the mock VCS-sci-RNA-seq3 data. The diagonal line represents y = x. (E) Heatmap of lineage-grouped pseudobulk expression profiles comparing standard and visually cell sorted nuclei. (F,G) Expression of canonical marker genes (from Qiu et al. 2024) across lineages for the reference (F) and VCS (G) data.

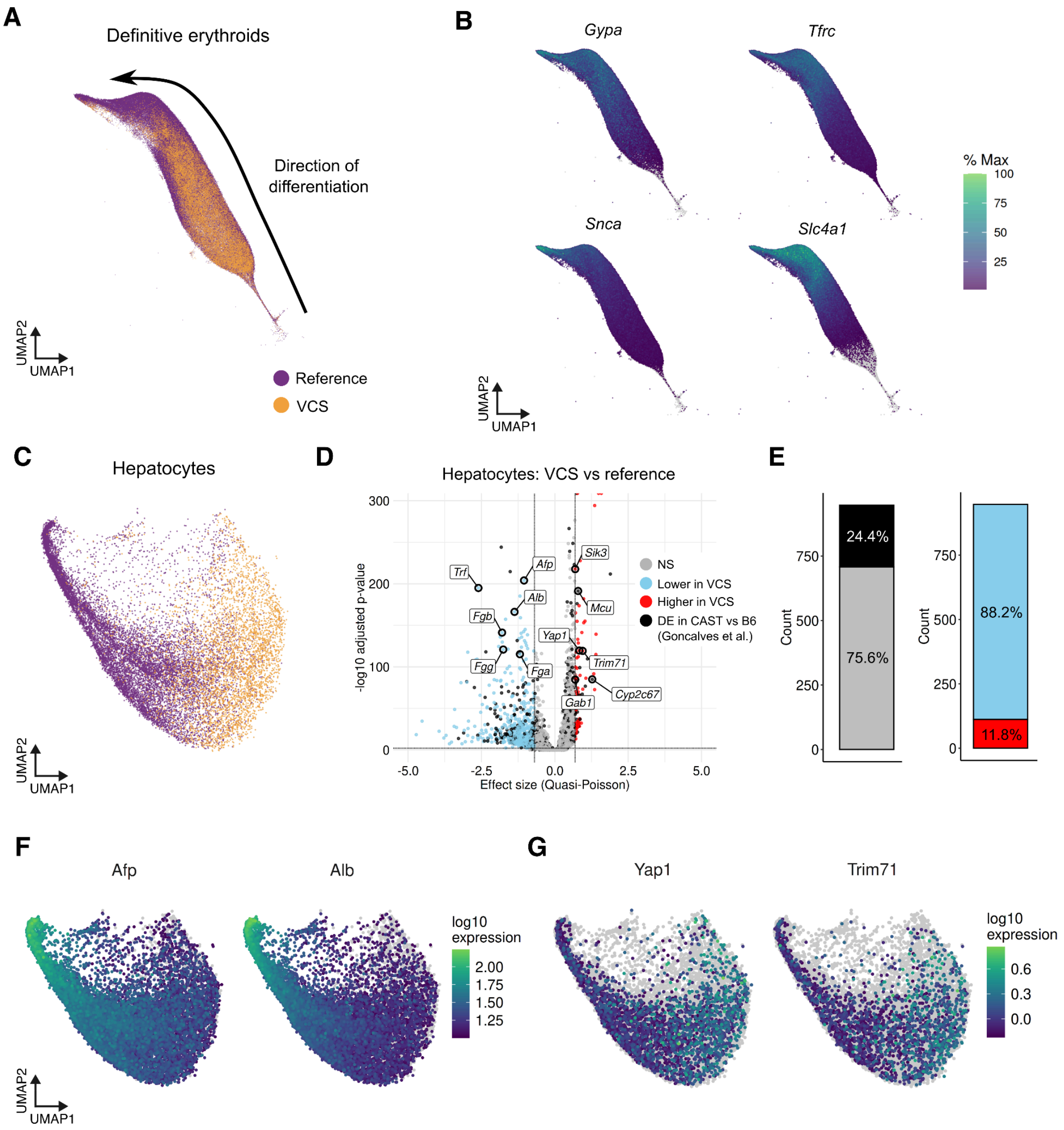


**Supplementary Figure 3. Transcriptome comparison of erythroid and hepatocyte lineages between the reference (B6) and VCS (B6xCAST) data.** (A,C) UMAP embeddings of erythroids (A) and hepatocytes (C), colored by dataset. (B) Expression levels of the canonical erythroid marker genes. (D) Volcano plot of differentially expressed genes (DEGs) in hepatocytes between reference (B6) and VCS (B6xCAST). (E) Barplots summarizing the overlap of detected DEGs with known strain-specific expression signatures (Goncalves et al.; left) and the total number of down- and up-regulated genes in the VCS data (right). (F,G) Expression levels of early hepatocyte markers (*Afp* and *Alb*) with increased expression in the B6 reference (F) and genes associated with organ growth and developmental timing (*Yap1* and *Trim71*) with increased expression in the B6xCAST VCS data (G).

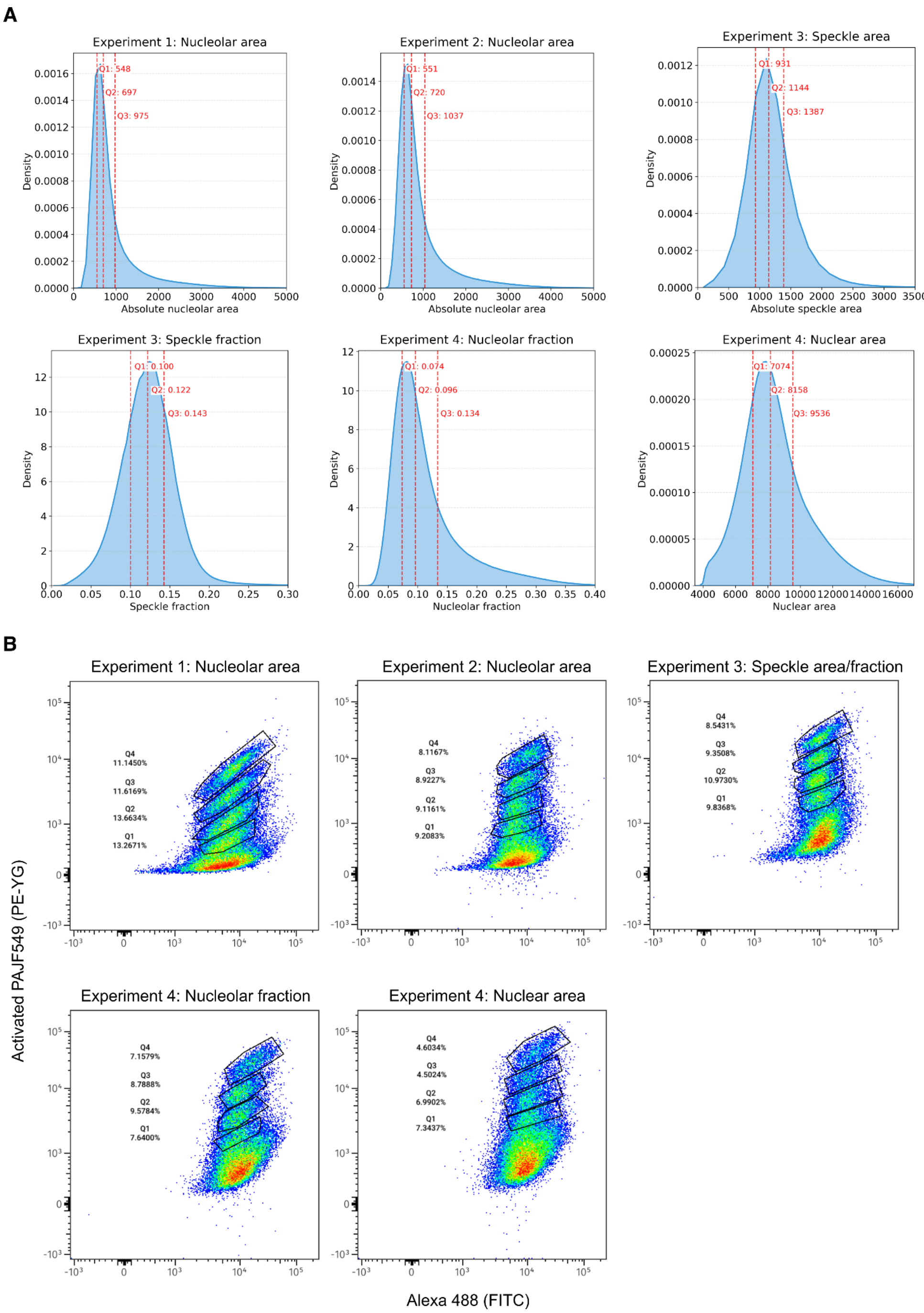


**Supplementary Figure 4. Visual Cell Sorting of E15 mouse embryos.** (A) Distributions of nuclear compartment phenotypes, indicating the average cutoff values applied across experiments. (B) Flow plots and gating used to sort the photoactivated populations.

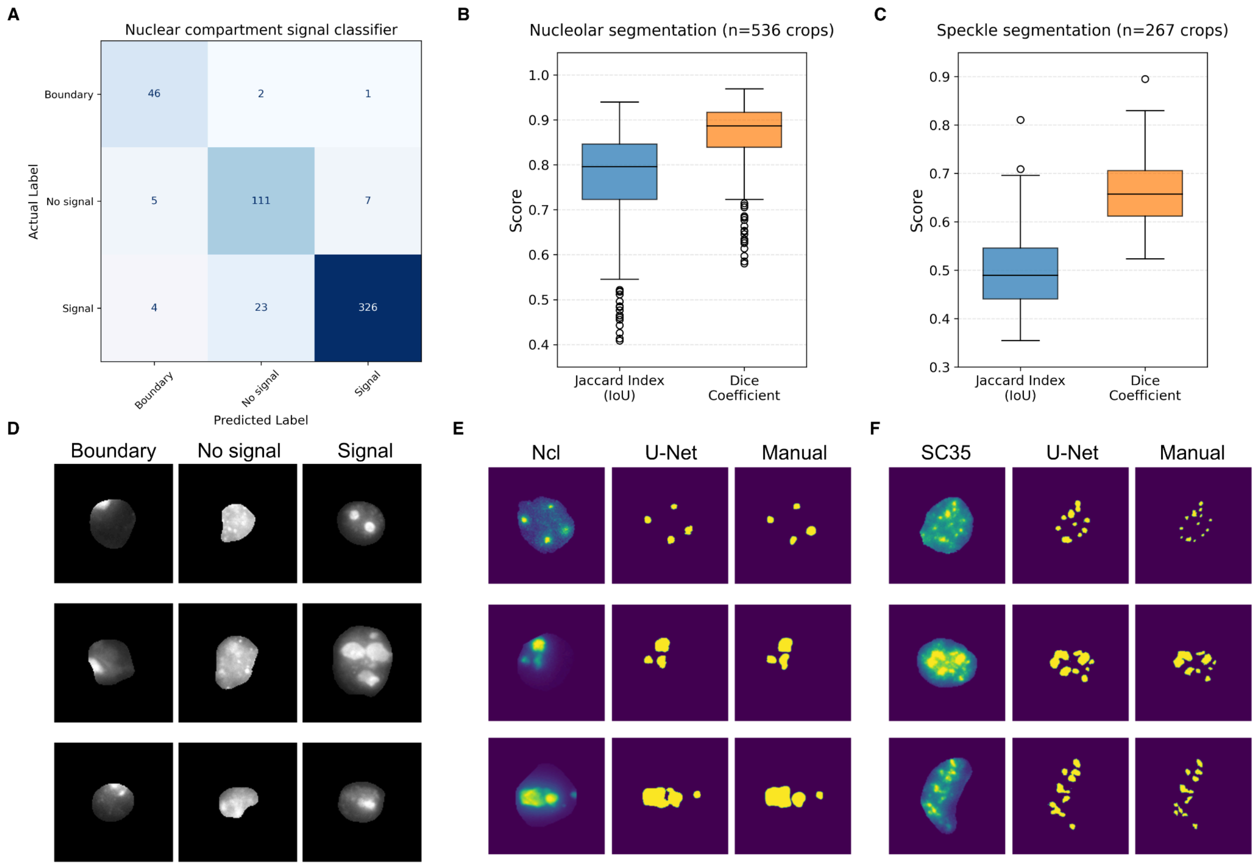


**Supplementary Figure 5. Performance of the nucleolar signal classifier and nuclear compartment segmentation models.** (A) Confusion matrix for the nucleolar signal classifier on the validation set. (B, C) Distributions of the Jaccard index and Dice coefficient for the segmentation of manually annotated nucleoli (B) and nuclear speckles (C). (D) Representative image crops corresponding to each signal classifier class. (E,F) Comparison between original crops, U-Net segmentations and manual annotations for Ncl (E) and SC35 (F) stains.

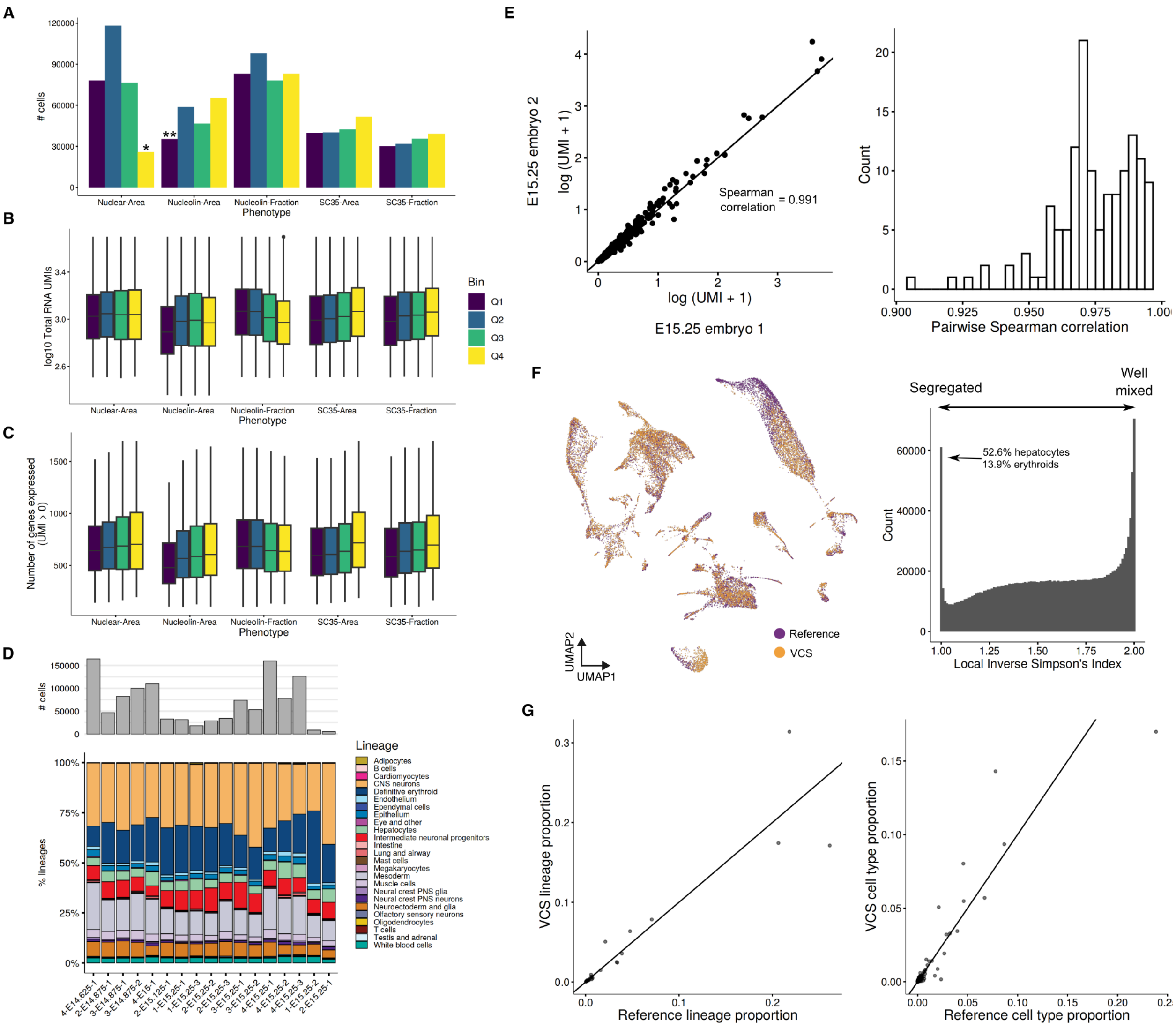


**Supplementary Figure 6. Quality control and reference comparison of E15 VCS-sci-RNA-seq3 data.** (A) Number of recovered nuclei, (B) total RNA UMIs, and (C) number of detected genes across experiments. *Reduced yield in the Q4 bin of the nuclear area experiment due to incomplete pelleting. **Reduced yield in the Q1 bin of one of the nucleolar area experiments due to stream misalignment. (D) Barplot of total cells recovered and stacked barplots of lineage proportions (bottom) for each embryo replicate. (E) Example scatterplot of expression patterns between embryo replicates E15.25-1 and E15.25-2 (left), and histogram of pairwise Spearman correlation coefficients across all replicates (right). (F) UMAP embedding showing the overlap between reference and VCS data (left), and the distribution of the local inverse Simpson's index (right), indicating high overlap with the least overlap observed in hepatocytes. (G) Concordance of lineage (left) and cell type (right) proportions between the reference and VCS-sci-RNA-seq3 data.

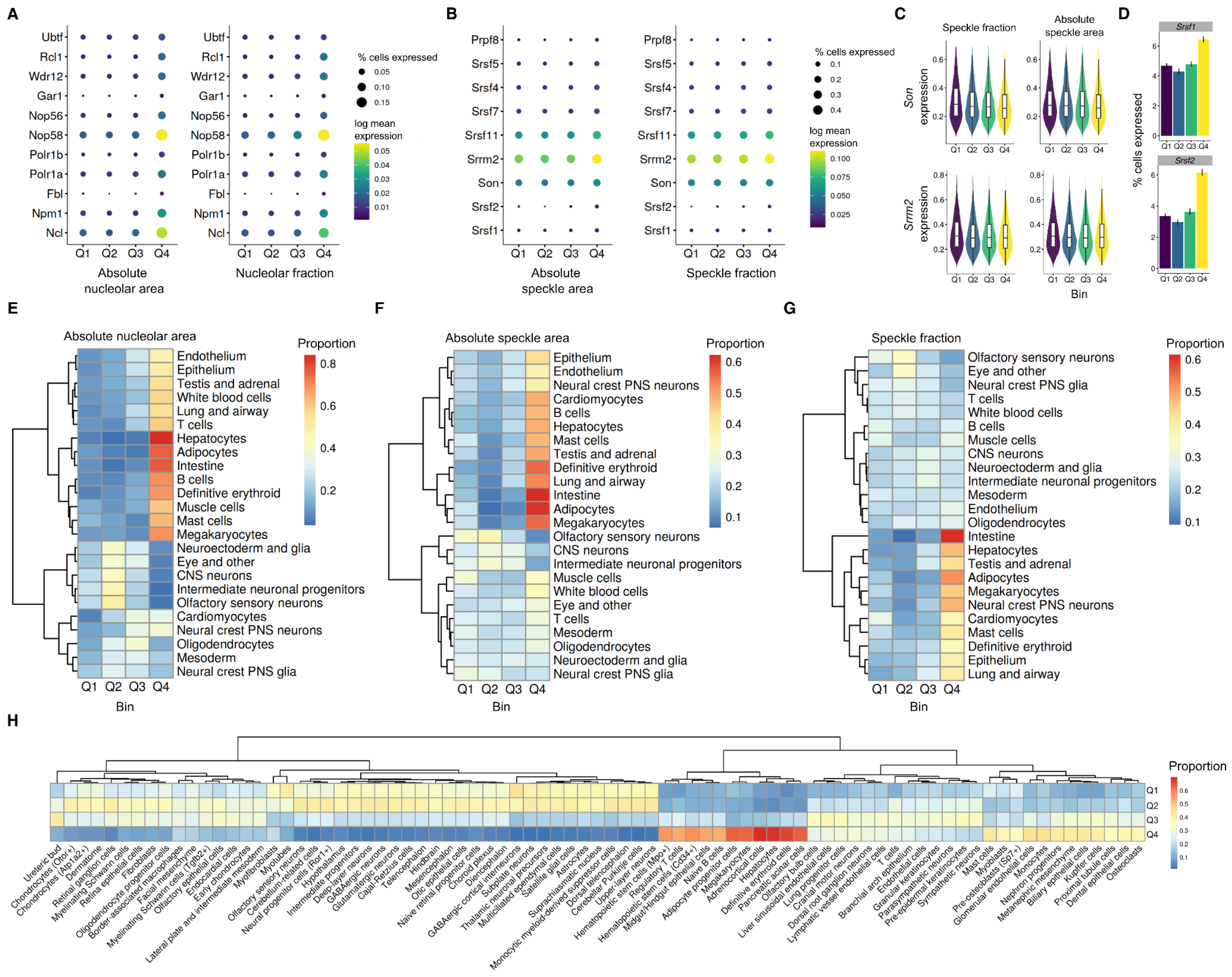


**Supplementary Figure 7. Comparison of VCS bins with nuclear compartment markers and lineage-specific distribution across bins.** (A, B) Expression dotplots of genes associated with the structure of the nucleolus (A) and nuclear speckles (B), across absolute area (left) and fraction (right) VCS bins. (C) Violin plots of *Son* and *Srrm2* expression grouped by absolute speckle size and fraction bins. (D) Barplots showing the percentage of cells with at least one UMI detected for *Srsf1* and *Srsf2*. (E-G) Heatmap showing the distribution of VCS bin labels for absolute nucleolar area (E), absolute speckle area (F), and speckle fraction (G) across major E15 cell lineages. (H) Heatmap of the nucleolar fraction bin distributions across granular cell type annotations.

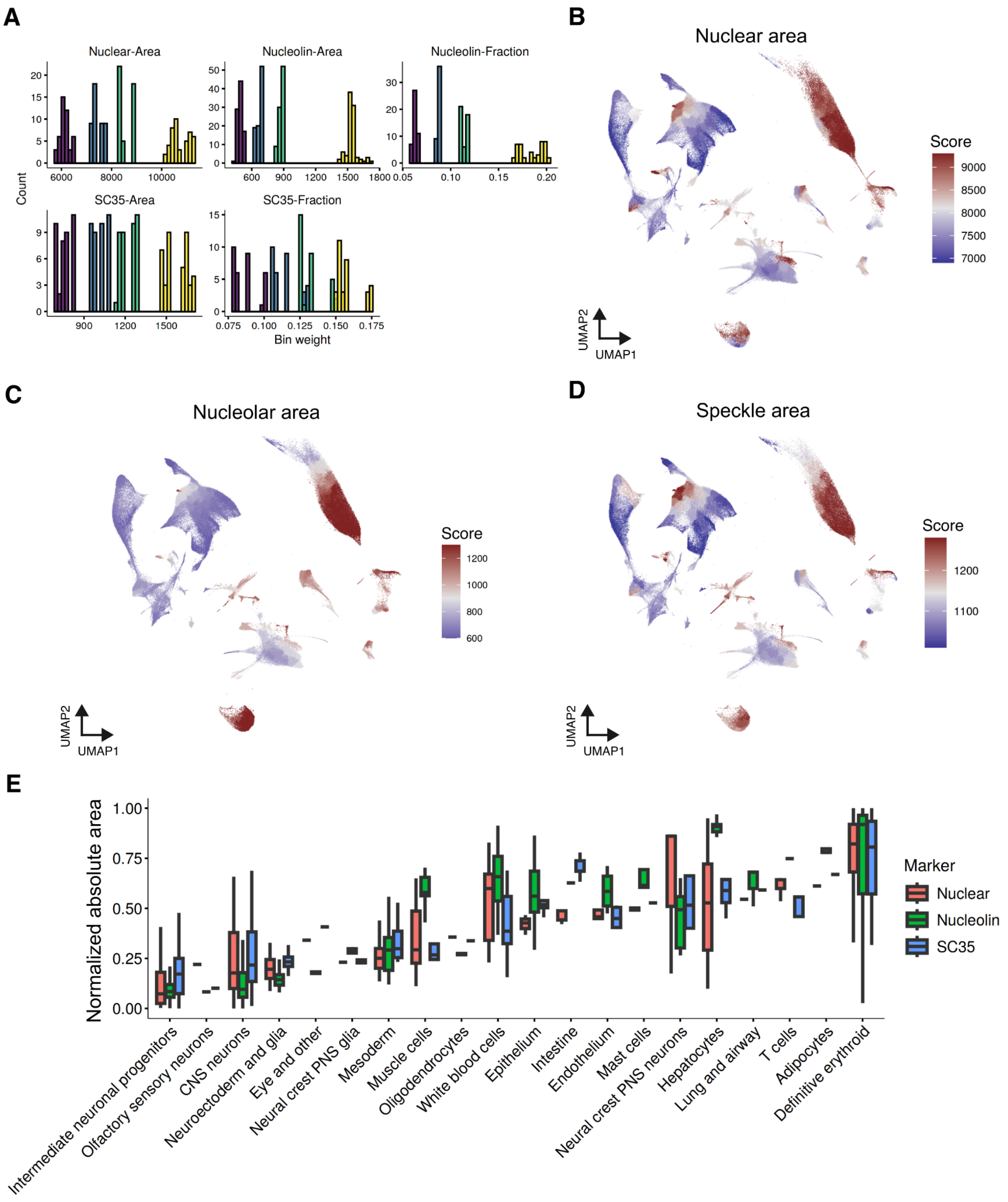


**Supplementary Figure 8. Distribution of the VCS-derived nuclear compartment scores.** (A) Distribution of the bin weights used to calculate the cluster-level weighted average scores across experiments. (B-D) UMAP embeddings of E15 embryo data colored by the scores for nuclear area (B), absolute nucleolar area (C), and absolute speckle area (D). (E) Boxplots of the absolute nuclear compartment area scores across major cell lineages.

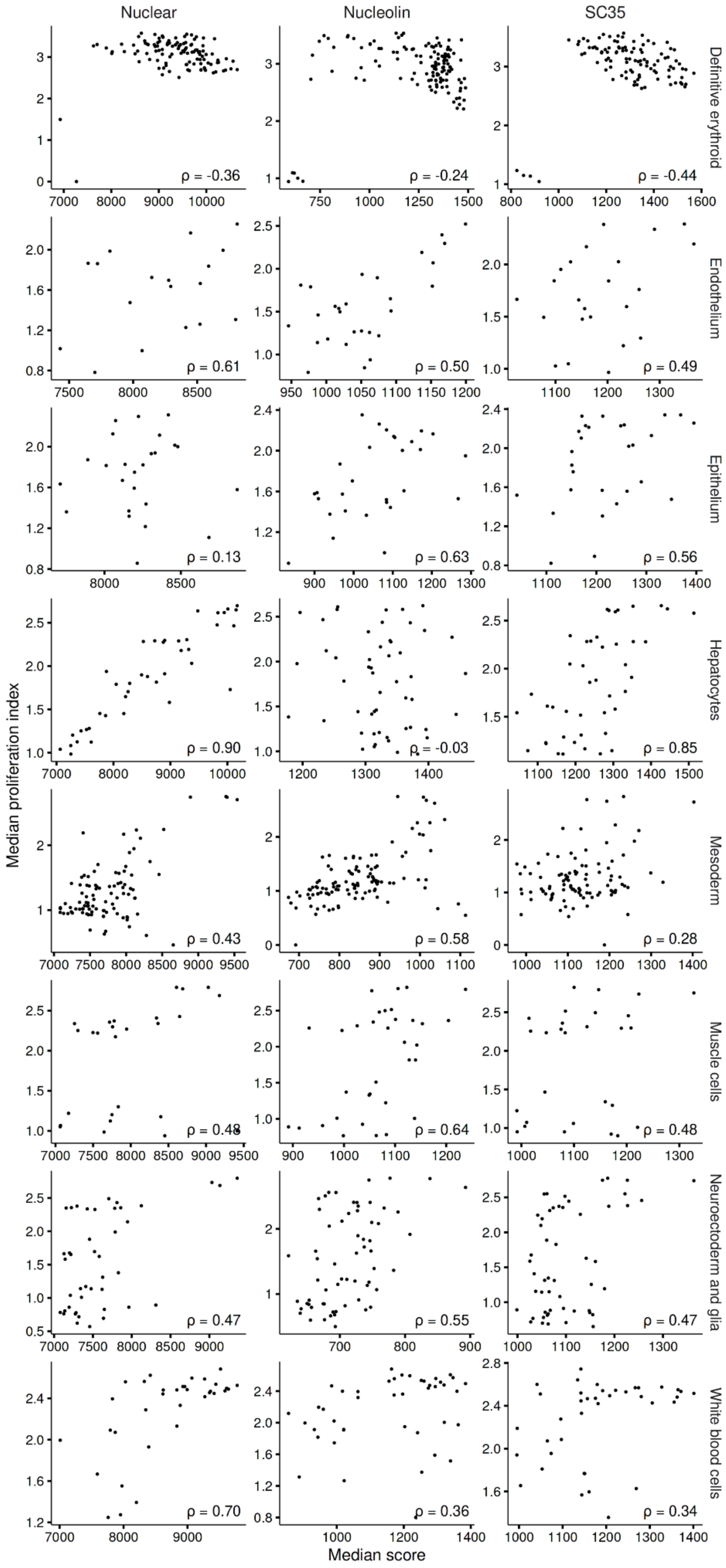

Nuclear
Nucleolin
SC35
Definitive erythroid
Endothelium
Epithelium
Hepatocytes
Mesoderm
Muscle cells
Neuroectoderm and glia
White blood cells
ρ = -0.36
ρ = -0.24
ρ = -0.44
ρ = 0.61
ρ = 0.50
ρ = 0.49
ρ = 0.13
ρ = 0.63
ρ = 0.56
ρ = 0.90
ρ = -0.03
ρ = 0.85
ρ = 0.43
ρ = 0.58
ρ = 0.28
ρ = 0.48
ρ = 0.64
ρ = 0.48
ρ = 0.47
ρ = 0.55
ρ = 0.47
ρ = 0.70
ρ = 0.36
ρ = 0.34
Median proliferation index
Median score

**Supplementary Figure 9. Lineage-specific comparison of the absolute size scores against proliferation index.** Scatterplots of the absolute size scores vs the proliferation index, grouped by major cell lineage.

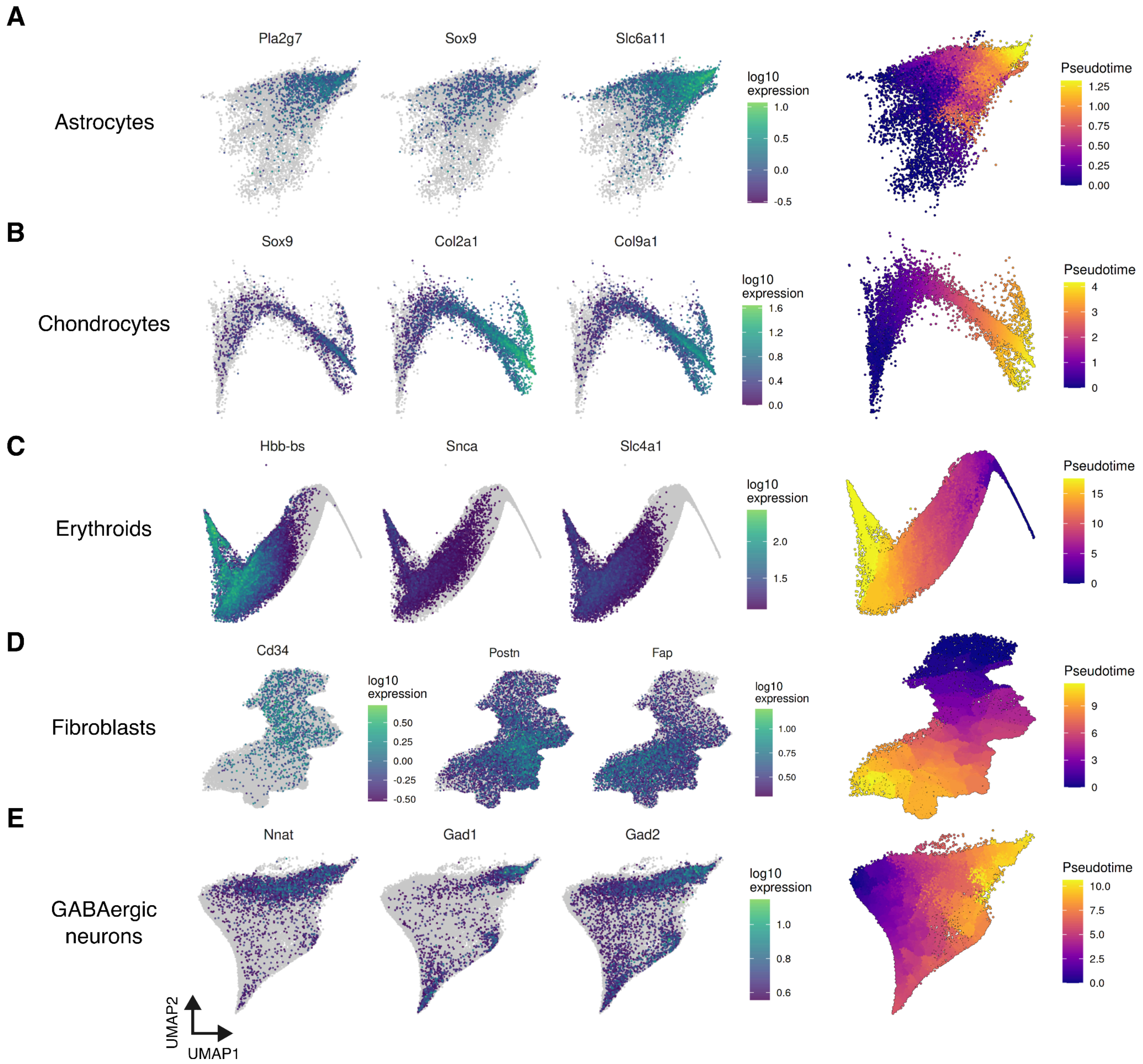


**Supplementary Figure 10. Marker gene expression and pseudotime trajectory of E15 embryonic lineages (from astrocytes to GABAergic neurons).** (A-E) Visualization of marker gene expression and corresponding pseudotime trajectories for astrocytes (A), chondrocytes (B), erythroids (C), fibroblasts (D), and GABAergic neurons (E).

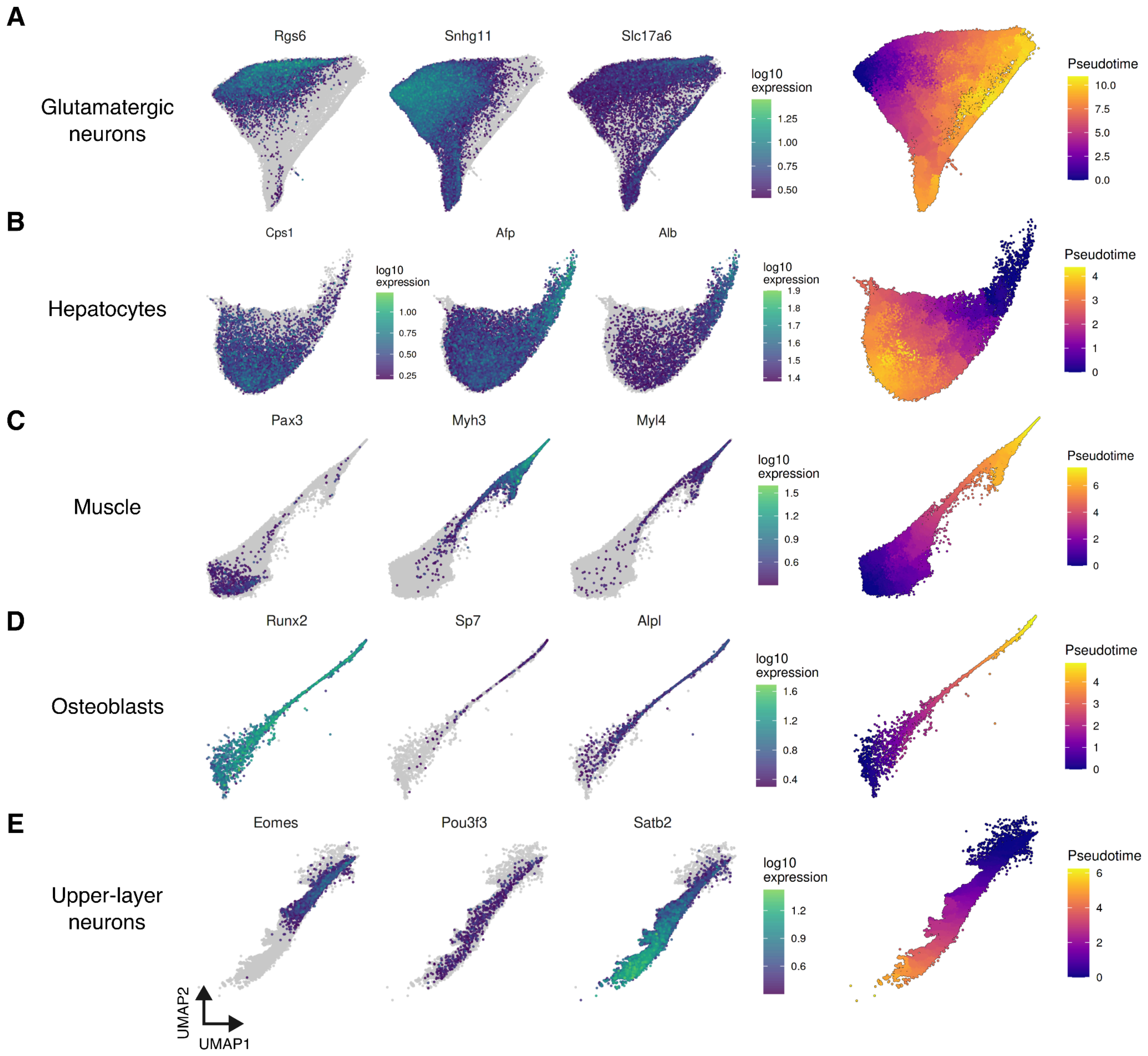


**Supplementary Figure 11. Marker gene expression and pseudotime trajectory of E15 embryonic lineages (from glutamatergic to upper-layer neurons).** (A-E) Visualization of marker gene expression and corresponding pseudotime trajectories for glutamatergic neurons (A), hepatocytes (B), muscle (C), osteoblasts (D), and upper-layer neurons (E).

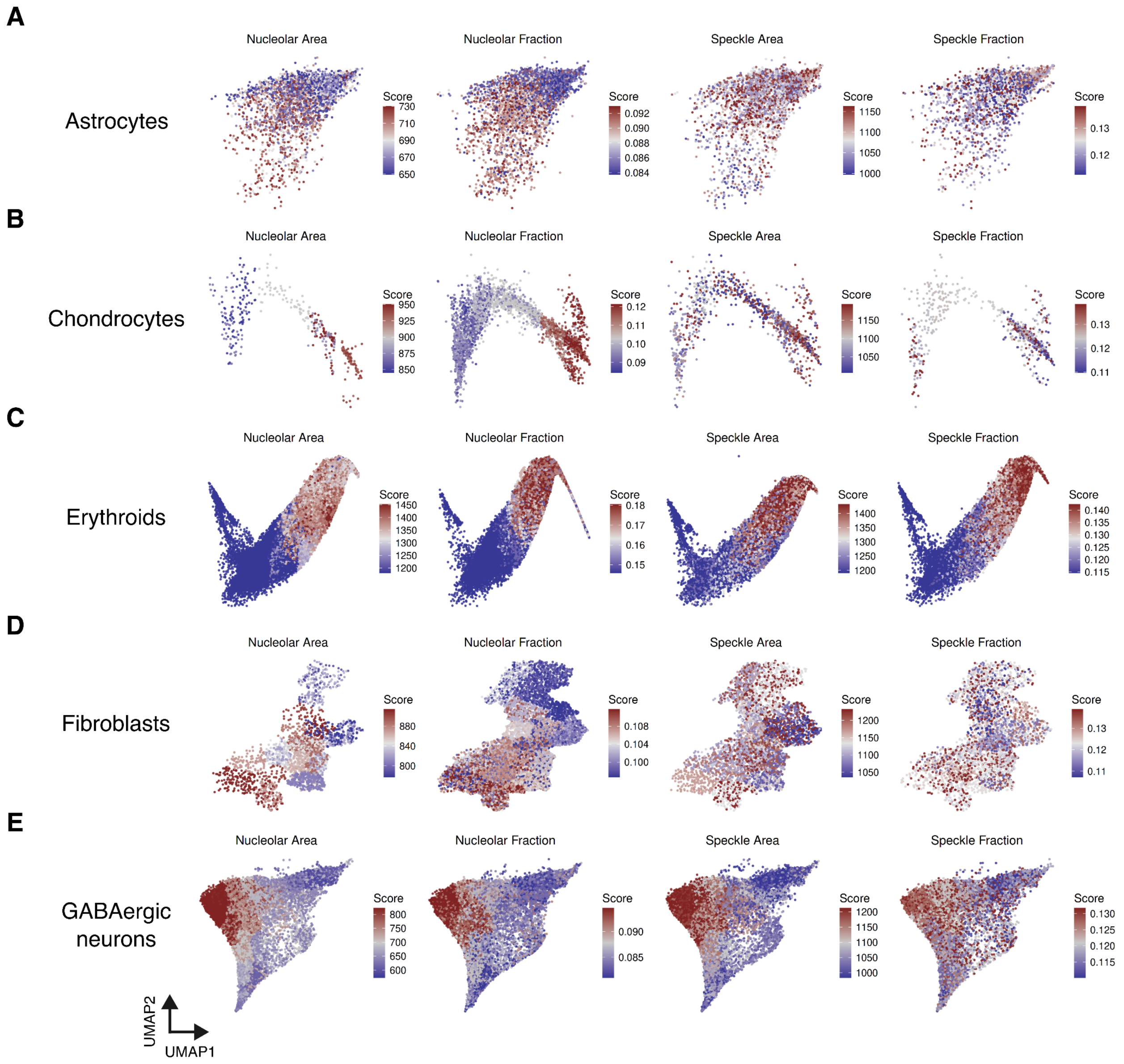


**Supplementary Figure 12. Mapping of the nuclear compartment scores across E15 embryonic lineages (from astrocytes to GABAergic neurons).** (A-E) Visualization of the nuclear compartment size phenotypes for (A) astrocytes, (B) chondrocytes, (C) erythroids, (D) fibroblasts, and (E) GABAergic neurons.

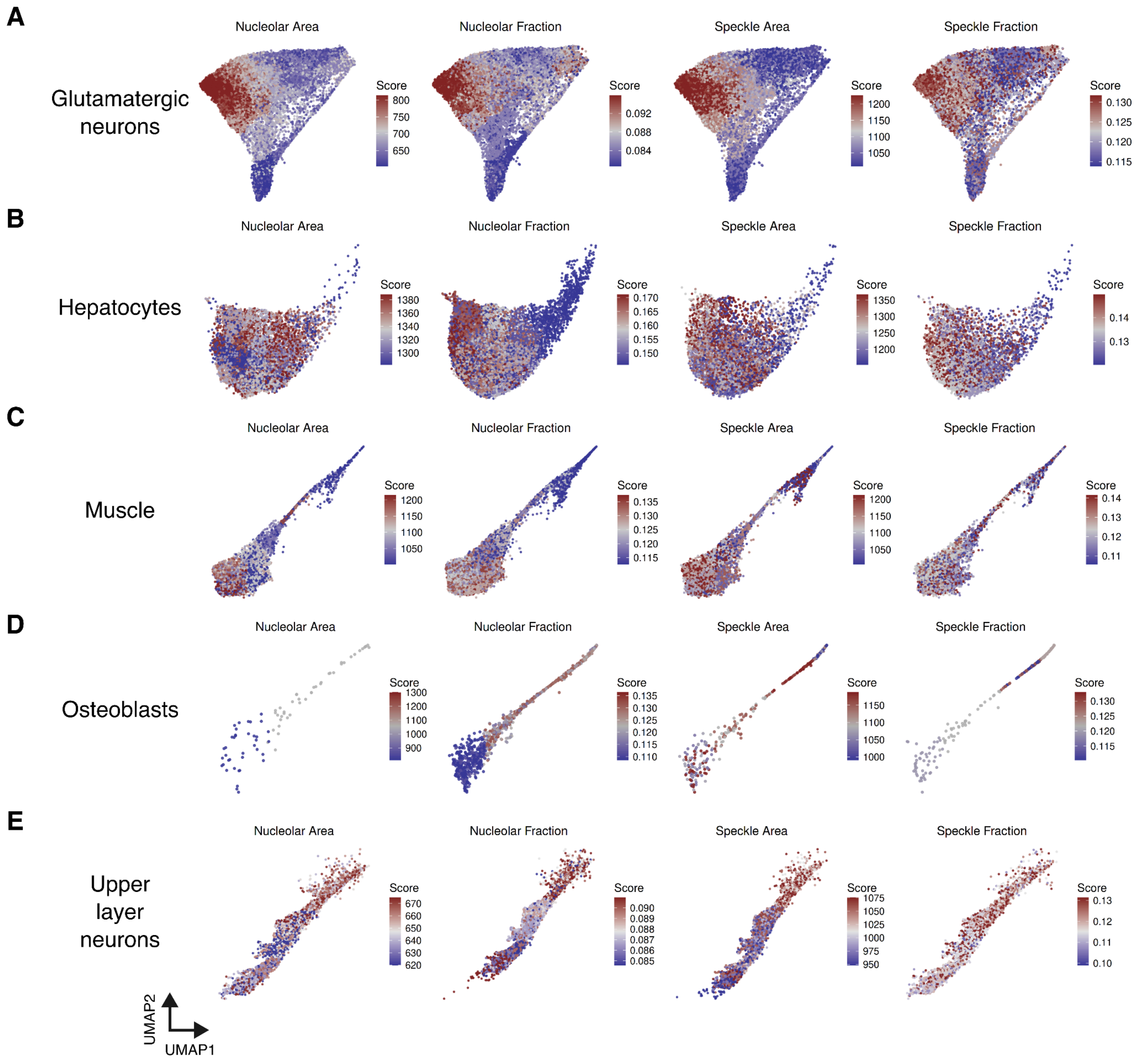


**Supplementary Figure 13. Mapping of the nuclear compartment scores across E15 embryonic lineages (from glutamatergic to upper-layer neurons).** (A-E) Visualization of the nuclear compartment size phenotypes for (A) glutamatergic neurons, (B) hepatocytes, (C) muscle, (D) osteoblasts, and (E) upper-layer neurons.

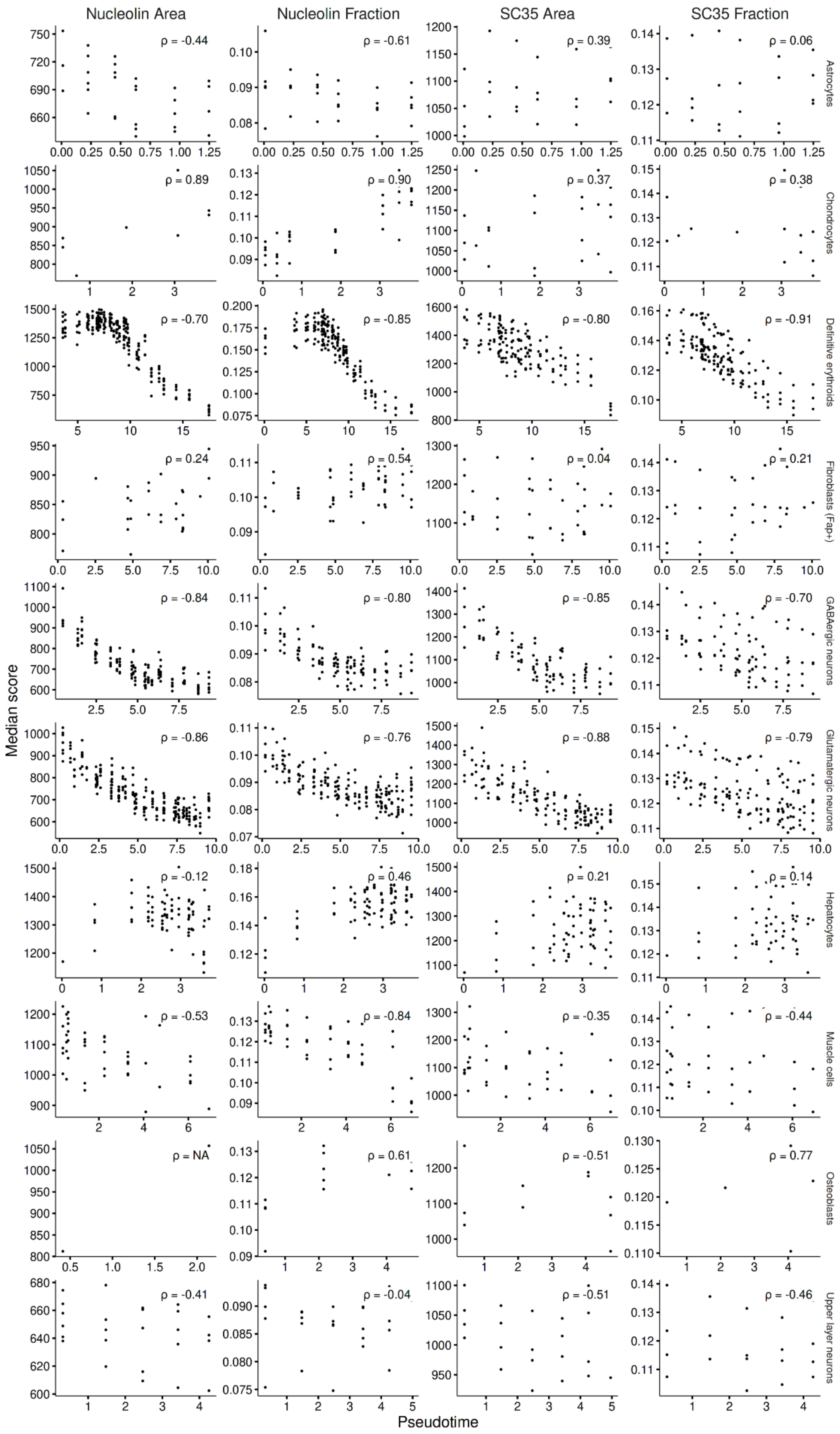

Nucleolin Area
Nucleolin Fraction
SC35 Area
SC35 Fraction
Astrocytes
ρ = -0.44
ρ = -0.61
ρ = 0.39
ρ = 0.06
Chondrocytes
ρ = 0.89
ρ = 0.90
ρ = 0.37
ρ = 0.38
Definitive erythroids
ρ = -0.70
ρ = -0.85
ρ = -0.80
ρ = -0.91
Fibroblasts (Fap+)
ρ = 0.24
ρ = 0.54
ρ = 0.04
ρ = 0.21
GABAergic neurons
ρ = -0.84
ρ = -0.80
ρ = -0.85
ρ = -0.70
Glutamatergic neurons
ρ = -0.86
ρ = -0.76
ρ = -0.88
ρ = -0.79
Hepatocytes
ρ = -0.12
ρ = 0.46
ρ = 0.21
ρ = 0.14
Muscle cells
ρ = -0.53
ρ = -0.84
ρ = -0.35
ρ = -0.44
Osteoblasts
ρ = NA
ρ = 0.61
ρ = -0.51
ρ = 0.77
Upper layer neurons
ρ = -0.41
ρ = -0.04
ρ = -0.51
ρ = -0.46
Median score
Pseudotime

**Supplementary Figure 14. Lineage-specific comparison of the nuclear compartment size scores against differentiation.** Scatterplots of the nuclear compartment size scores vs pseudotime across E15 cell lineages.

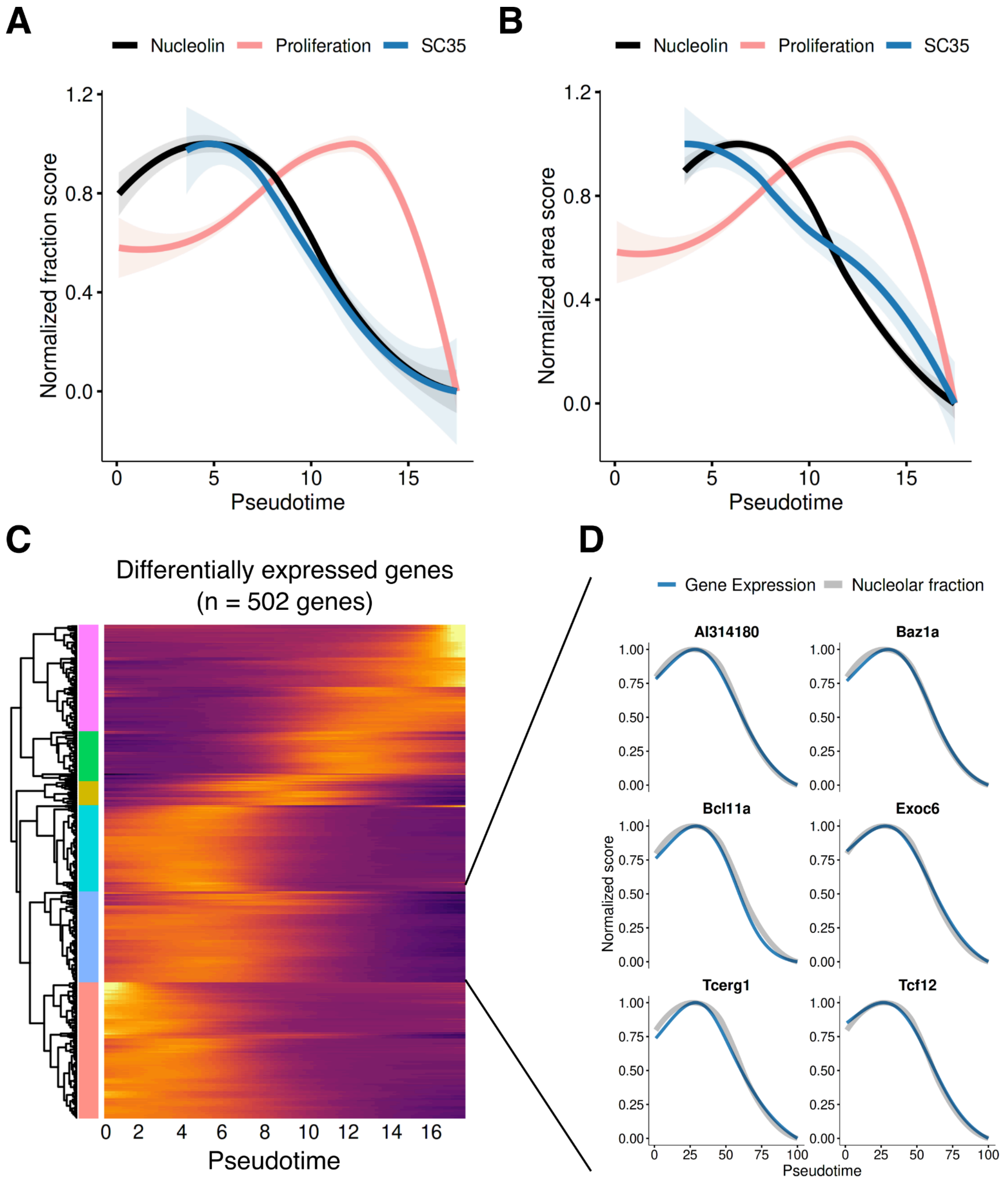


**Supplementary Figure 15. Dynamics of the nuclear compartment size during erythroid differentiation.** (A, B) Pseudotemporal changes in the nuclear compartment scores and the proliferation index for (A) normalized fraction scores and (B) normalized area scores. Curves are fitted using cubic spline regression. (C) Heatmap of differentially expressed genes modeled across pseudotime. Genes are hierarchically clustered, and the blue cluster represents expression patterns highly concordant with the nuclear compartment size scores. (D) Expression profiles of the six top ranked genes (blue) and the nucleolar fraction score (grey) along pseudotime.

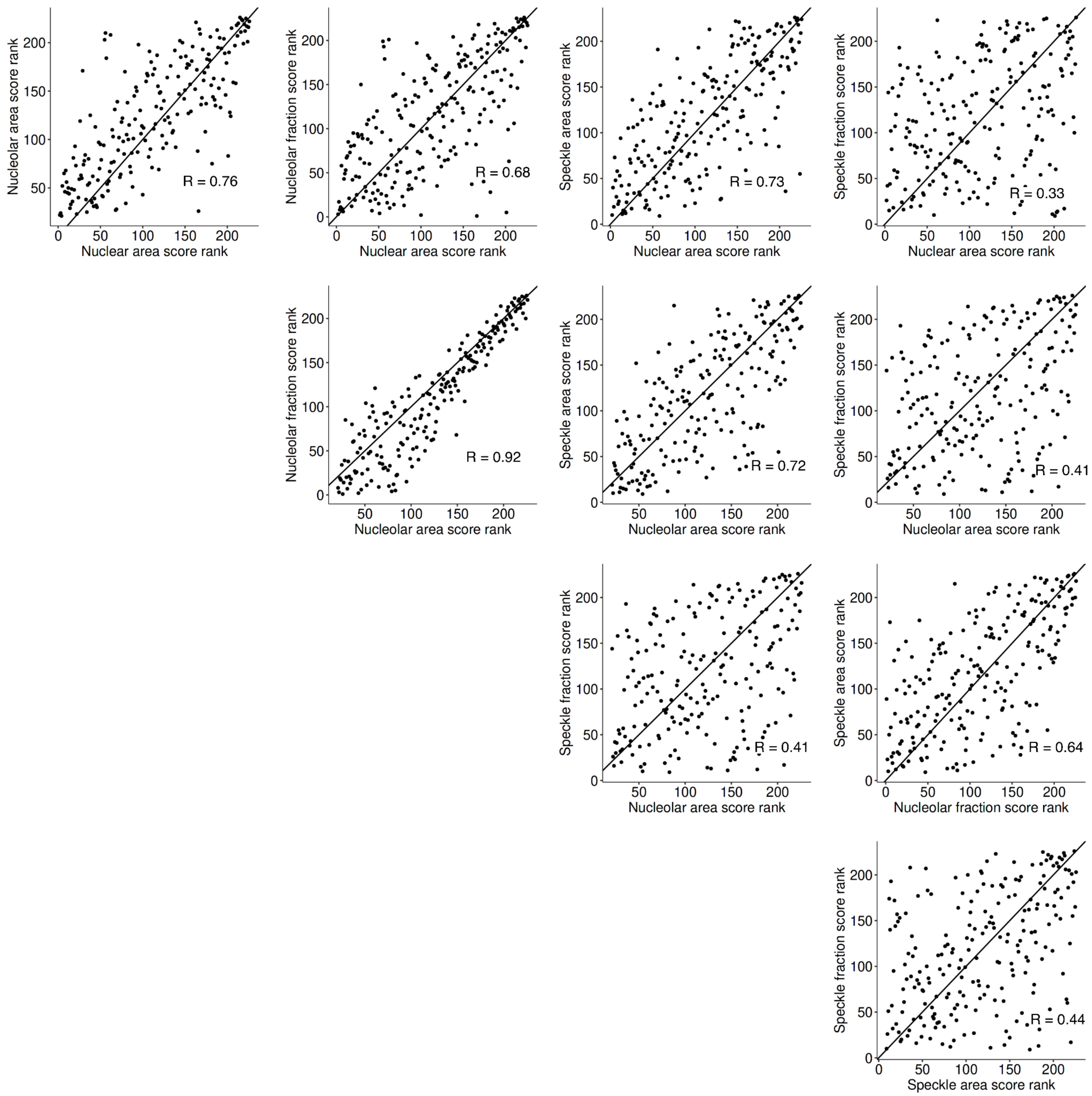


**Supplementary Figure 16. Pairwise correlations of the nuclear compartment size scores.** Scatterplots showing the relationships between all measured compartment size scores. Diagonal lines indicate the y = x, and the Spearman correlation coefficients are indicated within each panel.

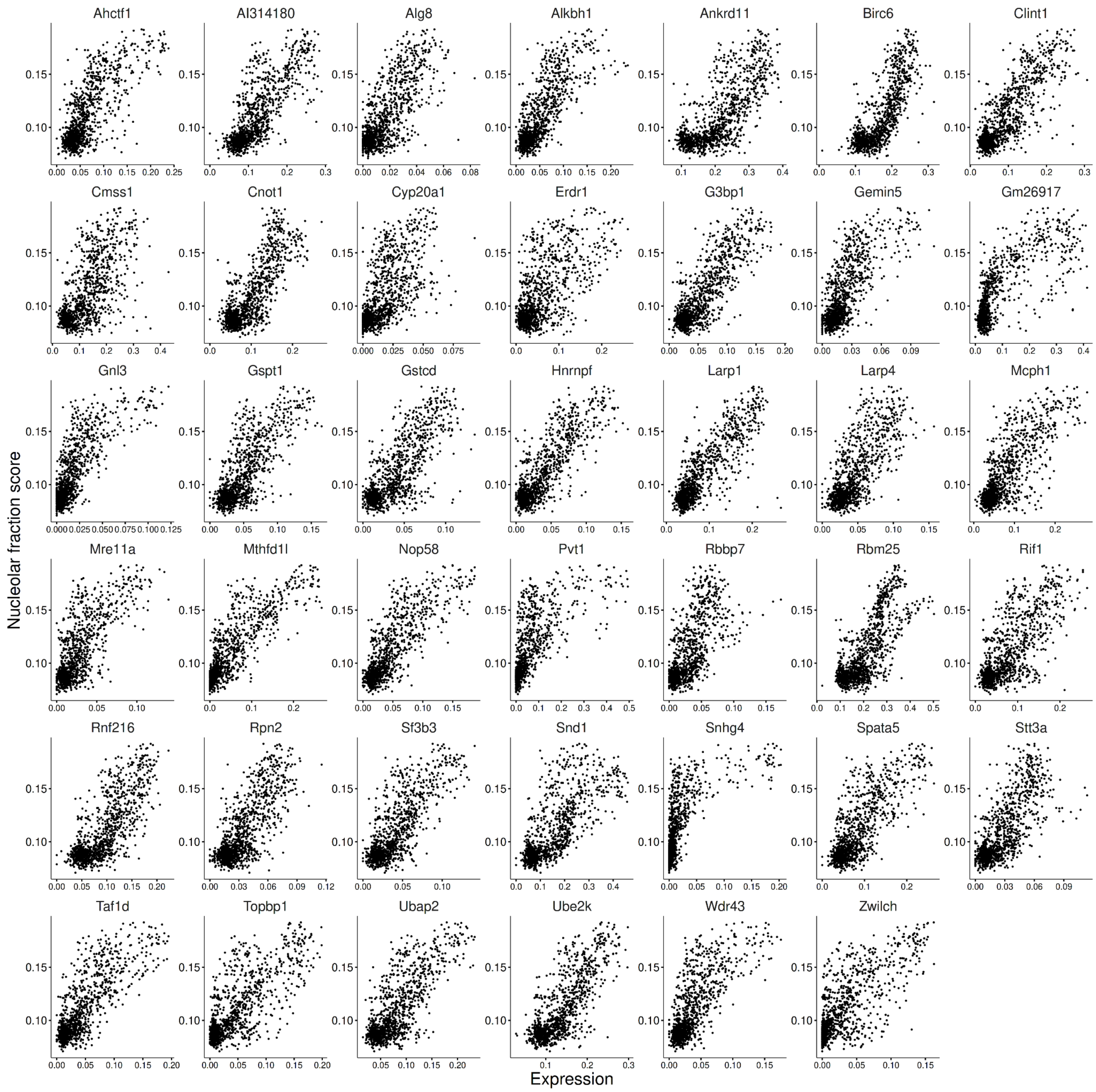


**Supplementary Figure 17. Relationship between gene expression and the nucleolar fraction score for the 41 candidate genes.** Faceted scatterplots showing the relationship between gene expression and the nucleolar fraction score for the 41 most strongly associated genes.

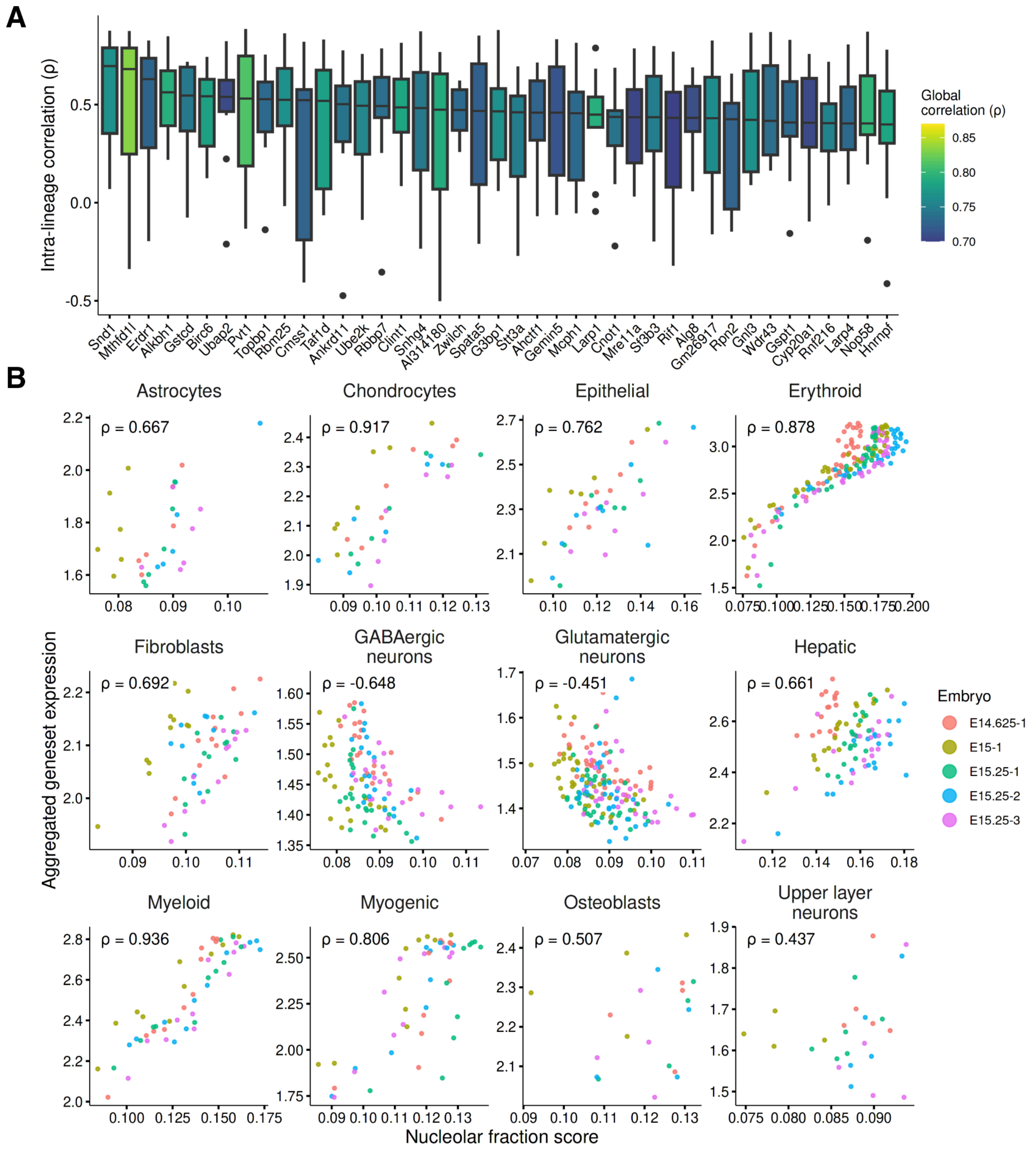


**Supplementary Figure 18. Intra-lineage correlation between the nucleolar fraction score and the 41 candidate genes.** (A) Boxplots of intra-lineage correlation coefficients for the top 41 candidate genes, colored by global rank correlation. (B) Comparison of the nucleolar fraction score and aggregate expression of the 41 candidate genes across the lineages and trajectory groups, colored by embryo replicate.

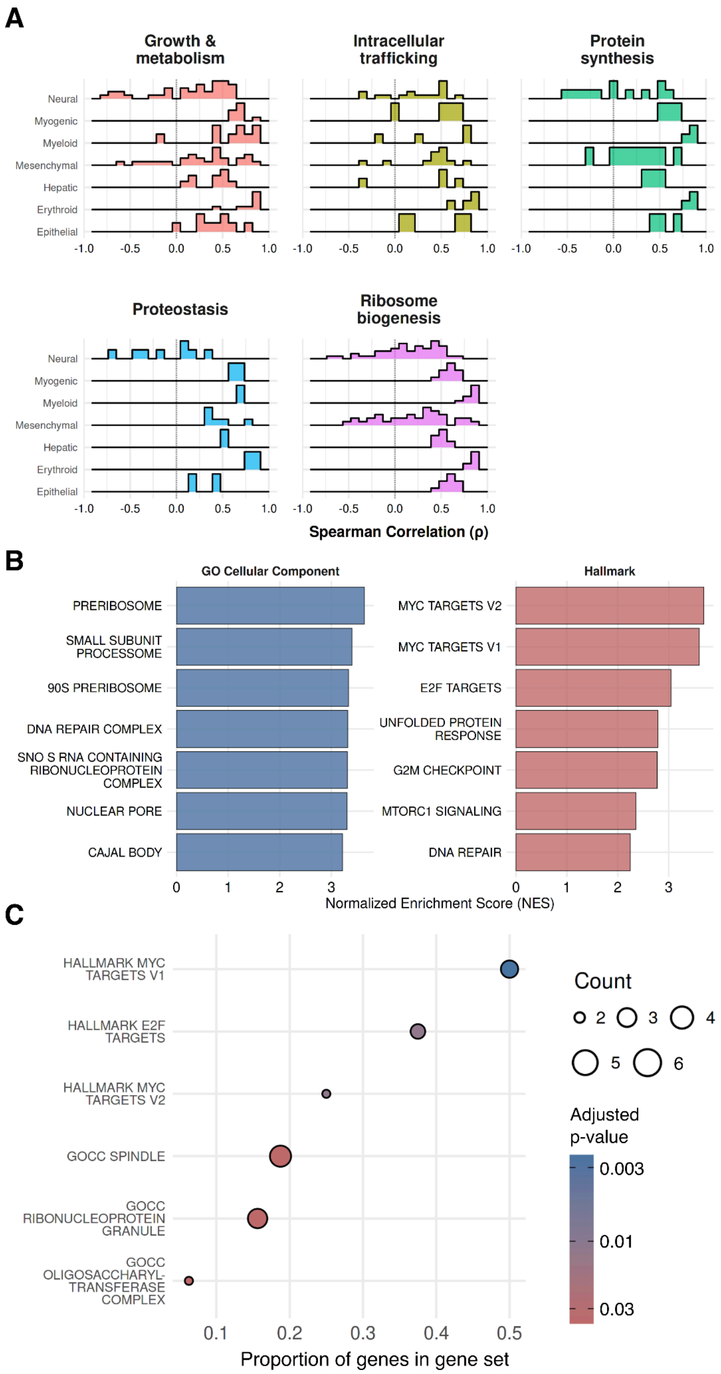


**Supplementary Figure 19. Functional enrichment analysis of the 41-gene nucleolar signature.** (A) Distribution of intra-lineage correlations between the nucleolar fraction score and established gene sets (MSigDB Hallmark, GOCC and KEGG), categorized into five broad biological functions. (B) Gene set enrichment analysis of the full correlation-rank gene list against GOCC (left) and Hallmark (right) gene sets. (C) Over-representation analysis of the top 41 candidate genes. The x-axis indicates the proportion of genes overlapping with each set, with significance colored by adjusted p-value.

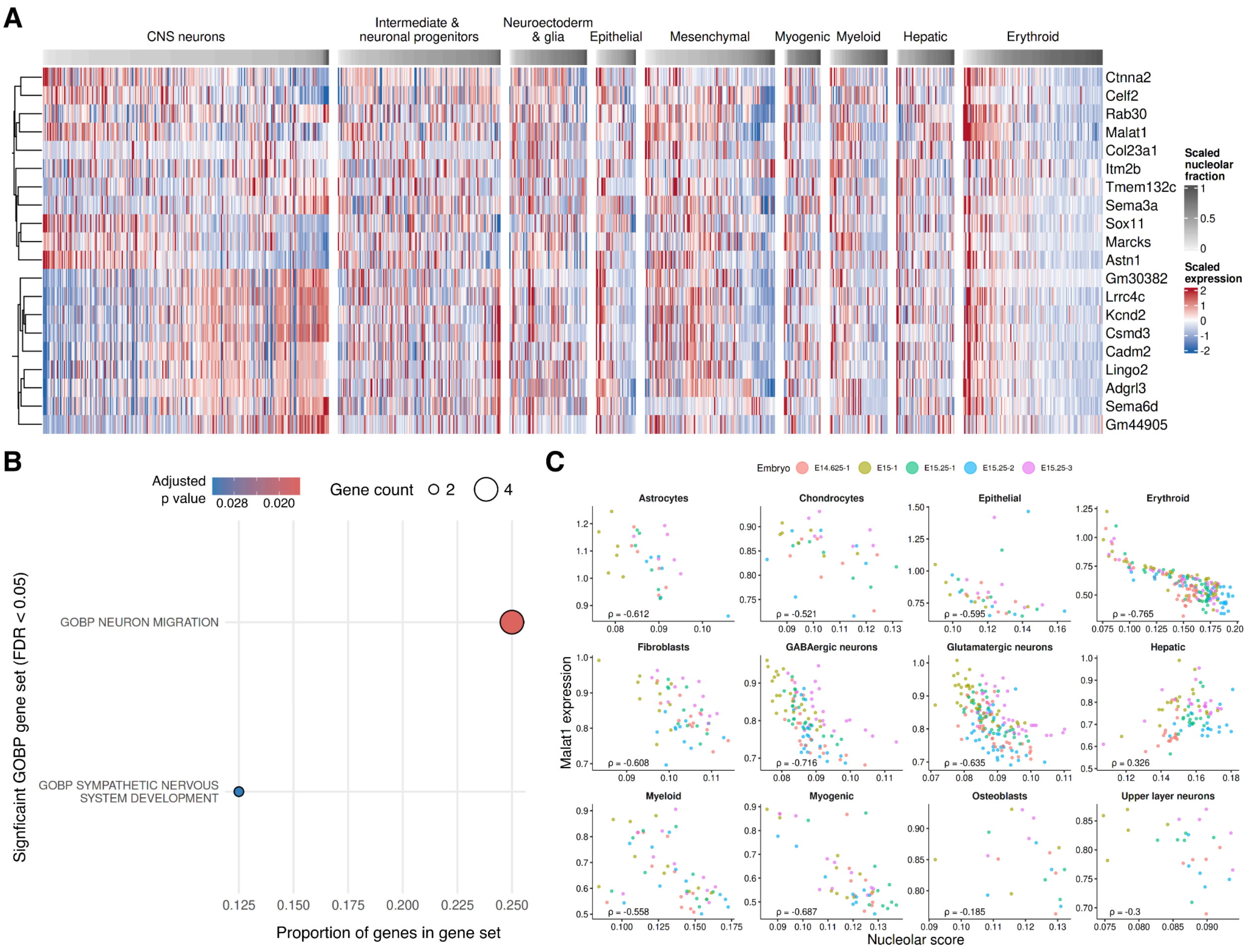


**Supplementary Figure 20. Genes negatively correlated with the nucleolar fraction score.** (A) Heatmap displaying scaled expression values of 20 negatively correlated genes with nucleolar fraction (intra-lineage correlation < -0.4 in at least 6 lineages and combined Stouffer's Z < -5) across major lineages. (B) Over-representation analysis of the 20 negatively correlated genes. (C) Comparison of *Malat1* expression and the nucleolar fraction score across lineages and trajectory groups, with points colored by embryo replicate. Negative correlation is observed across all groups except hepatocytes.

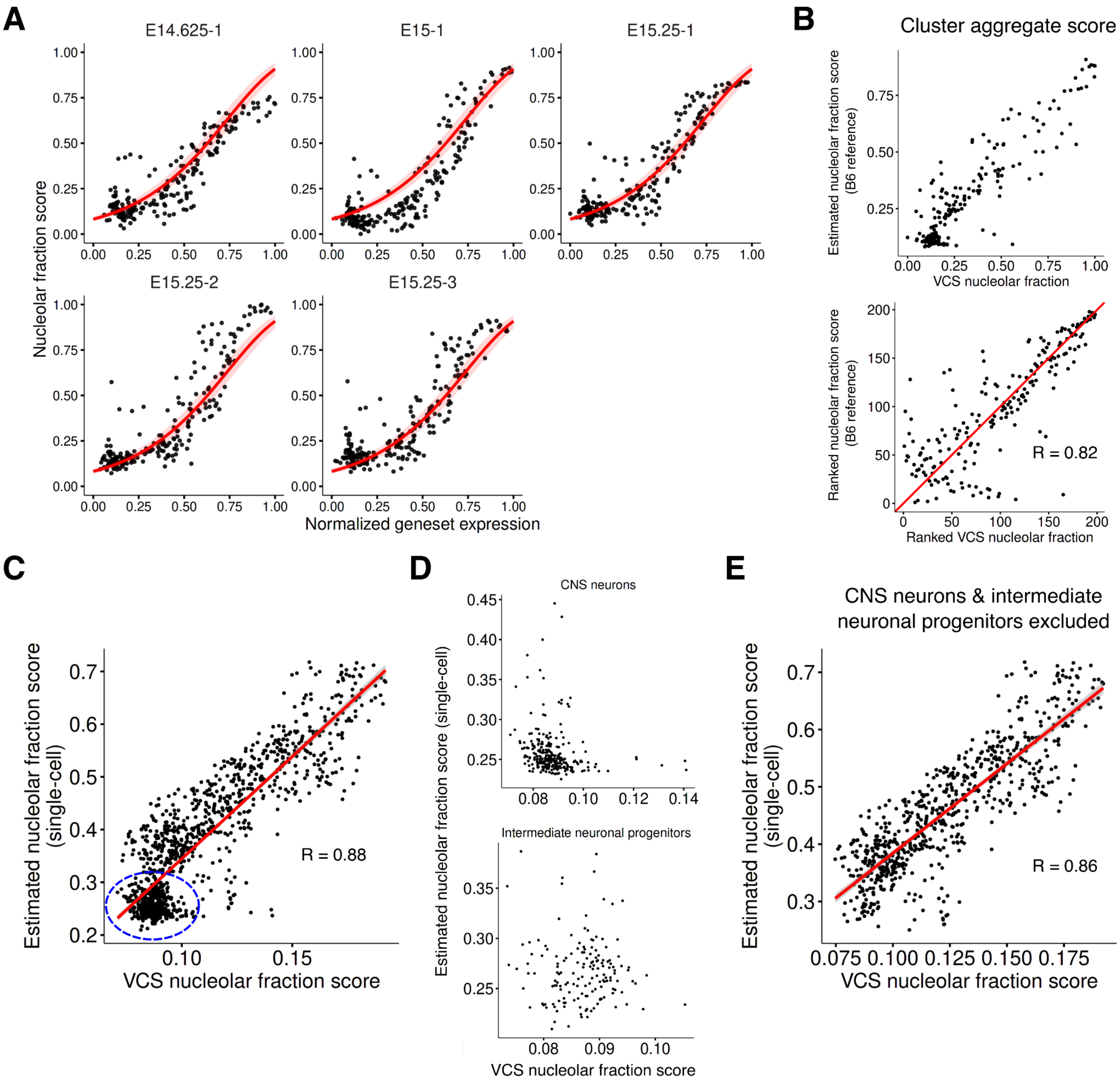


**Supplementary Figure 21. Concordance between the estimated and measured nucleolar fraction score.** (A) Relationship between the normalized aggregate expression of the nucleolar genes vs the measured nucleolar fraction score, faceted by embryo replicate and fitted with a general linear mixed model (red curves). (B) Pseudobulk-level comparison of the measured VCS nucleolar fraction against the estimated score derived from co-embedded atlas and VCS clusters, showing both raw values (top) and ranks (bottom). (C - E) Cluster level comparisons of the measured VCS nucleolar fraction against the median estimated nucleolar fraction score (calculated by applying the model to the VCS data at single-cell resolution) across (C) all clusters, (D) CNS neurons and intermediate neuronal progenitors (blue circle from C), and (E) all clusters excluding CNS neurons and intermediate neuronal progenitors.

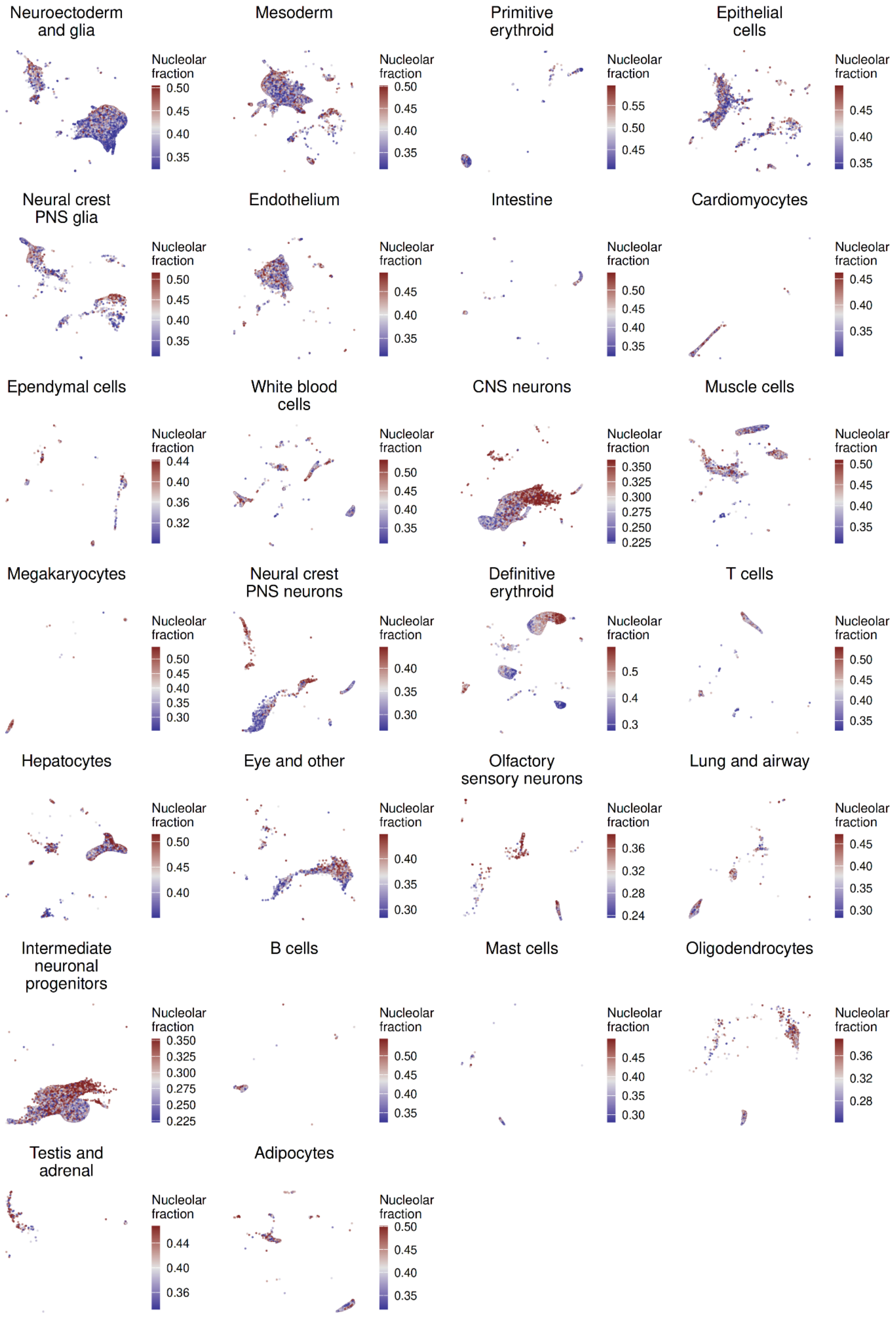


**Supplementary Figure 22. Mapping of the nucleolar fraction score across the major lineages in the mouse embryogenesis atlas.** UMAP embeddings of the mouse embryogenesis atlas faceted by major trajectory groups, with individual cells colored by the estimated nucleolar fraction score.

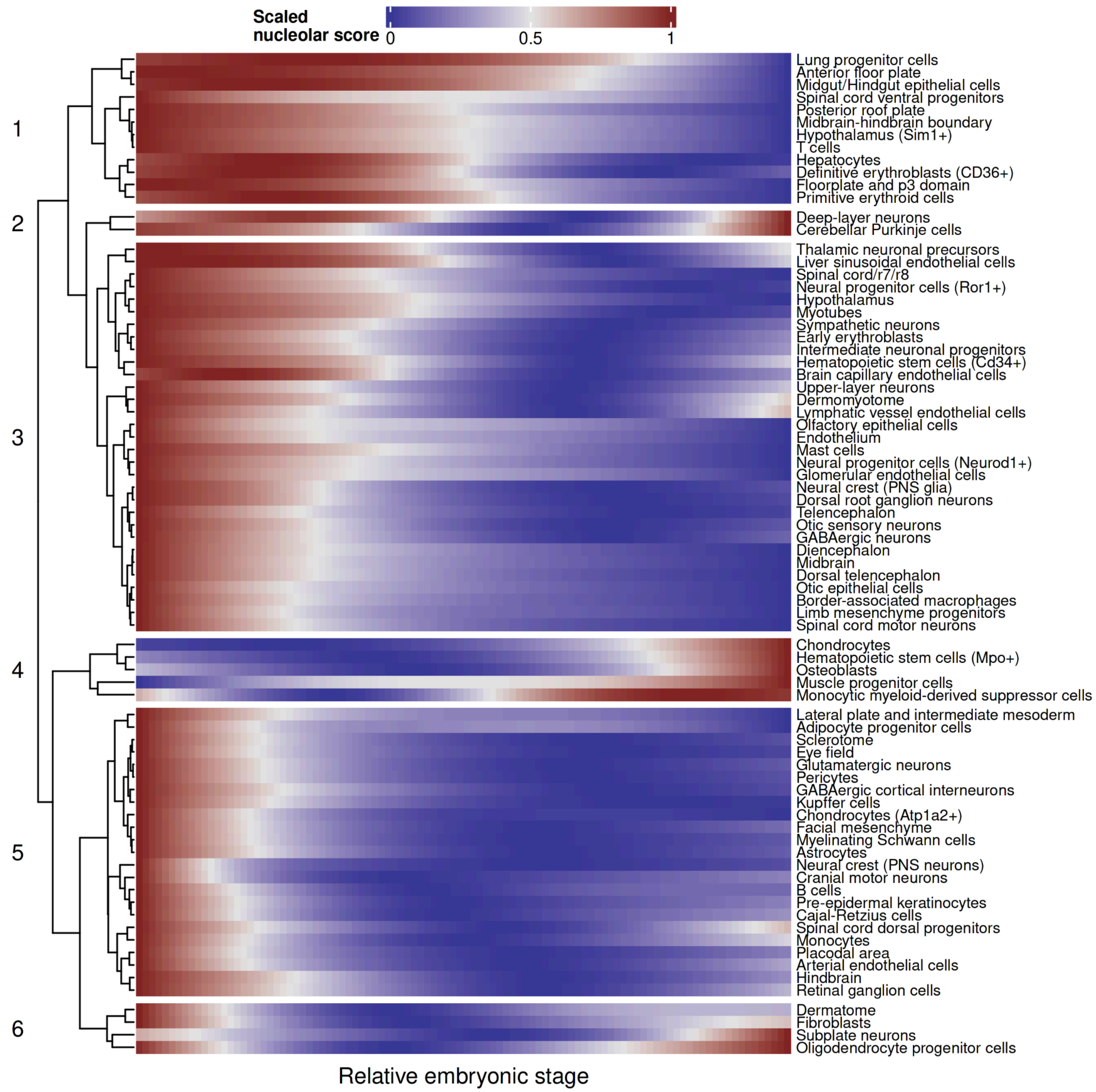


**Supplementary Figure 23. Dynamics of the estimated nucleolar fraction score across mouse embryonic development.** Heatmap of row-normalized, spline-smoothed nucleolar fraction score by cell type across developmental time. Analysis was restricted to cell types with at least 10 time points and 300 cells per timepoint. Only cell types demonstrating a significant trend (F-test vs a null intercept model) are shown. Rows are ordered by hierarchical clustering, with numbers on the left denoting the major identified clusters.

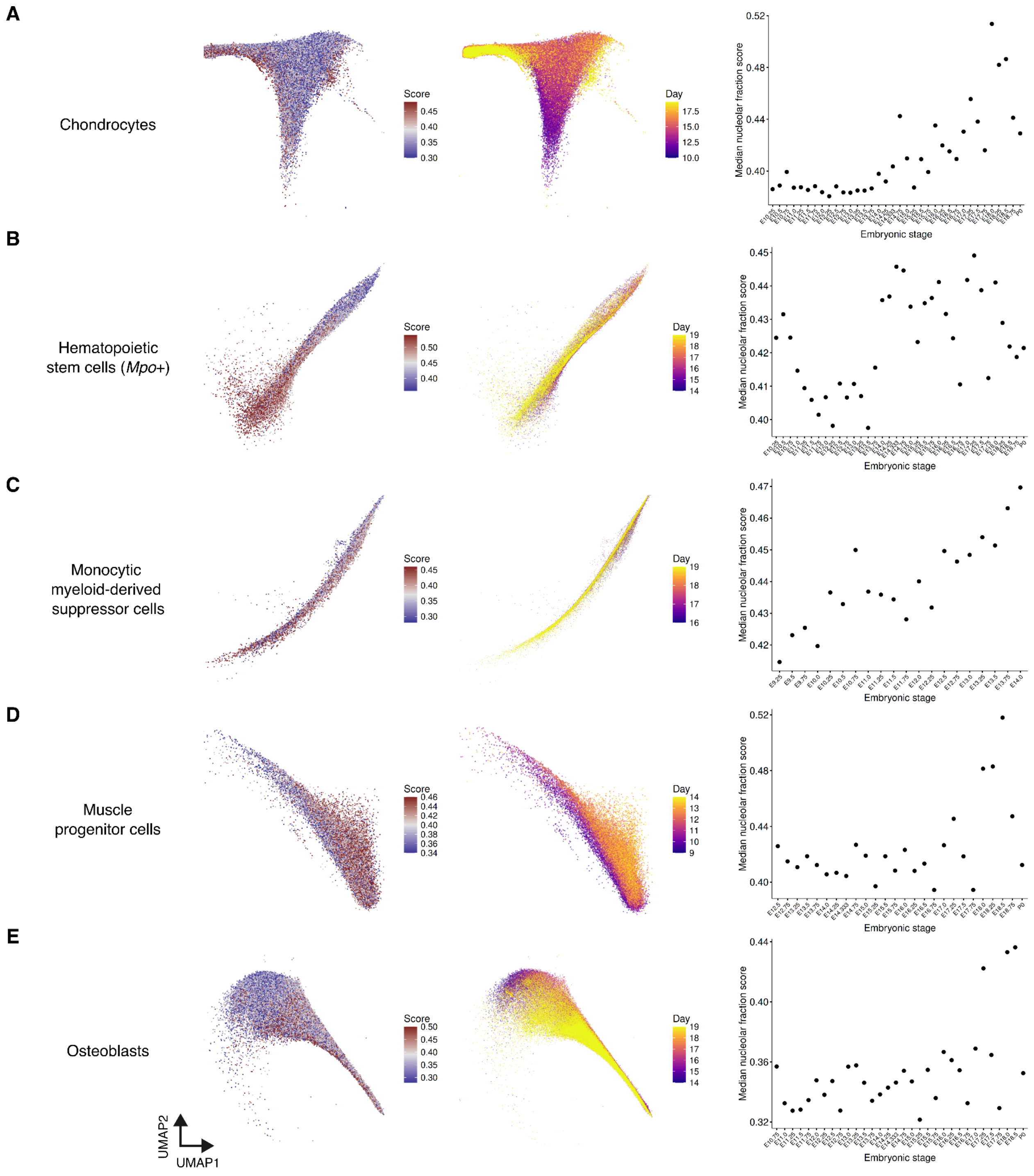


**Supplementary Figure 24. Cell types positively correlated between the estimated nucleolar fraction score and developmental time.** (A-E) UMAP embeddings of (A) chondrocytes, (B) *Mpo*-positive hematopoietic stem cells, (C) monocytic myeloid-derived suppressor cells, (D) muscle progenitor cells, and (E) osteoblasts, colored by the estimated nucleolar fraction score (left) and embryonic stage (middle). The right scatterplots show the increase in the median nucleolar fraction score over developmental time.

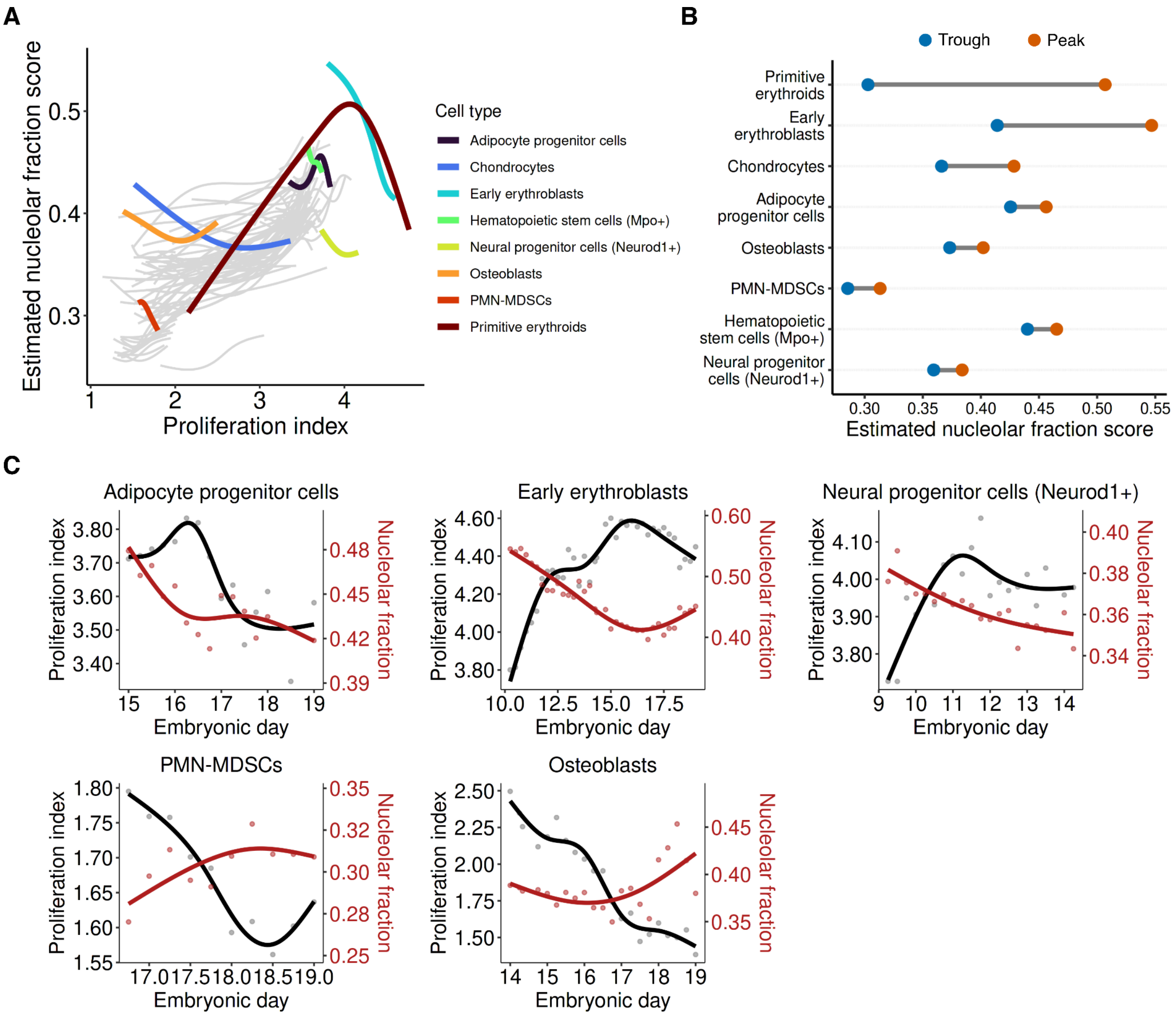


**Supplementary Figure 25. Cell types negatively correlated with the estimated nucleolar fraction score and proliferation index.** (A) Cubic spline regression models of the estimated nucleolar fraction score vs proliferation index across all cell types in the mouse embryogenesis atlas (grey). Cell types displaying a significant downward segment (segment Spearman correlation < -0.3 and minimum 5% score drop) are highlighted. (B) Dumbbell plots showing the difference in the maximum (peak) and minimum (trough) the estimated nucleolar fraction score within the negatively correlated segment for each identified lineage. (C) Dual-axis scatterplots quantifying the temporal changes in proliferation (black) and the estimated nucleolar fraction score (red) for the remaining five identified cell types not shown in Figure 6.

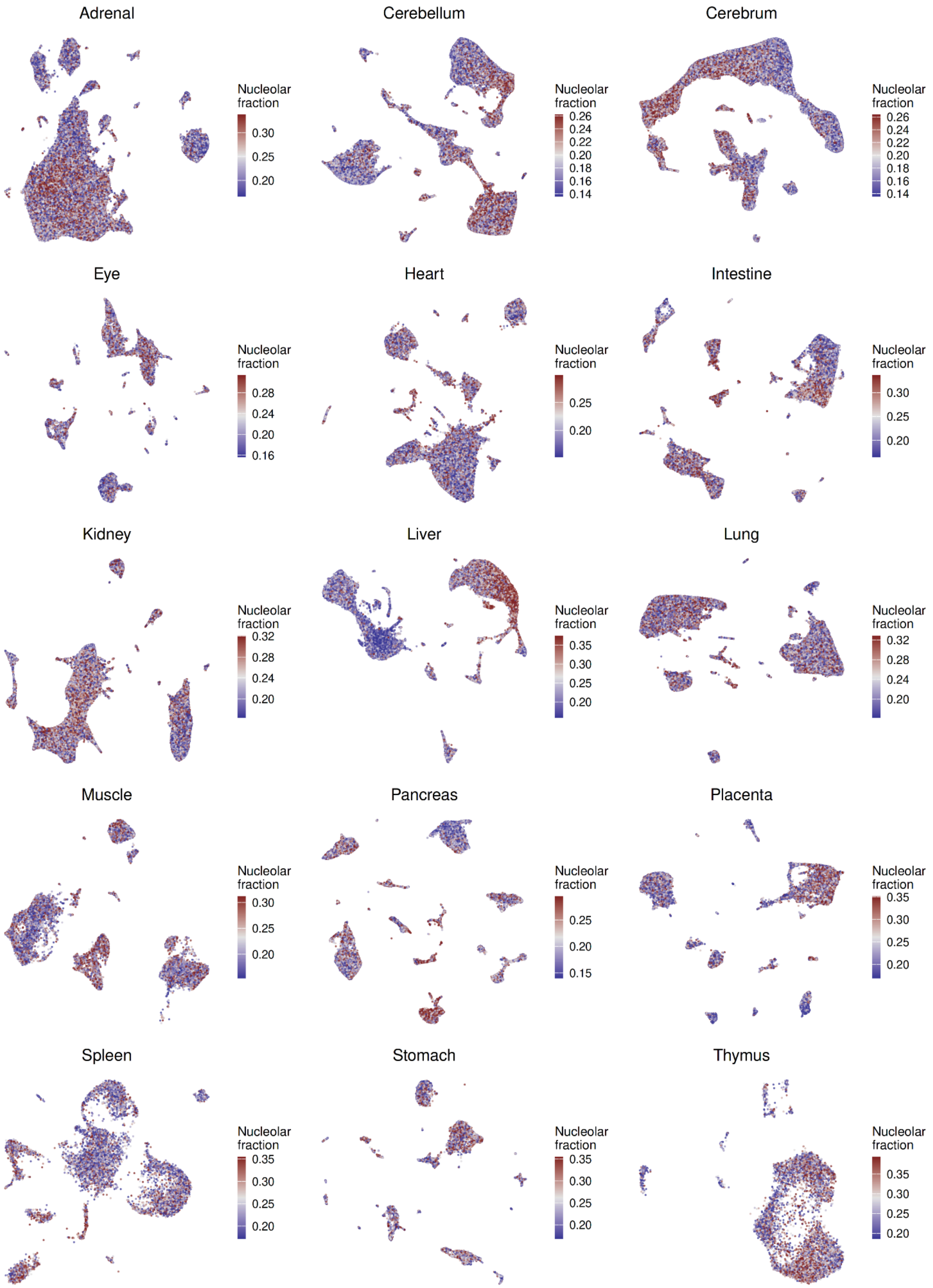


**Supplementary Figure 26. Mapping of the nucleolar fraction score across organs in the human fetal atlas.** UMAP embeddings of the human fetal atlas faceted by organ, with individual cells colored by the estimated nucleolar fraction score.

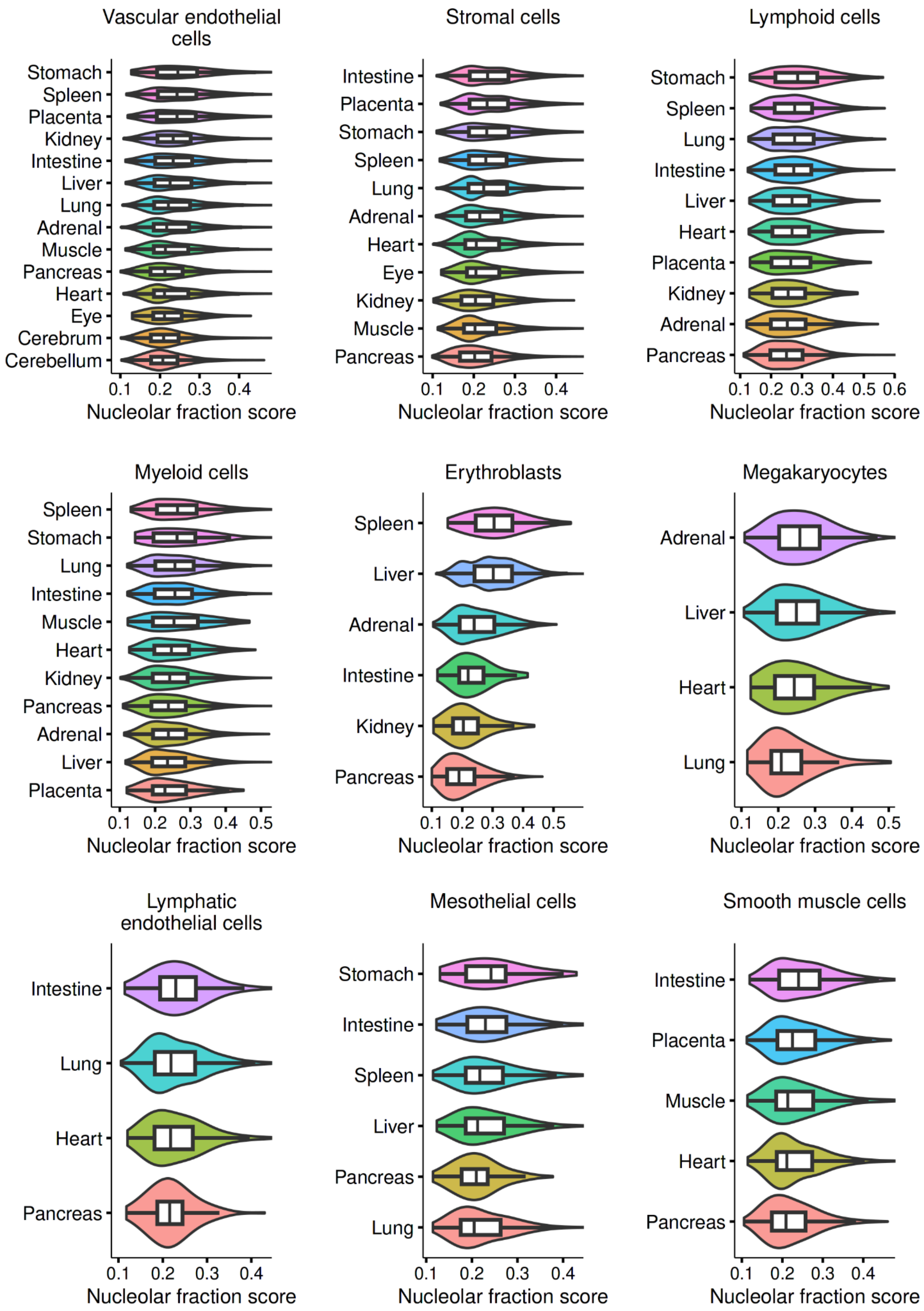


**Supplementary Figure 27. Comparison of the estimated nucleolar fraction score in cell types shared across organs.** Violin plots of the estimated nucleolar fraction score across organs for nine cell types common to multiple tissues within the human fetal atlas.

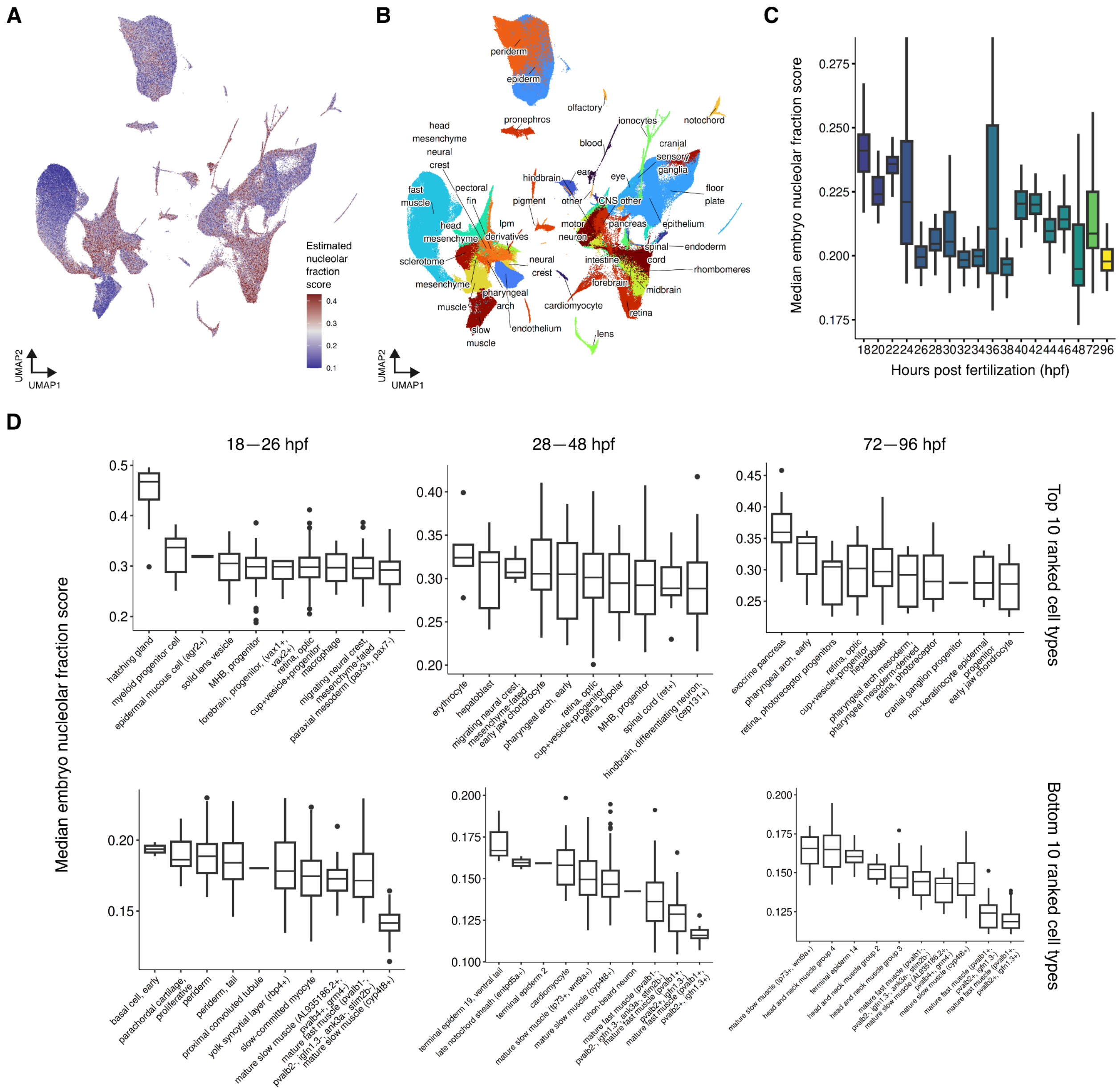

**Supplementary Figure 28. Estimation of the nucleolar fraction score on the zebrafish embryogenesis atlas.** (A, B) UMAP embeddings of the zebrafish embryogenesis atlas colored by (A) the estimated nucleolar fraction score and (B) major tissue type. (C) Boxplots of the median nucleolar fraction score at each timepoint, demonstrating a gradual decrease over developmental time. (D) Boxplots showing the score of the 10 highest and lowest ranked cell types, grouped by early (left; 18-26 hpf), mid (middle; 28-48 hpf) and late (right; 72-96 hpf) stages.
